# Mapping transmembrane binding partners for E-cadherin ectodomains

**DOI:** 10.1101/2020.05.08.084889

**Authors:** Omer Shafraz, Bin Xie, Soichiro Yamada, Sanjeevi Sivasankar

## Abstract

We combine proximity labeling and single molecule binding assays, to discover novel transmembrane protein interactions in cells. We first screen for candidate binding partners by tagging the extracellular and cytoplasmic regions of a bait protein with TurboID biotin ligase, and identify proximal proteins that are biotin-tagged on both their extracellular and intracellular regions. We then test direct binding interactions between the proximal proteins and the bait, using single molecule Atomic Force Microscope binding assays. Using this approach, we identify novel binding partners for the extracellular region of E-cadherin, an essential cell-cell adhesion protein. We show that the desmosomal proteins desmoglein-2 and desmocollin-3, the focal adhesion protein integrin-α2β1, and the receptor tyrosine kinase ligand ephrin-B1, all directly interact with E-cadherin ectodomains. Our discovery of previously unknown heterophilic E-cadherin binding interactions, suggest the existence of novel cadherin cross-talk in epithelial cells.

## INTRODUCTION

Transmembrane proteins play essential roles in coupling cells and in sensing mechanical and biochemical signals from the environment. However, it is extremely challenging to identify membrane protein interactions using traditional methods like affinity pulldown, because of the hydrophobic nature of membrane spanning regions, harsh extraction conditions that disrupt protein interactions, and inability of these methods to discriminate between cytoplasmic and extracellular protein interactions. Consequently, membrane protein interactomes have been mapped using several proximity-tagging systems such as BioID [1], APEX [2], and PUP-IT [3], where a protein of interest (the bait) is genetically fused to a proximity-based labeling enzyme which activates a substrate like biotin or Pup, and then releases the activated substrate to label proximal proteins. Recently, a micro-mapping platform using a photocatalytic induced carbene was also introduced for proximity labeling of membrane bound proteins [4]. While enormously powerful, these assays merely report on proteins that are proximal to the bait; testing direct interactions between proximate proteins and the bait require proximity-tagging techniques to be integrated with a complementary biophysical method.

Here we integrate the most widely used proximity-tagging scheme, BioID, with single molecule Atomic Force Microscope (AFM) binding assays, to identify transmembrane proteins that bind to the extracellular region of a transmembrane bait protein. In BioID, a promiscuous, mutant biotin ligase is fused to a bait protein and covalently tags nearby proteins with exogenously-supplied biotin [1, 5]; the biotin-tagged proteins are subsequently detected using mass spectrometry (MS). Unfortunately, the number of proximate proteins reported in a typical BioID screen are enormous (hundreds of candidate proteins), which limits the number of binding interactions that can be directly tested. We reasoned that transmembrane proteins that directly interact with a transmembrane bait would be positioned in close proximity to both the bait’s ectodomain and cytoplasmic regions. Consequently, if the proximity-based labeling enzyme was fused to both the bait’s extracellular and cytoplasmic regions, identifying transmembrane proteins that are biotinylated in both regions would enable us to narrow down the list of possible binding partners for subsequent AFM binding measurements. However, the BioID method has only been utilized to identify intracellular binding partners [6], including to the cytoplasmic region of cadherin cell adhesion proteins [7–9] and other junctional proteins [10]; it is unclear if BioID can successfully be used to screen for extracellular binding partners of transmembrane proteins.

Here, we demonstrate that BioID proximity tagging can be used to map ectodomain interactomes and that performing tandem BioID experiments on the extracellular and cytoplasmic region, narrows down potential transmembrane binding partners. Furthermore, we show that integrating BioID screens with single molecule AFM binding assays is extremely powerful in directly assaying candidate interactions. As a transmembrane bait, we use E-cadherin (Ecad), a ubiquitous cell-cell adhesion protein that plays an essential role in tissue morphogenesis, in maintaining tissue integrity and in facilitating the collective migration of cells [11, 12]. While over 170 proteins, including signaling molecules, scaffolding proteins and cytoskeletal regulators, have been reported to associate directly or indirectly with the Ecad cytoplasmic tail [13], Ecad ectodomains are primarily believed to interact with identical Ecad ectodomains from opposing cells. Although previous cell-free binding studies and microscopy suggest interactions between Ecad ectodomains and other adhesion proteins and cell-surface receptors [14–16], heterophilic Ecad binding partners have not been systematically catalogued.

We therefore fused BioID to the ectodomain and cytoplasmic regions of Ecad in epithelial cells. By comparing the extracellular and cytoplasmic interactomes, we identified just seven transmembrane proteins that are proximal to Ecad. Single molecule AFM binding assays revealed that out of these target proteins, the extracellular regions of the desmosomal proteins desmoglein-2 (Dsg2) and desmocollin-3 (Dsc3), the focal adhesion protein integrin-α2β1 (Intα2β1), and the receptor tyrosine kinase ligand ephrin-B1 (EfnB1), all directly interact with Ecad ectodomains. Other than a recent study showing direct interactions between Dsg2 and Ecad ectodomains [7], the other heterophilic interactions revealed by our measurements have not been previously identified. Our results demonstrate that Ecad ectodomains do not merely engage in homophilic binding, but instead, like the Ecad cytoplasmic region, also bind to a range of junctional proteins.

## RESULTS AND DISCUSSION

### Ecad tagged with BioID on their extracellular and cytoplasmic regions localize to cell-cell junctions and biotinylate proteins at intercellular contacts

Ecad ectodomains comprise of five tandemly arranged extracellular domains (EC1-5). In our experiments, we fused the biotin ligase TurboID (which labels proteins in 10 minutes compared to ~18 h for previously used ligases) [17] to either the EC2 extracellular domain (EC-BioID) or cytoplasmic region (C-BioID) of Ecad and expressed these fusion constructs in epithelial Madin-Darby Canine Kidney (MDCK) cells. Due to post translational cleavage of the N-terminal signal- and pro-peptide, inserting BioID at the Ecad N-terminus was not feasible. Furthermore, since the Ecad homophilic binding site is located on the protein’s N-terminal EC1 domain, modifications in this region abolish Ecad adhesion. We therefore generated the EC-BioID, by inserting TurboID on the EC2 domain, at amino acid 152 from the N-terminus (Fig.1a); this location was previously used to insert a Green Fluorescent Protein (GFP) without affecting Ecad function [18]. C-BioID was generated by fusing TurboID to the cytoplasmic C-terminus of Ecad (Fig. 1b). A GFP molecule was inserted at the C-terminus of both the EC-BioID and C-BioID constructs (Fig. 1a, b, S1 a, c). Because of high levels of endogenous Ecad in parental MDCK cells, we rescued Ecad knockout MDCK cells with EC-BioID and C-BioID and generated stable cell lines. Both EC-BioID and C-BioID localized to cell-cell junctions, verifying that the insertion of TurboID, did not disrupt the incorporation of Ecad into intercellular junctions (Fig. 1c, 1d, top row). Next, to demonstrate that the TurboID was functional, we incubated the cells with free biotin, and tagged the resulting biotinylated proteins with fluorescently labeled streptavidin (Fig. 1c, 1d, middle row). In the absence of exogenous biotin, we observed very little fluorescent streptavidin signal. Western blots (Fig. S1 b, d) also confirmed that the TurboID required an external source of biotin to efficiently label proteins. Merging GFP and streptavidin images (Fig. 1c, 1d, bottom row) showed that most of the biotinylated proteins were localized to intercellular junctions.

**Figure 1:**
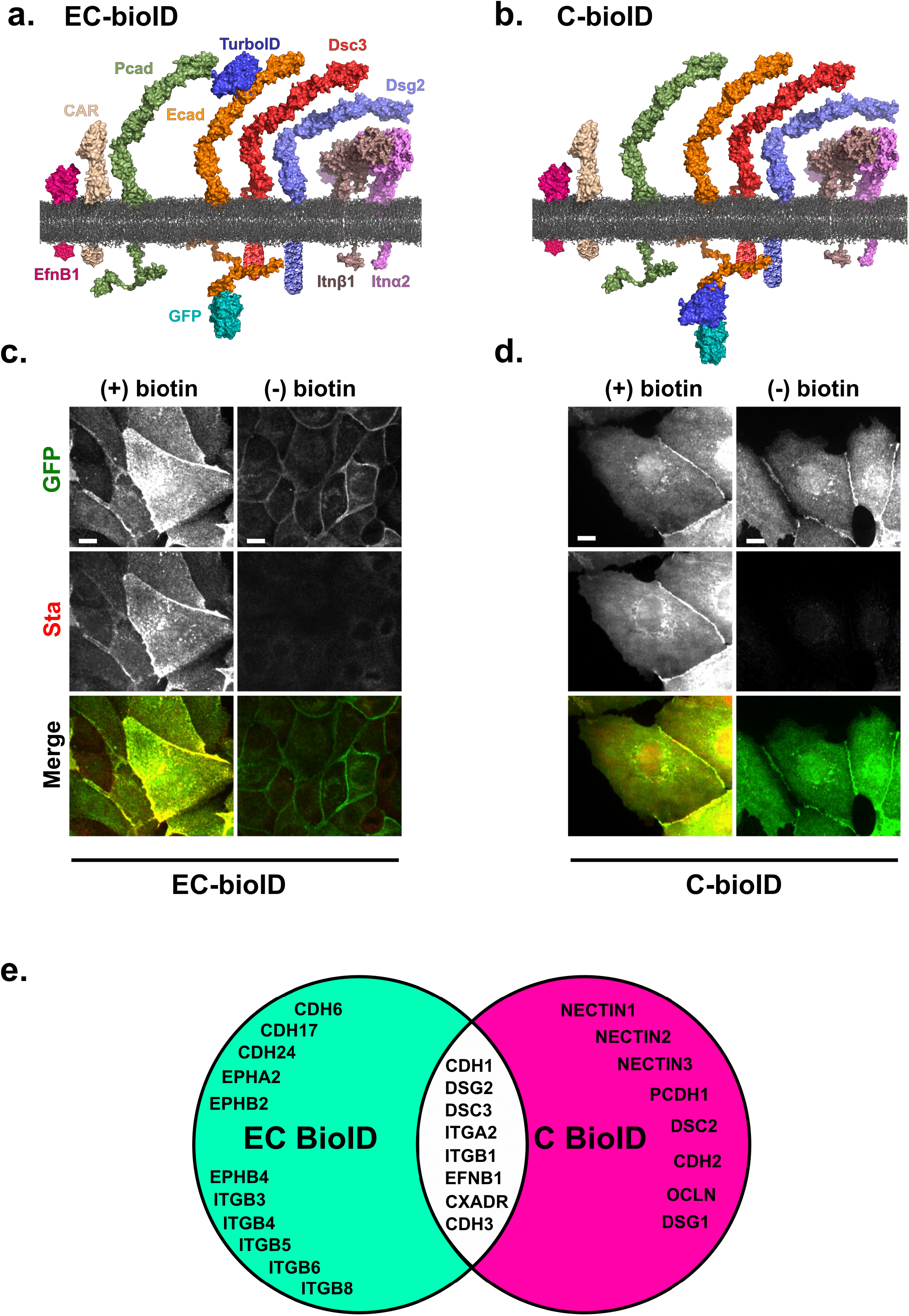
Characterizing the Ecad BioID constructs. TurboID was fused at **(a)** EC2 domain of Ecad (EC-BioID) and **(b)** C-terminal of Ecad (C-BioID). Proteomic analysis of both constructs identified ephrin-B1 (EfnB1), coxsackie and adenovirus receptor (CAR), P-cadherin (Pcad), desmocollin-3 (Dsc3), desmoglein-2 (Dsg2), Integrin β1 (Intβ1) and Integrin α2 (Intα2) as proteins that were proximal to Ecad. Proteins are depicted using deposited PDB structures or predicted from homology modelling using Modeller. The following PDB-IDs were used in generating the images: EfnB1: 6P7Y, CAR: 3J6N, TurboID: 4WF2, GFP: 1GFL, Ecad and Pcad: 3Q2V, Dsc3: 5IRY, Dsg2: 5ERD, Intα2: 3K6S, Intβ1: 3IJE, and membrane: POPE. Images were reconstructed using PyMOL. All cyto-plasmic domains are schematic representations. GFP localized to intercellular junction in **(c)** EC-BioID and **(d)** C-BioID confirming that Ecad is functional. Fluorescent streptavidin (Sta) staining showed that while BioID biotinylated proteins in the presence (+) of exogenous biotin, low levels of biotinylation was observed in the absence (-) of exogenous biotin. Merged images of GFP and Sta shows most of the biotinylation occur near the junction. **(e)** Venn diagram showing Ecad proximal transmembrane proteins identified from MS analysis of EC-BioID (green) and C-BioID (pink). Transmembrane candidate proteins observed in both BioID screens are listed in the overlapping region. Scale bar: 10 μm.

### Mass Spectrometry analysis identifies novel transmembrane proteins proximal to Ecad

To identify Ecad-proximate proteins, we incubated the EC-BioID and C-BioID cells with exogenous biotin and captured the biotinylated proteins from cell lysates using streptavidin coated beads. Trypsin digestion of the beads released protein fragments that were analyzed using MS. The proteomics analysis reported 300 proximal proteins in EC-BioID cells and 946 proximal proteins in C-BioID cells (MS data attached in SI), similar to previous published cytoplasmic interactomes of Ecad [7, 9]. However, since our primary focus was to identify extracellular binding partners, we selected transmembrane proteins found in all replicates with more than 4.8% peptide coverage (ratio between detected peptides and predicted peptides from trypsin digestion). This resulted in 19 proteins that were found in the EC-BioID (Fig. 1e, table S1) and 16 that were found in C-BioID (Fig. 1e, table S2). Of these ‘hits’, 7 proteins in addition to Ecad were common to both samples and consequently are possible binding partners for Ecad (Fig. 1e). These proteins (along with their gene IDs) are: Desmoglein-2 (DSG2), Desmocollin-3 (DSC3), Integrin alpha 2 (ITGA2), Integrin beta 1 (ITGB1), Ephrin-B1 (EFNB1), Coxsackie and adenovirus receptor (CXADR; CAR), and P-cadherin (CDH3; Pcad). Only few of these proteins were detected at a significant level in the absence of biotin (table S1, S2). It is important to note that our BioID experiments cannot distinguish if these proteins bound to Ecad in *trans* (from opposing cells) or *cis* (from the same cell) orientations since *trans* interacting proteins could still be in close *cis* proximity to Ecad.

### Single molecule AFM binding assays

Since BioID merely screens for protein proximity, we next probed the direct binding of ectodomains of 5 of the 7 candidate protein hits (Dsc3, Intα2-Intβ1 heterodimers, EfnB1, and CAR) with Ecad ectodomains, using single molecule AFM binding assays. We have already shown, using single molecule and cellular structure-function experiments, that the ectodomain of Dsg2 (the sixth protein hit in the BioID experiments) directly interacts with Ecad ectodomains [14]. Consequently, only Pcad-Ecad interactions remain untested. In our AFM experiments, we used the complete extracellular region of all the target proteins expressed and purified from mammalian cells. The C-terminus of the proteins were either tagged with human Fc-dimers (EfnB1 and CAR), biotin (Dsc3 and Ecad) or human c-Jun and c-Fos (Intα2β1 heterodimer). The biotin, c-Jun and Fc tags were used to immobilize the proteins on AFM tips and glass coverslips (Fig. 2a, 2c, 2e, 2g, 2i; Methods). Briefly, AFM cantilevers and coverslips were functionalized with a monolayer of Polyethylene Glycol (PEG) decorated with streptavidin. Biotinylated Ecad and Dsc3 were directly attached to the streptavidin (Fig. 2a, 2c), while heterodimeric Intα2β1 was linked to biotinylated protein G via a c-Jun antibody (2e). Fc tagged EfnB1 and CAR were linked to streptavidin using biotinylated protein G (Fig. 2g, 2i) (Methods).

**Figure 2:**
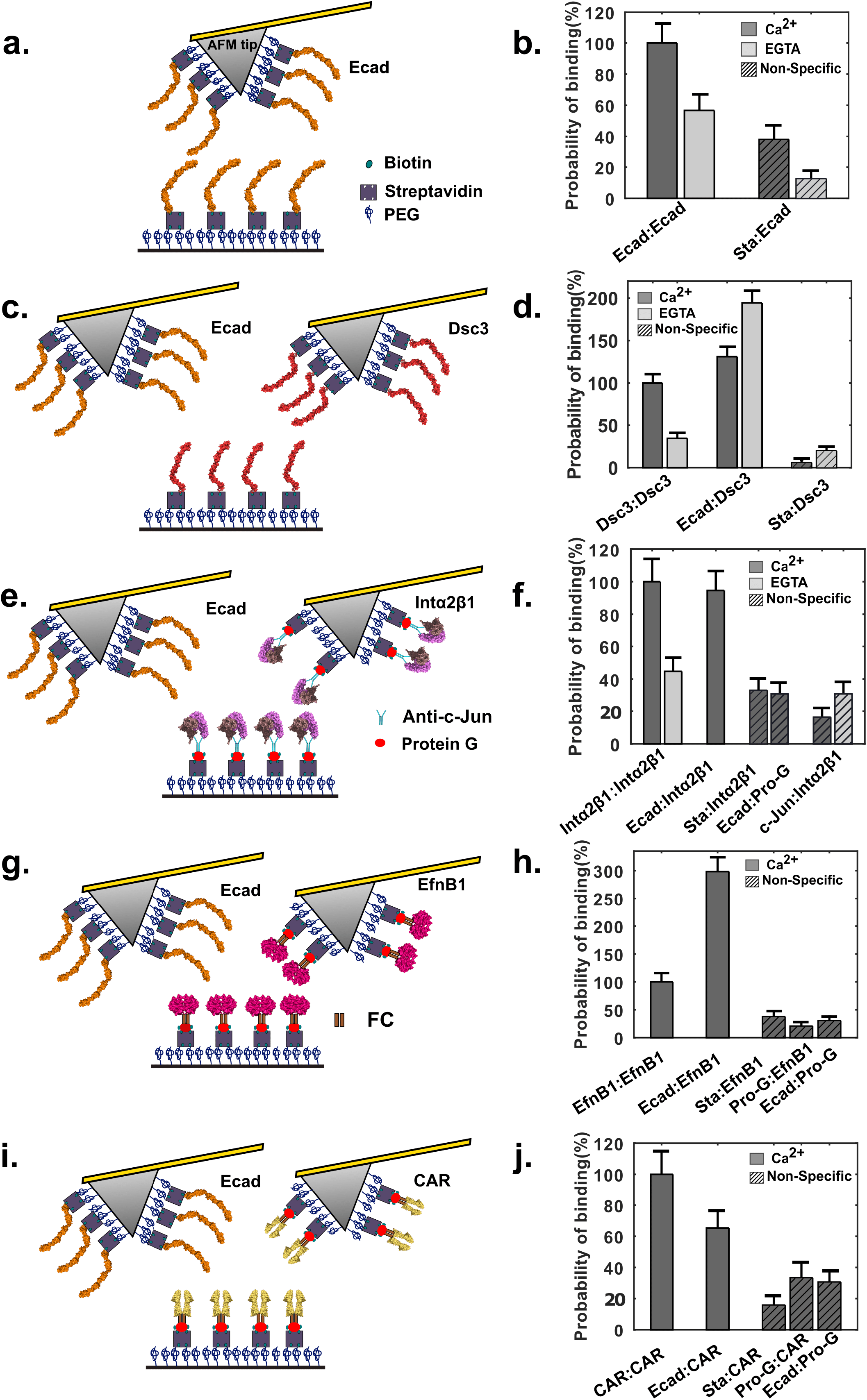
Single molecule AFM force measurements confirm novel binding partners for Ecad. **(a)** Schematic of single molecule AFM force measurement experiment. The AFM tip and substrate were functionalized with PEG linkers, some of which were decorated with streptavidins (Sta). Biotinylated Ecad was directly attached to Sta. **(b)** Ecad showed Ca^2+^ dependent homophilic interactions. **(c)** Homophilic binding probability was determined with biotinylated Dsc3 immobilized on both AFM tip and substrate. Heterophilic measurements were performed with Ecad on AFM tip and biotinylated Dsc3 on substrate. **(d)** Dsc3 showed Ca^2+^ dependent homophilic interactions and Ca^2+^ independent interactions with Ecad. **(e)** Intα2β1 heterodimer tagged with c-Jun was immobilized using an anti-c-Jun antibody and biotinylated protein G (Pro-G) **(f)** Intα2β1 formed Ca^2+^ dependent homophilic complexes and also formed heterophilic complexes with Ecad. **(g)** Fc-tagged EfnB1 was attached to Sta using biotinylated Pro-G **(h)** EfnB1 interacted homophilically and also bound heterophilic to Ecad. **(i)** CAR-Fc was immobilized using biotinylated Pro-G. **(j)** CAR showed low probability of heterophilic interactions with Ecad. Error bars are s.e. calculated using bootstrap with replacement.

At the start of each experiment, the AFM cantilever and substrate were brought into contact to allow opposing proteins to interact and the tip was then withdrawn from the substrate. As previously described [14], interaction of proteins resulted in unbinding events characterized by non-linear stretching of the PEG tethers which served as a molecular fingerprint for single molecule unbinding. We measured the homophilic interaction of every candidate protein, the heterophilic interaction of every candidate protein with Ecad, and nonspecific interactions under each measurement condition. Nonspecific interactions were determined using both an Ecad functionalized AFM tip and a coverslip lacking the candidate protein and also using an AFM tip lacking Ecad and a coverslip functionalized with the candidate proteins. For ease of comparison, all binding probabilities were normalized by the corresponding Ca^2+^-dependent homophilic binding rates. We confirmed that the Ecad is functional by measuring its homophilic binding properties (Fig. 2b). Since Ecad is a Ca^2+^ dependent adhesion protein, we confirmed that its homophilic binding probability was reduced in the presence of EGTA, a Ca^2+^ chelating agent (Fig. 2b, table S3).

#### Dsc3 dimerizes with Ecad independent of Ca^2+^

One of the most abundant proteins found in both EC-BioID and C-BioID were different isoforms of the desmosome associated proteins, Dsc and Dsg, which mediate robust cell-cell adhesion in tissues like the epidermis and heart that are exposed to significant levels of mechanical stress. While Dsg2 and Dsc3 were identified in both EC-BioID and C-BioID, Dsc2 and Dsg1 were identified in C-BioID. As predicted by previous biophysical studies [19], our AFM measurements showed that Dsc3 interacts homophilically in a Ca^2+^ dependent fashion (Fig. 2d, table S3). Surprisingly, Dsc3 ectodomains also interacted heterophillically with Ecad ectodomains in a Ca^2+^ independent fashion (Fig. 2d, table S3). It is important to point out that we have already demonstrated that Ecad and Dsg2 directly bind via a conserved Leu175 on the Ecad, which promotes desmosome assembly.

The interaction of Dsg2 and Dsc3 with Ecad provide a molecular explanation for previous studies showing that blocking Ecad adhesion with antibodies or knocking down Ecad, delay desmosome formation in cells [20, 21], and that classical cadherin deficient mice show defective desmosome assembly [22]. While we have shown that Dsc2 does not directly bind to Ecad [14], AFM and structural studies demonstrate that Dsg2 interacts with Dsc2 [14, 23]. Consequently, Dsc2 still appears as a ‘hit’ in our C-BioID assay. We also detect Dsg1 in C-BioID, likely because Dsg1 forms a heterotypic dimer with Dsc3 [23]. Intriguingly, although Dsg3 colocalizes with Ecad in keratinocytes [24–26], we do not measure it in the BioID assays.

#### Ecad is a ligand for Integrin α2β1

Interestingly, both EC-BioID and C-BioID showed that the integrin focal adhesion proteins, Intα2 and Intβ1, were proximate to Ecad ectodomain. Since integrins are composed of non-covalently associated αβ heterodimers and Intα2 forms a heterodimer exclusively with Intβ1, we used Intα2β1 in our AFM binding measurements. Similar to Intα3β1 that interacts homophilically in a cation dependent manner [27], our AFM measurements demonstrated homophilic Ca^2+^ dependent Intα2β1 adhesion (Fig. 2f, table S3). Furthermore, as anticipated from the BioID results, we measured heterophilic adhesion between Ecad ectodomains and Intα2β1 ectodomains in Ca^2+^ (Fig. 2f, table S3).

The discovery of interactions between Ecad and Intα2β1 ectodomains is particularly exciting since although there is a large body of evidence supporting the existence of crosstalk between integrins and cadherins [28–30], this crosstalk is primarily believed to be mediated by cytoplasmic proteins. Even though Intα2β1 is a receptor for the ECM proteins collagen and laminin [31], immunofluorescence studies have shown that Intα2β1 localizes to cell-cell contacts in keratinocytes [32] and with Ecad and N-cadherin in melanoma cells [33]. Furthermore, cell-spreading assays suggest that interaction between cells and an Ecad coated surface is abolished by monoclonal antibodies that target Intα2β1 [34]. Our demonstration of homophilic binding and heterotypic adhesion with Ecad suggest that Intα2β1 could play a functional role in cell-cell adhesion and in collective cell migration.

#### EfnB1 forms heterotypic complexes with Ecad

Next, we measured the homophilic and heterophilic binding of EfnB1, a ligand for Eph, the largest subfamily of receptor tyrosine kinases [35]. In agreement with previous structural reports [36], our measurements showed that EfnB1 dimerizes homophilically (Fig. 2h, table S3). Interestingly, the heterophilic binding between Ecad monomers and EfnB1-Fc dimers was almost 3 fold higher than EfnB1 homophilic binding (Fig. 2h, table S3).

While EfnB1 is known to interact with the tight junction protein claudin [37], direct binding of EfnB1 and Ecad has not been previously reported. It has been shown that EphB2 receptors interact with Ecad and the metalloproteinase ADAM10 at sites of adhesion and their activation induces shedding of Ecad by ADAM10 at interfaces with EfnB1-expressing cells, which suppress tumor progression in colorectal cancer [38, 39]. It is possible that Ecad ectodomains interact with EfnB1 and structurally hinder ADAM10 mediated Ecad-cleavage. Consequently, when EfnB1 dissociates from Ecad and binds to EphB2 the ADAM10 cleavage site on Ecad become accessible. Finally, we note that while our AFM force measurement shows that EfnB1 forms homodimers, previous analytical ultracentrifugation measurements suggest that EfnB1 is monomeric [40]. This discrepancy may arise from the higher, single molecule, sensitivity of our AFM binding measurements.

#### CAR does not interact directly with Ecad

The last candidate protein we tested using AFM force measurements were Fc-dimers of CAR, a transmembrane virus receptor that localizes to tight junctions. In agreement with structural and sedimentation analysis [41], our AFM data also showed homophilic CAR interactions (Fig. 2j, table S3). However, the heterophilic binding probability of CAR and Ecad was only ~30% greater than nonspecific binding (Fig. 2j, table S3). This suggests that CAR ectodomains either do not interact with Ecad or bind to Ecad with a low probability.

Previous studies show that while Ecad and CAR are not colocalized in stable cell-cell junctions, CAR localization is observed with internalized Ecad in vesicles [42]. This suggests that CAR can be biotinylated in internalized vesicles with both EC-BioID and C-BioID, even though it might not directly interact with Ecad on the cell surface. Another possible explanation for the low Ecad-CAR heterophilic binding measured in our AFM experiments is that CAR and Ecad interactions are primarily mediated by their respective cytoplasmic domains which would still result in detection in EC-BioID and C-BioID screens, but not in the single molecule AFM binding assay that only tests for ectodomain interactions. Nonetheless, the results we obtain with CAR clearly demonstrate the advantage of integrating a direct assay for protein interaction with proximity screening assays.

The final protein candidate from our BioID assay was Pcad, a type-I classical cadherin that shares high sequence homology with Ecad [43]. Previous biophysical measurements show that Ecad and Pcad form weak heterodimers in solution [43] and cells expressing similar levels of Ecad and Pcad form intermixed cells that fail to sort out [44, 45]. This suggests that Pcad can bind heterophilically to Ecad in cells, which supports the results of our BioID screens. However, direct binding between Ecad and Pcad, remains to be resolved at the single molecule level. Notwithstanding, the results of our integrated assay demonstrate that Ecad ectodomains do not merely engage in homophilic binding, but instead, like the Ecad cytoplasmic region, also bind to a range of junctional proteins. We anticipate that integrated BioID-AFM experiments can be similarly applied to identify heterophilic binding partners for other transmembrane proteins.

## Methods

### Molecular cloning of EC-BioID and C-BioID

We inserted TurboID in Ecad at a site where GFP had previously been inserted in mouse N-cadherin using *in vitro* transposition [18]. We identified from sequence alignment, a similar sequence at position 152 from N-terminal sequence in canine Ecad and inserted TurboID with a peptide linker on both ends; the linker was composed of the same amino acids (***AYSILT***-LSLIHIWRA-**V5-TurboID**-GRARADVYKRQ-***QDPLLP***; canine Ecad sequence is in bold italicized font while linker sequence is in normal font) used previously [18].

We PCR amplified Ecad using pEGFP-N1-Ecad plasmid [46] and V5-TurboID using mutant BirA R118S (TurboID) (Addgene [17]) using PCR primers (Table S4). The PCR products were inserted between EcoRV and BglII using gibson assembly (New England Biolabs) in pEGFP-N1-Ecad vector. We PCR amplified the EC-BioID sequence and inserted it into the PiggyBac-GFP vector using NheI site (PB533A, System Biosciences, Mountain View, CA). For C-BioID, we PCR amplified using PCR primers (Table S4) the complete Ecad using pEGFP-N1-Ecad plasmid. The PCR products were inserted at BamHI site in a PiggyBac vector.

The PiggyBac EC-BioID-GFP and C-BioID-EGFP plasmids were transfected into Ecad-KO cells [47] using Lipofectamine2000 (Invitrogen) along with PiggyBac Transposase expression plasmid (System Biosciences) and selected with 500 μg/ml G418 (Invitrogen). G418-resistant cells were sub-cloned and selected by confocal microscopy and western blotting to obtain homogeneous cell populations.

### Immuno-fluorescence

EC-BioID and C-BioID expressing MDCK cells were grown to confluency and switched to serum free DMEM cell culture media and supplemented with 50 μM biotin overnight. To selectively detect the extracellular biotinylation, AlexaFluor-568-conjugated streptavidin (Invitrogen) was added for 30 minutes before cell fixation and permeabilization. Cells were fixed using 3% paraformaldehyde and 0.3% Triton X-100 in PBS for 10 minutes and blocked with 1% BSA and 0.3% Triton X-100 in PBS for 30 minutes. Anti-GFP (rabbit polyclonal, Invitrogen) antibody and AlexaFluor-488-conjugated anti-rabbit antibody were used to detect GFP-tagged proteins. For C-BioID expressing cells, biotinylated proteins were detected by incubating the fixed cells for 30 mins with AlexaFluor-568-conjugated streptavidin. Cells were imaged using a Zeiss AxioObserver equipped with a Yokogawa CSU-10 spinning disk confocal system, 40× objective, 488- and 561-nm solid-state lasers, a Photometrics CoolSNAP HQ2 camera, and Slidebook software (Intelligent Imaging Innovations). Images were reconstructed using ImageJ.

### Purification of biotinylated proteins for MS analysis

Four replicates of EC-BioID and two replicates of C-BioID were used for MS analysis. Cells expressing EC-BioID and C-BioID were seeded on p150 dishes and cultured to 70%-90% confluency in Dulbecco’s Modified Eagle’s Medium (DMEM) (Invitrogen) supplemented with 10% fetal bovine serum (Gibco), penicillin (Invitrogen), streptomycin (Invitrogen) and 200 μg/ml G418 (Invitrogen). Cells were then incubated overnight with serum free media supplemented with 50 μM biotin. After three PBS washes, cells were scraped and centrifuged. Pelleted cells were re-suspended in lysis buffer [48, 49] (50 mM Tris pH 7.5, 150 mM NaCl, 0.4 % SDS, 1 % IGEPAL CA-630, 1.5 mM MgCl2, 1 mM EGTA) with 2 μL/ml of protease inhibitor cocktail (Sigma-Aldrich), and 1 μL/ml Benzonase (250 U/μL) (EMD-Millipore). Cell lysate was incubated for 30 min at 4 °C and then sonicated at 10%-30% duty ratio for 1 min, and centrifuged at 13000 rpm for 30 min at 4 °C. The supernatant was collected, and their concentrations were measured with a RC/DC protein assay kit (Bio-Rad). Supernatant (1 ml at ~4-5 mg/ml) was incubated with 50 μl of superparamagnetic Dynabeads Streptavidin C1 (Invitrogen) and rotated overnight at 4 °C. First, beads were washed with the lysis buffer and transferred to new tubes. The beads were washed again with the 2% SDS in 50 mM Tris HCl pH 7.4, then washed twice with lysis buffer. Next, the beads were washed three times with 100mM ammonium bicarbonate (NH_4_HCO_3_) and re-suspended in 30uL of digestion buffer containing 1 μg/μL of trypsin gold in 50 mM NH_4_HCO_3_ for overnight digestion at 37 °C. To stop the digestion, the reaction mixture was acidified with 1% trifluoroacetic acid (TFA). The resulting peptides were recovered from the beads using a magnet and then acidified with 0.5% TFA final concentration. The eluted tryptic peptides were dried in a vacuum centrifuge and re-constituted in 0.1% formic acid. Tryptic peptides were analyzed using nano-scale liquid chromatographic tandem mass spectrometry (nLC-MS/MS).

### Nano-scale liquid chromatography

Nano-scale liquid chromatography (nLC) was performed on an ultra-high pressure nano-flow Easy nLC system (Bruker Baltonics). Liquid chromatography was performed at 40 °C with a constant flow rate of 400 nL/min on an in-house packed reversed-phase column (25 cm x 75 μm i.d.) with a pulled emitter tip, packed with 1.9 μm C18-coated porous silica beads (Dr. Maisch, Ammerbuch-Entringen, Germany). Mobile phases A and B were water with 0.1% formic acid (v/v) and 80/20/0.1% ACN/water/formic acid (v/v/v), respectively. Peptides were separated using a 33 min gradient from 5% to 38% B within 25 min, followed by an increase to 95% B within 1 min, a 1 min washing step at 95% B, ending with 8 min of re-equilibration at 5% B.

### Mass Spectrometry

Mass Spectrometry (MS) was performed on a hybrid trapped ion mobility spectrometry-quadrupole time of flight mass spectrometer (timsTOF Pro, Bruker Daltonics, Bremen, Germany) with a modified nano-electrospray ion source (CaptiveSpray, Bruker Daltonics). In the experiments described here, the mass spectrometer was operated in PASEF mode. All experiments were acquired with a 100 ms ramp and 10 PASEF MS/MS scans per topN acquisition cycle. Low-abundance precursors with an intensity below a ‘target value’ were repeatedly scheduled for PASEF-MS/MS scans until the summed ion counts reached a target value of 20,000 a.u. MS and MS/MS spectra were recorded from m/z 100 to 1700. A polygon filter was applied to the m/z and ion mobility plane to select features most likely representing peptide precursors rather than singly charged background ions. The quadrupole isolation width was set to 2 Th for m/z under 700 and 3 Th for m/z larger than 700, and the collision energy was ramped stepwise as a function of increasing ion mobility: 52 eV for 0%–19% of the ramp time; 47 eV from 19%–38%; 42 eV from 38%–57%; 37 eV from 57%–76%; and 32 eV for the remainder.

### MS Data Analysis

Mass spectrometry raw files were processed with MsFragger [50]. For all searches, a protein sequence database of reviewed canine proteins (accessed 11/27/2019 from UniProt; 1886 entries including decoys and 115 common contaminant sequences) was used. Decoy sequences were generated and appended to the original database for MSFragger. A maximum of two missing cleavages were allowed, the required minimum peptide sequence length was 7 amino acids, and the peptide mass was limited to a maximum of 5000 Da. Carbamidomethylation of cysteine residues was set as a fixed modification, and methionine oxidation and acetylation of protein N termini as variable modifications. The initial maximum mass tolerances were 50 ppm for precursor and fragment ions. A reversed sequence library was generated/used to control the false discovery rate (FDR) at less than 1% for peptide spectrum matches and protein group identifications. Decoy database hits, proteins identified as potential contaminants, and proteins identified exclusively by one site modification were excluded from further analysis. Label-free protein quantification was performed with the IonQuant algorithm [50]. All other MsFragger parameters were kept at their default values.

### Cloning, purification and biotinylation of Ecad and Dsc3 monomers for single molecule AFM experiments

The extracellular region of the Dsc3 was PCR amplified from full length human Dsc3 purchased from DNASU plasmid repository (clone ID: HsCD00821324). For Dsc3, the following PCR primers were used: 5’-tgtggtggaattctgcagatatcATGGCTGCCGCTGGCCCC (forward) and gcaagctttcgctagcCTTTCCCAGAATCACGCCTGTGCTTCTG (reverse). Avi-tag-Tev-6x His (ATH) sequence was amplified using pcDNA3.1(+) Ecad-pATH [51] using forward primer 5’-aaaggctagcGAAAGCTTGCTTGGTGGC and reverse primer 5’-gttcgaagggccctctagactcgagGATTAGTGATGATGGTGATGG. The Dsc3 extracellular region and and ATH fragments were cloned into pcDNA3.1(+) Ecad-pATH between EcoRV and XhoI sites.

Cloning of Ecad-ATH plasmids has been described elsewhere [52]. The Ecad and Dsc3 plasmids were transiently transfected into HEK 293T cells using Polyethylenimine (PEI) (Polysciences, Inc.). Four days post transfection, the conditioned media was collected for protein purification. As described previously [14], protein was purified using the NGC Chromatography Systems (BioRad). In 4 °C, media containing cadherin was flowed through Ni-NTA agarose beads (Qiagen) packed on 1 ml Empty Bio-Scale Mini Cartridges (Biorad #7324660). The column was then washed with buffer at pH 7.5 (25mM HEPES, 5mM NaCl, 1mM CaCl_2_, pH 7.5). Proteins bound to Ni-NTA were biotinylated by incubating the beads with BirA enzyme (BirA500 kit; Avidity) for 1hr in 30 °C followed by an overnight incubation at 4 °C. Next, the column was washed with 25 mM HEPES, 500mM NaCl, 1mM CaCl_2_ with 20mM Imidazole at pH 7.5 and then eluted with the same buffer containing 250 mM imidazole. Following purification, the protein was exchanged to a Tris 10mM, NaCl 100mM, KCl 10mM, CaCl_2_ 2.5mM, buffer at pH 7.5.

### Single molecule AFM force measurements

Biotinylated Ecad, biotinylated Dsc3, Intα2β1-c-Jun/c-Fos, EfnB1-Fc, and CAR-Fc were immobilized on coverslips (CS) and AFM cantilevers (Olympus, model TR400PSA) using previously described method [51]. Briefly, the CS and cantilevers were cleaned with 25% H_2_O_2_:75% H2SO4 and washed with DI water. Then the CS was cleaned with 1 M KOH and washed with DI water. Both the CS and cantilevers were washed with acetone and functionalized using 2% (v/v) 3-aminopropyltriethoxysilane (Sigma) dissolved in acetone. Next, N-hydroxysuccinimide ester functionalized PEG spacers (MW 5000, Lysan Bio) were covalently attached to the silanized AFM tip and coverslip; 7% of the PEG spacers were decorated with biotin groups. Prior to a measurement, the functionalized AFM cantilever and coverslip were incubated overnight with BSA (1 mg/ml) to further reduce non-specific binding. The tip and surface were then incubated with 0.1 mg/ml streptavidin for 30 minutes and biotinylated cadherins were attached to the streptavidin. Recombinant rat EfnB1-FC (R&D catalog no: 1596-B1) and recombinant human CAR-FC (R&D catalog no: 3336-CX) were linked to streptavidin using biotinylated protein G as previously shown [52]. Human Integrin α2β1 heterodimer with human c-Jun and human c-Fos (R&D catalog no: 5698-A2) was linked to biotinylated protein G by c-Jun antibody (R&D catalog no: MAB2670). Finally, the surfaces were incubated with 0.02 mg/ml biotin for 10 minutes to block the free biotin binding sites on streptavidin.

Force measurements were performed using an Agilent 5500 AFM with a closed loop scanner. The spring constants of the cantilevers were measured using the thermal fluctuation method [53]. All the experiments were performed in a pH 7.5 buffer containing 10 mM Tris-HCl, 100 mM NaCl and 10 mM KCl with either 2.5 mM Ca^2+^ or 2 mM EGTA. For each experiment 923-1448 force curves were collected (table S3). Force curves with non-linear polymer stretching greater than the contour length of a single PEG molecule, was counted as an interaction.

## Acknowledgements

Research in SS lab was supported in part by the National Institute of General Medical Sciences of the National Institutes of Health (R01GM121885). SY’s research was supported in part by NIH R03 EB021636 and NSF 1562095. We thank Dr. Gabriela Grigorean for performing LC-MS/MS and data analysis in Proteomics Core Facility of the Genome Center at University of California, Davis.

