## Supplemental Information for "Mapping transmembrane binding partners for E-cadherin ectodomains"

### Figure S1

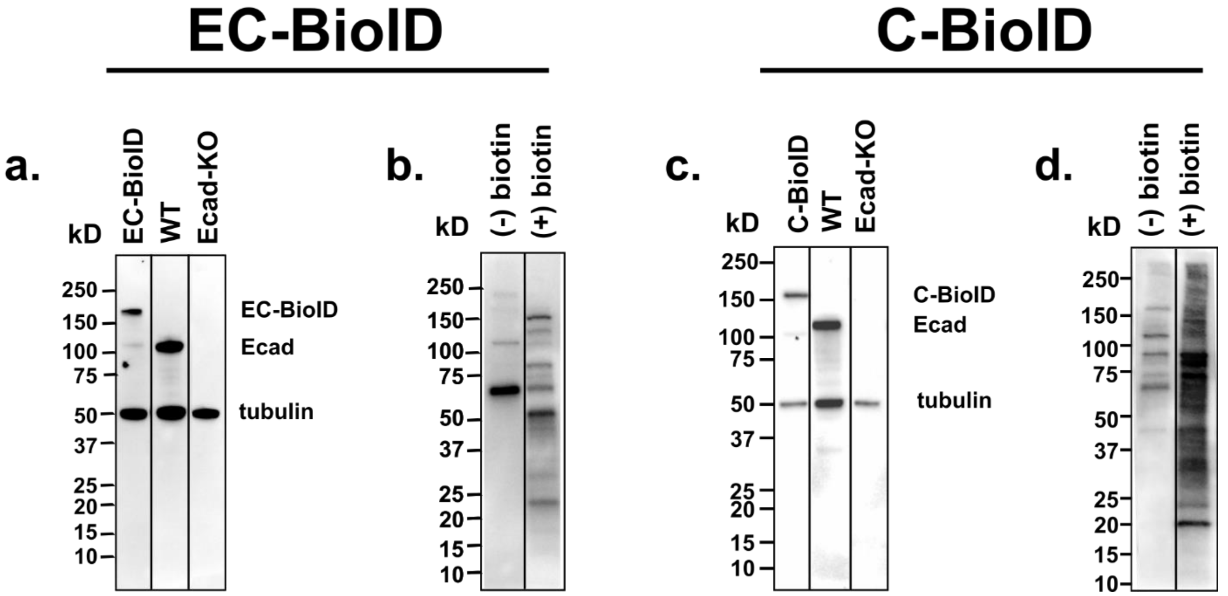

**Figure S1: Western blots a.** EC-BioID, WT and Ecad-KO cell lysates stained for Ecad and tubulin. **b.** HRP-streptavidin staining of biotinylated proteins eluted from streptavidin coated magnetic beads incubated with cell lysates of EC-BioID with (+) and without (-) exogenous biotin. **c.** C-BioID, WT and Ecad-KO cell lysates stained for Ecad and tubulin. **d.** HRP-streptavidin staining of biotinylated proteins eluted from streptavidin coated magnetic beads incubated with cell lysates of C-BioID with (+) and without (-) exogenous biotin.

| | (+) Biotin | | | | (-) Biotin | | | |
|---|---|---|---|---|---|---|---|---|
| | **Sample 1** | **Sample 2** | **Sample 3** | **Sample 4** | **Sample 1** | **Sample 2** | **Sample 3** | **Sample 4** |
| **Gene ID** | **Percent Coverage** | **Percent Coverage** | **Percent Coverage** | **Percent Coverage** | **Percent Coverage** | **Percent Coverage** | **Percent Coverage** | **Percent Coverage** |
| **CDH1** | 29.6 | 31.4 | 41.1 | 36.5 | 10.8 | 6.7 | 28.8 | 29.1 |
| **DSG2** | 26 | 14.6 | 45 | 37 | 0.8 | 1.9 | 1.6 | 18.7 |
| **CXADR** | 30.2 | 26.2 | 32.7 | 27.1 | 0.0 | 0.0 | 0.0 | 6.9 |
| **EFNB1** | 24.3 | 30.6 | 24 | 30.3 | 0.0 | 0.0 | 0.0 | 0.0 |
| **ITGA2** | 16.5 | 22.2 | 30.1 | 33.4 | 1.1 | 1.1 | 5.2 | 7.2 |
| **CDH3** | 21.8 | 9.7 | 20.6 | 25.3 | 1.3 | 1.3 | 0.0 | 0.0 |
| **ITGB1** | 11.8 | 16.7 | 23.9 | 20.3 | 0.0 | 2.9 | 8.5 | 5.8 |
| **DSC3** | 9.7 | 7.5 | 11.5 | 13.3 | 0.0 | 0.0 | 2.6 | 0.0 |
| EPHA2 | 23.2 | 31.6 | 31.6 | 30.5 | 0.8 | 0.0 | 0.0 | 5.7 |
| ITGB4 | 21.8 | 27.8 | 33.1 | 30.7 | 0.0 | 1.2 | 3.9 | 4.4 |
| ITGB3 | 23.5 | 22.2 | 26.8 | 24.7 | 0.0 | 0.0 | 5.2 | 9.1 |
| CDH6 | 22.8 | 18.1 | 28.6 | 24.3 | 0.0 | 0.0 | 0.0 | 9.1 |
| CDH17 | 8.8 | 12.4 | 20.7 | 18.4 | 0.0 | 0.0 | 0.0 | 0.0 |
| ITGB6 | 12.7 | 10.4 | 14 | 17.1 | 0.0 | 0.0 | 0.0 | 1.7 |
| EPHB4 | 11.4 | 8.1 | 14.2 | 16.3 | 0.0 | 0.0 | 0.0 | 0.0 |
| ITGB8 | 5 | 10 | 15 | 17.6 | 0.0 | 0.0 | 0.0 | 0.0 |
| ITGB5 | 6.2 | 9.5 | 15.2 | 13.8 | 0.0 | 0.0 | 0.0 | 0.0 |
| EPHB2 | 8.5 | 4.8 | 9.8 | 12.1 | 0.0 | 0.0 | 0.0 | 0.0 |
| CDH24 | 5.9 | 7.2 | 8.3 | 9 | 0.0 | 0.0 | 0.0 | 0.0 |

**Table S1: EC-BioID transmembrane protein hits.** Hits that also occur in C-BioID are in bold.

| | (+) Biotin | | (-) Biotin | | |
|---|---|---|---|---|---|
| | **Sample 1** | **Sample 2** | **Sample 1** | **Sample 2** | |
| **Gene ID** | **Percent coverage** | **Percent coverage** | **Percent Coverage** | **Percent Coverage** | **Previously reported in BioID** |
| **DSG2** | 54.2 | 61.1 | 14.3 | 35.3 | Guo et al.[1], Van Itallie et al [2] |
| **CXADR** | 53.4 | 55.3 | 0.0 | 24.9 | Guo et al., Van Itallie et al |
| **CDH1** | 30.7 | 27.2 | 24.3 | 22.7 | Guo et al., Van Itallie et al |
| **DSC3** | 22.7 | 23.9 | 4.2 | 9.0 | Guo et al., Van Itallie et al |
| **ITGA2** | 18.2 | 18.2 | 0.0 | 0.0 | |
| **EFNB1** | 18 | 18 | 0.0 | 0.0 | Van Itallie et al |
| **CDH3** | 7 | 12.4 | 0.0 | 0.0 | |
| **ITGB1** | 11.9 | 10.2 | 0.0 | 0.0 | Guo et al. |
| PCDH1 | 17 | 19 | 0.0 | 2.5 | Guo et al. |
| NECTIN2 | 44.5 | 38.1 | 0.0 | 0.0 | Guo et al., Van Itallie et al |
| DSC2 | 16.9 | 19.7 | 0.0 | 0.0 | Van Itallie et al |
| NECTIN3 | 14 | 18.4 | 0.0 | 0.0 | |
| CDH2 | 7.2 | 10.8 | 0.0 | 0.0 | |
| OCLN | 21.7 | 19.2 | 0.0 | 0.0 | Guo et al., Van Itallie et al |
| NECTIN1 | 6 | 7.2 | 0.0 | 0.0 | Guo et al. |
| DSG1 | 8.7 | 4.8 | 11.6 | 12.7 | |

**Table S2: C-BioID transmembrane protein hits.** Hits that also occur in EC-BioID are in bold.

| Experiment | Binding probability (%) | Total number of measured force curves |
|---|---|---|
| Ecad:Ecad (Ca $^{2+}$) | 100.0 ± 12.8 | 1297 |
| Ecad:Ecad (EGTA) | 56.7 ± 10.3 | 1134 |
| Ecad:Pro-G (Ca $^{2+}$) | 30.6 ± 7.2 | 1305 |
| Sta:Ecad (Ca $^{2+}$) | 38.0 ± 9.1 | 997 |
| Sta:Ecad (EGTA) | 12.6 ± 5.2 | 1060 |
| Dsc3:Dsc3 (Ca $^{2+}$) | 100.0 ± 10.4 | 1174 |
| Dsc3:Dsc3 (EGTA) | 34.5 ± 6.3 | 1225 |
| Ecad:Dsc3 (Ca $^{2+}$) | 131.1 ± 11.6 | 1224 |
| Ecad:Dsc3 (EGTA) | 194.4 ± 14 | 1224 |
| Sta:Dsc3 (Ca $^{2+}$) | 6.0 ± 2.7 | 1176 |
| Sta:Dsc3 (EGTA) | 19.9 ± 4.7 | 1274 |
| IntA2B1:IntA2B1 (Ca $^{2+}$) | 100.0 ± 14 | 961 |
| IntA2B1:IntA2B1 (EGTA) | 44.7 ± 8.4 | 1225 |
| Ecad:IntA2B1 (Ca $^{2+}$) | 94.6 ± 11.9 | 1265 |
| Sta:IntA2B1 (Ca $^{2+}$) | 32.8 ± 7.6 | 1076 |
| c-Jun:IntA2B1 (Ca $^{2+}$) | 16.3 ± 5.7 | 961 |
| c-Jun:IntA2B1 (EGTA) | 30.6 ± 7.7 | 960 |
| EfnB1:EfnB1 (Ca $^{2+}$) | 100.0 ± 15.7 | 1200 |
| Ecad :EfrnB1 (Ca $^{2+}$) | 298.3 ± 25.7 | 1310 |
| Sta:EfrnB1 (Ca $^{2+}$) | 37.6 ± 9.9 | 1162 |
| Pro-G:EfrnB1 (Ca $^{2+}$) | 20.6 ± 7.3 | 1213 |
| CAR:CAR (Ca $^{2+}$) | 100.0 ± 14.9 | 1206 |
| Ecad:CAR (Ca $^{2+}$) | 65.3 ± 11.1 | 1448 |
| Sta:CAR (Ca $^{2+}$) | 15.8 ± 6.0 | 1227 |
| Pro-G:CAR (Ca $^{2+}$) | 33.3 ± 10.0 | 923 |

**Table S3: Measured binding probability for single molecule force measurements**

| EC-BioID primers | | |
|---|---|---|
| Ecad 1-308 | Forward (F) | 5’-tgtgctggaattctgcagatATCCATCACACTGGCGGC |
| | Reverse (R) | 5’- tgtggatcaggctcagTGTGAGGATGCTGTAAGCG |
| V5-TurboID | F | 5’-catcctcacactgagcctgatccacatctggcgggccGGCAAGCCCATCCCCAAC |
| | R | 5’- cccgggcccggccCTTTTCGGCAGACCGCAG |
| Ecad 309-885 | F | 5’-tgccgaaaagggccgggcccgggccgacgtgtacaagcggcagATCCTCACACAAGACCCC |
| | R | 5’ -gggagatgtgttggggggaagatcTGGATCTGGATCAATGATGTTG |
| **C-BioID primers** | | |
| Ecad | F | 5’- gcgaattcgaatttaaatcgATGGGCCCTCGGTACGGC |
| | R | 5’- GTCGTCCTCGCCACCTCC |
| V5-TurboID | F | 5’- atggaggtggcgaggacgacGGCAAGCCCATCCCCAAC |
| | R | 5’- tgctcaccatCTTTTCGGCAGACCGCAG |
| EGFP | F | 5’- tgccgaaaagATGGTGAGCAAGGGCGAG |
| | R | 5’ - gggagaggggcgcggccgcgTTACTTGTACAGCTCGTCCATG |

**Table S4: List of PCR primers used in EC-BioID and C-BioID**

**Methods:**

**Western blot:** Ecad expression in lysates of WT, Ecad-KO, EC-BioID and C-BioID cells grown on 6-well plates were tested using western blotting. Anti-Ecad (BD Biosciences) and anti- tubulin antibodies (Cell Signaling Technology) were used with anti-mouse antibody conjugated with Horse Radish Peroxidase (HRP; Invitrogen). To test for biotinylation, an identical method, as described in main text, was used for cell lysate preparation and bead incubation. After the wash step, proteins bound to beads were eluted with 25 mM biotin at 95 °C and used for western blot analysis. HRP conjugated streptavidin (Invitrogen) was used for staining of biotinylated proteins. For HRP detection WesternBright Quantum Chemiluminescence Kit (Advansta) was used.

**Complete proteomic list obtained from MS analysis of EC-BioID**

**Bold**- Common hits in both EC-BioID and C-BioID; *Italic*- Transmembrane proteins in C-BioID only

| EC- BioID | | | | | | |
|---|---|---|---|---|---|---|
| | | | Sample 1 | Sample 2 | Sample 3 | Sample 4 |
| Gene ID | Organism | Protein Description | Percent Coverage | Percent Coverage | Percent Coverage | Percent Coverage |
| **CDH1** | Canis lupus familiaris OX=9615 | Cadherin 1 | 29.6 | 31.4 | 41.1 | 36.5 |
| **DSG2** | Canis lupus familiaris OX=9615 | Desmoglein 2 | 26 | 14.6 | 45 | 37 |
| **CXADR** | Canis lupus familiaris OX=9615 | Coxsackievirus and adenovirus receptor | 30.2 | 26.2 | 32.7 | 27.1 |
| **EFNB1** | Canis lupus familiaris OX=9615 | Ephrin B1 | 24.3 | 30.6 | 24 | 30.3 |
| **ITGA2** | Canis lupus familiaris OX=9615 | Integrin subunit alpha 2 | 16.5 | 22.2 | 30.1 | 33.4 |
| **CDH3** | Canis lupus familiaris OX=9615 | Cadherin 3 | 21.8 | 9.7 | 20.6 | 25.3 |
| **ITGB1** | Canis lupus familiaris OX=9615 | Integrin beta 1 | 11.8 | 16.7 | 23.9 | 20.3 |
| **DSC3** | Canis lupus familiaris OX=9615 | Desmocollin 3 | 9.7 | 7.5 | 11.5 | 13.3 |
| *EPHA2* | Canis lupus familiaris OX=9615 | EPH receptor A2 | 23.2 | 31.6 | 31.6 | 30.5 |
| *ITGB4* | Canis lupus familiaris OX=9615 | Integrin beta 4 | 21.8 | 27.8 | 33.1 | 30.7 |
| *ITGB3* | Canis lupus familiaris OX=9615 | Integrin beta 3 | 23.5 | 22.2 | 26.8 | 24.7 |
| *CDH6* | Canis lupus familiaris OX=9615 | Cadherin 6 | 22.8 | 18.1 | 28.6 | 24.3 |
| *CDH17* | Canis lupus familiaris OX=9615 | Cadherin 17 | 8.8 | 12.4 | 20.7 | 18.4 |
| *ITGB6* | Canis lupus familiaris OX=9615 | Integrin beta 6 | 12.7 | 10.4 | 14 | 17.1 |
| *EPHB4* | Canis lupus familiaris OX=9615 | EPH receptor B4 | 11.4 | 8.1 | 14.2 | 16.3 |
| *ITGB8* | Canis lupus familiaris OX=9615 | Integrin beta 8 | 5 | 10 | 15 | 17.6 |
| *ITGB5* | Canis lupus familiaris OX=9615 | Integrin beta 5 | 6.2 | 9.5 | 15.2 | 13.8 |
| *EPHB2* | Canis lupus familiaris OX=9615 | EPH receptor B2 | 8.5 | 4.8 | 9.8 | 12.1 |
| *CDH24* | Canis lupus familiaris OX=9615 | Cadherin 24 | 5.9 | 7.2 | 8.3 | 9 |
| NENF | Canis lupus familiaris OX=9615 | Neudesin neurotrophic factor | 70.7 | 70.7 | 69.8 | 69 |
| HSP90B1 | Canis lupus familiaris OX=9615 | Endoplasmin | 64.8 | 68.4 | 72.1 | 66.1 |
| PDIA4 | Canis lupus familiaris OX=9615 | Protein disulfide-isomerase A4 | 66.2 | 61.7 | 70.9 | 65.4 |
| HYOU1 | Canis lupus familiaris OX=9615 | Hypoxia up-regulated 1 | 60.9 | 60.8 | 58.3 | 61.8 |
| NUCB2 | Canis lupus familiaris OX=9615 | Nucleobindin 2 | 60.4 | 68.3 | 66.8 | 61.3 |
| SERPINH1 | Canis lupus familiaris OX=9615 | Serpin family H member 1 | 40.4 | 53.3 | 58.4 | 61 |
| P4HB | Canis lupus familiaris OX=9615 | Protein disulfide-isomerase | 60.4 | 67.1 | 68.2 | 60.6 |
| CKAP4 | Canis lupus familiaris OX=9615 | Cytoskeleton associated protein 4 | 54.7 | 49.5 | 57.2 | 60.1 |
| PC | Canis lupus familiaris OX=9615 | Pyruvate carboxylase | 55.3 | 62.6 | 61 | 59.5 |
| NUCB1 | Canis lupus familiaris OX=9615 | Nucleobindin 1 | 54.9 | 58.4 | 53 | 58.6 |
| VIM | Canis lupus familiaris OX=9615 | Vimentin | 19.1 | 16.3 | 54.7 | 57.5 |
| CALU | Canis lupus familiaris OX=9615 | Calumenin | 62.9 | 73.3 | 58.7 | 57.5 |
| PRXL2A | Canis lupus familiaris OX=9615 | Peroxiredoxin like 2A | 26.9 | 38 | 57.4 | 56.5 |
| HSPA5 | Canis lupus familiaris OX=9615 | Heat shock protein family A (Hsp70) member 5 | 61.3 | 61.2 | 56.4 | 55.2 |
| PDIA3 | Canis lupus familiaris OX=9615 | Protein disulfide-isomerase | 55 | 50.3 | 56.4 | 54.5 |
| DNAJC3 | Canis lupus familiaris OX=9615 | Uncharacterized protein | 57.2 | 59.4 | 58.4 | 54.1 |
| LRPAP1 | Canis lupus familiaris OX=9615 | LDL receptor related protein associated protein 1 | 56 | 59.9 | 54.1 | 53.3 |
| PPIB | Canis lupus familiaris OX=9615 | Peptidyl-prolyl cis-trans isomerase | 51.9 | 56.5 | 53.7 | 53.2 |
| RCN1 | Canis lupus familiaris OX=9615 | Reticulocalbin 1 | 28.9 | 68.7 | 62.2 | 51.6 |
| FAM172A | Canis lupus familiaris OX=9615 | Family with sequence similarity 172 member A | 40.2 | 49.6 | 48.8 | 51.5 |
| OS9 | Canis lupus familiaris OX=9615 | OS9 endoplasmic reticulum lectin | 54 | 46.1 | 53.6 | 51.2 |
| GOLM1 | Canis lupus familiaris OX=9615 | Golgi membrane protein 1 | 65.8 | 58.4 | 52 | 51 |
| PCCA | Canis lupus familiaris OX=9615 | Propionyl-CoA carboxylase subunit alpha | 53.1 | 56 | 54.6 | 48.8 |
| SPP1 | Canis lupus familiaris OX=9615 | Secreted phosphoprotein 1 | 50.8 | 45.8 | 46.2 | 47.8 |
| INHBA | Canis lupus familiaris OX=9615 | Inhibin subunit beta A | 18.6 | 22.2 | 42.5 | 45.8 |
| PDIA6 | Canis lupus familiaris OX=9615 | Uncharacterized protein | 48.4 | 49.8 | 44.5 | 45.7 |
| UGGT1 | Canis lupus familiaris OX=9615 | UDP-glucose glycoprotein glucosyltransferase 1 | 63.9 | 64.8 | 46.7 | 45.7 |
| KRT19 | Canis lupus familiaris OX=9615 | Keratin 19 | 19 | 24.6 | 52.9 | 45.4 |
| ANXA2 | Canis lupus familiaris OX=9615 | Annexin A2 | 24.8 | 28 | 38.6 | 45.4 |
| ACACA | Canis lupus familiaris OX=9615 | Acetyl-CoA carboxylase alpha | 40.8 | 48.2 | 42.7 | 45.4 |
| SIL1 | Canis lupus familiaris OX=9615 | SIL1 nucleotide exchange factor | 23.3 | 38.5 | 41 | 45 |
| PDIA5 | Canis lupus familiaris OX=9615 | Protein disulfide isomerase family A member 5 | 33.1 | 41.6 | 51.4 | 44.9 |
| TXNDC12 | Canis lupus familiaris OX=9615 | Thioredoxin domain containing 12 | 25.6 | 50 | 47.7 | 44.8 |
| NOP56 | Canis lupus familiaris OX=9615 | NOP56 ribonucleoprotein | 34.5 | 40.2 | 43.2 | 44.4 |
| MCCC1 | Canis lupus familiaris OX=9615 | Methylcrotonoyl-CoA carboxylase 1 | 38.3 | 24.1 | 37.4 | 43.9 |
| KRT14 | Canis lupus familiaris OX=9615 | Keratin 14 | 41.1 | 44.6 | 42.9 | 43.8 |
| HS3ST1 | Canis lupus familiaris OX=9615 | Sulfotransferase | 26.2 | 28.8 | 39.2 | 43.4 |
| PCMT1 | Canis lupus familiaris OX=9615 | Protein-L-isoaspartate O-methyltransferase | 22 | 43.4 | 46.2 | 43.4 |
| CNPY4 | Canis lupus familiaris OX=9615 | Canopy FGF signaling regulator 4 | 53.2 | 52 | 38.8 | 43.2 |
| GANAB | Canis lupus familiaris OX=9615 | Glucosidase II alpha subunit | 29.9 | 41.3 | 43.3 | 42.6 |
| P4HA2 | Canis lupus familiaris OX=9615 | Prolyl 4-hydroxylase subunit alpha 2 | 35.1 | 31.1 | 39.2 | 42 |
| TMEM43 | Canis lupus familiaris OX=9615 | Uncharacterized protein | 20 | 31.5 | 38.8 | 41.8 |
| RPN1 | Canis lupus familiaris OX=9615 | Dolichyl-diphosphooligosaccharide--protein glycosyltransferase subunit 1 | 27.2 | 37.1 | 40.5 | 40.9 |
| PCYOX1 | Canis lupus familiaris OX=9615 | Prenylcysteine oxidase 1 | 26.5 | 36.8 | 37.6 | 40.8 |
| C1GALT1 | Canis lupus familiaris OX=9615 | Glycoprotein-N-acetylgalactosamine 3-beta-galactosyltransferase 1 | 17.5 | 20.6 | 34.8 | 40.7 |
| APMAP | Canis lupus familiaris OX=9615 | Adipocyte plasma membrane associated protein | 15.2 | 22.7 | 36.9 | 40.7 |
| P4HA1 | Canis lupus familiaris OX=9615 | Prolyl 4-hydroxylase subunit alpha 1 | 30.9 | 35.4 | 37.6 | 40.6 |
| LMNA | Canis lupus familiaris OX=9615 | Lamin A/C | 14.9 | 14.3 | 38.2 | 40.3 |
| CTSD | Canis lupus familiaris OX=9615 | Cathepsin D | 33.7 | 36.6 | 35.6 | 40.2 |
| MYH9 | Canis lupus familiaris OX=9615 | Myosin-9 | 7.7 | 7.8 | 22.4 | 40.1 |
| ASAH1 | Canis lupus familiaris OX=9615 | N-acylsphingosine amidohydrolase 1 | 30.1 | 36.6 | 44.7 | 39.6 |
| HSP90AB1 | Canis lupus familiaris OX=9615 | HATPase_c domain-containing protein | 10.5 | 17.5 | 33.8 | 39.4 |

| | | | | | | |
|---|---|---|---|---|---|---|
| TSPAN4 | Canis lupus familiaris OX=9615 | Chitinase domain containing 1 | 16.5 | 36.6 | 33.1 | 39.4 |
| TTC13 | Canis lupus familiaris OX=9615 | Tetratricopeptide repeat domain 13 | 20.1 | 26.3 | 40.9 | 39.1 |
| FUCA2 | Canis lupus familiaris OX=9615 | Alpha-L-fucosidase | 49.1 | 47.4 | 39 | 39 |
| SDF4 | Canis lupus familiaris OX=9615 | Stromal cell derived factor 4 | 36.9 | 47.3 | 42.5 | 38.9 |
| GPX8 | Canis lupus familiaris OX=9615 | Glutathione peroxidase | 25.8 | 34.4 | 38.8 | 38.8 |
| AHNAK | Canis lupus familiaris OX=9615 | AHNAK nucleoprotein | 11.5 | 6 | 18 | 38.6 |
| HSPA8 | Canis lupus familiaris OX=9615 | Heat shock protein family A (Hsp70) member 8 | 30 | 24.3 | 35.6 | 38.5 |
| FKBP7 | Canis lupus familiaris OX=9615 | Peptidylprolyl isomerase | 41.7 | 48.2 | 34.9 | 38.5 |
| POGLUT1 | Canis lupus familiaris OX=9615 | Protein O-glucosyltransferase 1 | 23.2 | 35.6 | 31.9 | 38.2 |
| POGLUT1 | Canis lupus familiaris OX=9615 | Protein O-glucosyltransferase 1 | 29.3 | 35.6 | 31.9 | 38.2 |
| ANXA1 | Canis lupus familiaris OX=9615 | Annexin | 17.1 | 25.2 | 37.1 | 38 |
| ST3GAL4 | Canis lupus familiaris OX=9615 | ST3 beta-galactoside alpha-2,3-sialyltransferase 4 | 10.3 | 10.7 | 40 | 37.9 |
| CANX | Canis lupus familiaris OX=9615 | Calnexin | 33.3 | 30.1 | 41.7 | 37.6 |
| CLU | Canis lupus familiaris OX=9615 | Clusterin | 25.2 | 26.3 | 37.1 | 37.1 |
| TOR1AIP1 | Canis lupus familiaris OX=9615 | Torsin 1A interacting protein 1 | 42.7 | 32.5 | 35.5 | 36.9 |
| ERLEC1 | Canis lupus familiaris OX=9615 | Endoplasmic reticulum lectin 1 | 37.7 | 38.4 | 38.6 | 36.8 |
| ERO1A | Canis lupus familiaris OX=9615 | Endoplasmic reticulum oxidoreductase 1 alpha | 49.1 | 51.9 | 43.8 | 36.8 |
| MIA3 | Canis lupus familiaris OX=9615 | MIA SH3 domain ER export factor 3 | 25.3 | 23.7 | 35.2 | 36.6 |
| AGR2 | Canis lupus familiaris OX=9615 | Anterior gradient 2, protein disulphide isomerase family member | 52.6 | 53.1 | 36.6 | 36.6 |
| HTATIP2 | Canis lupus familiaris OX=9615 | HIV-1 Tat interactive protein 2 | 21.5 | 20.2 | 18.9 | 35.7 |
| ADAM17 | Canis lupus familiaris OX=9615 | ADAM metallopeptidase domain 17 | 17.4 | 13.7 | 30.1 | 34.9 |
| HIST1H1C | Canis lupus familiaris OX=9615 | H15 domain-containing protein | 49.3 | 48.8 | 36.2 | 34.7 |
| EIF2AK3 | Canis lupus familiaris OX=9615 | Eukaryotic translation initiation factor 2 alpha kinase 3 | 26.3 | 26.8 | 33 | 34 |
| MAN2A1 | Canis lupus familiaris OX=9615 | Alpha-mannosidase | 20.6 | 20.5 | 32.9 | 33.9 |
| LOC489992 | Canis lupus familiaris OX=9615 | Uncharacterized protein | 25.9 | 40.9 | 36.9 | 33.8 |
| LOC488269 | Canis lupus familiaris OX=9615 | H15 domain-containing protein | 43.4 | 47 | 35.2 | 33.8 |
| LMAN1 | Canis lupus familiaris OX=9615 | Lectin, mannose binding 1 | 19.3 | 29.9 | 34.4 | 33.6 |
| FAM3C | Canis lupus familiaris OX=9615 | Family with sequence similarity 3 member C | 37.9 | 24.7 | 33.5 | 33.5 |
| IKBIP | Canis lupus familiaris OX=9615 | IKBKB interacting protein | 27.7 | 24.3 | 28.8 | 33.3 |
| CALR | Canis lupus familiaris OX=9615 | Calreticulin | 33 | 34 | 35.7 | 33.3 |
| SDF2 | Canis lupus familiaris OX=9615 | Stromal cell derived factor 2 | 33.2 | 29.4 | 39.8 | 32.7 |
| P3H2 | Canis lupus familiaris OX=9615 | Prolyl 3-hydroxylase 2 | 21.6 | 36.2 | 37.7 | 32.7 |
| RPL6 | Canis lupus familiaris OX=9615 | 60S ribosomal protein L6 | 13.5 | 15.3 | 35.4 | 32.6 |
| EMD | Canis lupus familiaris OX=9615 | Emerin | 45.2 | 43 | 32.6 | 32.6 |
| TMTC3 | Canis lupus familiaris OX=9615 | Transmembrane and tetratricopeptide repeat containing 3 | 24.5 | 34 | 33.2 | 32.5 |
| PRDX4 | Canis lupus familiaris OX=9615 | Peroxiredoxin 4 | 35.7 | 37.5 | 34.9 | 32.4 |
| HNRNPA3 | Canis lupus familiaris OX=9615 | Heterogeneous nuclear ribonucleoprotein A3 | 14.5 | 18.3 | 32.5 | 32.3 |
| EXTL2 | Canis lupus familiaris OX=9615 | Exostosin like glycosyltransferase 2 | 8.5 | 25.9 | 35.8 | 32.3 |
| ERP29 | Canis lupus familiaris OX=9615 | Endoplasmic reticulum protein 29 | 18.8 | 20.2 | 32.2 | 32.2 |
| HNRNPA2B1 | Canis lupus familiaris OX=9615 | Heterogeneous nuclear ribonucleoprotein A2/B1 | 19.6 | 7 | 44.5 | 31.9 |
| EDEM3 | Canis lupus familiaris OX=9615 | alpha-1,2-Mannosidase | 16 | 24.1 | 32.7 | 31.9 |
| MMP13 | Canis lupus familiaris OX=9615 | Matrix metallopeptidase 13 | 28.2 | 27.6 | 43.3 | 31.7 |
| EXT2 | Canis lupus familiaris OX=9615 | Exostosin glycosyltransferase 2 | 6.7 | 16.6 | 34.2 | 31.5 |
| TOR1B | Canis lupus familiaris OX=9615 | Torsin | 10.2 | 29.2 | 34.8 | 31.3 |
| LOC483167 | Canis lupus familiaris OX=9615 | Histone H4 | 33.3 | 12.5 | 31.2 | 31.2 |
| GLCE | Canis lupus familiaris OX=9615 | Glucuronic acid epimerase | 29.4 | 43.7 | 36.5 | 31.2 |
| PRKCSH | Canis lupus familiaris OX=9615 | Protein kinase C substrate 80K-H | 25 | 22.5 | 39.7 | 31.1 |
| GALNT2 | Canis lupus familiaris OX=9615 | Polypeptide N-acetylgalactosaminyltransferase | 20.1 | 23 | 33.6 | 31 |
| TXNL1 | Canis lupus familiaris OX=9615 | Thioredoxin like 1 | 16 | 7 | 21 | 30.8 |
| CNPY3 | Canis lupus familiaris OX=9615 | Canopy FGF signaling regulator 3 | 29.4 | 37.3 | 25.4 | 30.8 |
| EMC7 | Canis lupus familiaris OX=9615 | ER membrane protein complex subunit 7 | 13.9 | 15 | 21.7 | 30.7 |
| DNAJB11 | Canis lupus familiaris OX=9615 | DnaJ homolog subfamily B member 11 | 30.2 | 25.7 | 38.3 | 30.7 |
| POGLUT3 | Canis lupus familiaris OX=9615 | KDEL motif containing 2 | 9.8 | 18.3 | 20.9 | 30.3 |
| TMPO | Canis lupus familiaris OX=9615 | Thymopoietin | 38.8 | 40.5 | 28 | 30 |
| RPS27 | Canis lupus familiaris OX=9615 | 40S ribosomal protein S27 | 13.1 | 14.3 | 29.8 | 29.8 |
| JUP | Canis lupus familiaris OX=9615 | Junction plakoglobin | 16.5 | 21.6 | 31.6 | 29.7 |
| CLCC1 | Canis lupus familiaris OX=9615 | Chloride channel CLIC like 1 | 21.8 | 19.2 | 27.3 | 29.5 |
| ASPH | Canis lupus familiaris OX=9615 | Aspartate beta-hydroxylase | 27.3 | 27.5 | 31.8 | 29.5 |
| FKBP10 | Canis lupus familiaris OX=9615 | Peptidylprolyl isomerase | 31.8 | 48.1 | 28.5 | 29.4 |
| PLOD3 | Canis lupus familiaris OX=9615 | Procollagen-lysine,2-oxoglutarate 5-dioxygenase 3 | 9.9 | 13.6 | 26.2 | 29.3 |
| LAMB3 | Canis lupus familiaris OX=9615 | Laminin subunit beta 3 | 17.9 | 19.4 | 31.7 | 29.2 |
| FUCA1 | Canis lupus familiaris OX=9615 | Alpha-L-fucosidase | 9.7 | 25.8 | 23.6 | 29.2 |
| SCPEP1 | Canis lupus familiaris OX=9615 | Carboxypeptidase | 15.4 | 21.7 | 29 | 29 |
| SSR4 | Canis lupus familiaris OX=9615 | Signal sequence receptor subunit 4 | 17.9 | 16.8 | 28.9 | 28.9 |
| L1CAM | Canis lupus familiaris OX=9615 | L1 cell adhesion molecule | 17.2 | 18 | 25.4 | 28.9 |
| CASC4 | Canis lupus familiaris OX=9615 | Uncharacterized protein | 32.6 | 29.4 | 27.1 | 28.4 |
| GGH | Canis lupus familiaris OX=9615 | Folate gamma-glutamyl hydrolase | 13.2 | 15.1 | 28.3 | 28.3 |
| KRT42 | Canis lupus familiaris OX=9615 | IF rod domain-containing protein | 22.1 | 24.3 | 24.1 | 28.3 |
| ERAP2 | Canis lupus familiaris OX=9615 | Aminopeptidase | 7.3 | 13 | 23.5 | 27.9 |
| RPS9 | Canis lupus familiaris OX=9615 | Ribosomal protein S9 | 7.7 | 7.7 | 23.2 | 27.8 |
| RPL3 | Canis lupus familiaris OX=9615 | Uncharacterized protein | 8.9 | 7.9 | 33.3 | 27.8 |
| ST3GAL6 | Canis lupus familiaris OX=9615 | ST3 beta-galactoside alpha-2,3-sialyltransferase 6 | 12.7 | 18.7 | 23.6 | 27.8 |
| SELENOT | Canis lupus familiaris OX=9615 | Selenoprotein T | 10.8 | 16.9 | 23.6 | 27.7 |
| DNAJC10 | Canis lupus familiaris OX=9615 | DnaJ homolog subfamily C member 10 | 21.8 | 24.9 | 28.7 | 27.5 |
| DNASE1L1 | Canis lupus familiaris OX=9615 | Deoxyribonuclease | 16.9 | 31.9 | 24.7 | 27.5 |
| DSP | Canis lupus familiaris OX=9615 | Desmoplakin | 10.8 | 11.7 | 23.5 | 27.3 |
| ST3GAL1 | Canis lupus familiaris OX=9615 | ST3 beta-galactoside alpha-2,3-sialyltransferase 1 | 8.5 | 13.2 | 26.9 | 26.9 |

| | | | | | | |
|---|---|---|---|---|---|---|
| CPM | Canis lupus familiaris OX=9615 | Carboxypeptidase M | 11.4 | 19.3 | 25 | 26.9 |
| RACK1 | Canis lupus familiaris OX=9615 | Receptor for activated C kinase 1 | 9.1 | 5 | 24.6 | 26.8 |
| POGLUT2 | Canis lupus familiaris OX=9615 | KDEL motif containing 1 | 6 | 10.8 | 30.1 | 26.7 |
| MCFD2 | Canis lupus familiaris OX=9615 | Multiple coagulation factor deficiency 2 | 21.2 | 10 | 10 | 26.5 |
| HNRNPA1 | Canis lupus familiaris OX=9615 | Uncharacterized protein | 18.5 | 18.5 | 29 | 26.5 |
| GM2A | Canis lupus familiaris OX=9615 | GM2 ganglioside activator | 26.5 | 22.4 | 30.6 | 26.5 |
| MGAT1 | Canis lupus familiaris OX=9615 | Uncharacterized protein | 13 | 16.3 | 26.2 | 26.4 |
| PTGS2 | Canis lupus familiaris OX=9615 | Cyclooxygenase 2 | 18.9 | 19.2 | 26 | 26.3 |
| TAPBP | Canis lupus familiaris OX=9615 | Tapasin | 11.8 | 20 | 23.8 | 26.3 |
| LOX | Canis lupus familiaris OX=9615 | Lysyl oxidase | 16.4 | 13 | 27.1 | 26.2 |
| PBXIP1 | Canis lupus familiaris OX=9615 | PBX homeobox interacting protein 1 | 18.9 | 22.7 | 33.4 | 26.2 |
| DAG1 | Canis lupus familiaris OX=9615 | Dystroglycan | 10.9 | 12.7 | 24.2 | 26 |
| TGOLN2 | Canis lupus familiaris OX=9615 | Uncharacterized protein | 31.7 | 22.5 | 30.5 | 25.9 |
| LAMC2 | Canis lupus familiaris OX=9615 | Laminin subunit gamma 2 | 13.6 | 14.5 | 20 | 25.8 |
| B4GALT1 | Canis lupus familiaris OX=9615 | Beta-1,4-galactosyltransferase 1 | 26 | 19 | 25.6 | 25.6 |
| LMAN2L | Canis lupus familiaris OX=9615 | Lectin, mannose binding 2 like | 17.7 | 24.5 | 27.9 | 25.6 |
| MANBA | Canis lupus familiaris OX=9615 | Mannosidase beta | 17.7 | 19.9 | 16.4 | 25.5 |
| RPN2 | Canis lupus familiaris OX=9615 | Dolichyl-diphosphooligosaccharide--protein glycosyltransferase subunit 2 | 13.9 | 18.7 | 15.8 | 25.3 |
| TTC17 | Canis lupus familiaris OX=9615 | Tetratricopeptide repeat domain 17 | 12 | 19.2 | 20.9 | 25.3 |
| TTC17 | Canis lupus familiaris OX=9615 | Tetratricopeptide repeat domain 17 | 14.7 | 19.2 | 20.9 | 25.3 |
| FGFBP1 | Canis lupus familiaris OX=9615 | Fibroblast growth factor binding protein 1 | 25.1 | 20 | 12.9 | 25.1 |
| CLPTM1 | Canis lupus familiaris OX=9615 | CLPTM1 regulator of GABA type A receptor forward trafficking | 8.2 | 10.8 | 26 | 25 |
| ABHD12 | Canis lupus familiaris OX=9615 | Abhydrolase domain containing 12 | 6.8 | 12.2 | 16.5 | 25 |
| RPL23 | Canis lupus familiaris OX=9615 | 60S ribosomal protein L23 | 10.7 | 12.9 | 19.3 | 25 |
| MOGS | Canis lupus familiaris OX=9615 | Mannosyl-oligosaccharide glucosidase | 12.9 | 25.6 | 30.4 | 25 |
| SRSF3 | Canis lupus familiaris OX=9615 | Serine and arginine rich splicing factor 3 | 24.4 | 34.8 | 24.4 | 25 |
| TXNDC5 | Canis lupus familiaris OX=9615 | Thioredoxin domain containing 5 | 31.3 | 42.7 | 40.9 | 24.8 |
| NUDC | Canis lupus familiaris OX=9615 | Nuclear distribution C, dynein complex regulator | 28 | 16.6 | 15.7 | 24.7 |
| LMAN2 | Canis lupus familiaris OX=9615 | Vesicular integral-membrane protein VIP36 | 9.8 | 20.8 | 21.1 | 24.7 |
| LOC10215262( | Canis lupus familiaris OX=9615 | Polypeptide N-acetylgalactosaminyltransferase | 19.6 | 23.7 | 27.1 | 24.5 |
| ERO1B | Canis lupus familiaris OX=9615 | Endoplasmic reticulum oxidoreductase 1 beta | 21.6 | 20.3 | 29.6 | 24.4 |
| TFRC | Canis lupus familiaris OX=9615 | Transferrin receptor protein 1 | 9.5 | 7.9 | 21.2 | 24.3 |
| TMED10 | Canis lupus familiaris OX=9615 | Transmembrane p24 trafficking protein 10 | 24.1 | 35.7 | 24.1 | 24.1 |
| UGGT2 | Canis lupus familiaris OX=9615 | UDP-glucose glycoprotein glucosyltransferase 2 | 16.3 | 15.2 | 20.3 | 23.9 |
| STIM1 | Canis lupus familiaris OX=9615 | Stromal interaction molecule 1 | 17.8 | 22.9 | 24.1 | 23.8 |
| B3GAT3 | Canis lupus familiaris OX=9615 | Galactosylgalactosylxylosylprotein 3-beta-glucuronosyltransferase | 8.1 | 9.9 | 21.2 | 23.6 |
| CCDC134 | Canis lupus familiaris OX=9615 | Uncharacterized protein | 15.7 | 31.7 | 23.2 | 23.2 |
| ADAM10 | Canis lupus familiaris OX=9615 | ADAM metallopeptidase domain 10 | 24.7 | 22.4 | 23.2 | 22.9 |
| NUDT9 | Canis lupus familiaris OX=9615 | Nudix hydrolase 9 | 22 | 29.4 | 23.7 | 22.9 |
| VDAC2 | Canis lupus familiaris OX=9615 | Voltage dependent anion channel 2 | 6.8 | 15.6 | 25.5 | 22.8 |
| GLB1L | Canis lupus familiaris OX=9615 | Galactosidase beta 1 like | 10.1 | 12.7 | 24.7 | 22.7 |
| SUN2 | Canis lupus familiaris OX=9615 | Sad1 and UNC84 domain containing 2 | 17 | 12.8 | 26.1 | 22.6 |
| CTSL | Canis lupus familiaris OX=9615 | Cathepsin L1 | 25.5 | 26.7 | 23.7 | 22.5 |
| DNAJB9 | Canis lupus familiaris OX=9615 | DnaJ heat shock protein family (Hsp40) member B9 | 31.5 | 36.5 | 22.5 | 22.5 |
| TMPRSS11E | Canis lupus familiaris OX=9615 | Transmembrane protease serine | 8.5 | 7.6 | 14 | 22.3 |
| RPL8 | Canis lupus familiaris OX=9615 | Ribosomal protein L8 | 9.3 | 9.3 | 22.2 | 22.2 |
| TOR1A | Canis lupus familiaris OX=9615 | Torsin | 5.2 | 16 | 27.4 | 22.2 |
| SERPINE2 | Canis lupus familiaris OX=9615 | Serpin family E member 2 | 22.4 | 22.4 | 22.7 | 22.2 |
| SLC38A10 | Canis lupus familiaris OX=9615 | Solute carrier family 38 member 10 | 11.9 | 10.8 | 21.2 | 22.1 |
| WNT7A | Canis lupus familiaris OX=9615 | Protein Wnt | 13.2 | 16.6 | 27.2 | 22.1 |
| LAMA3 | Canis lupus familiaris OX=9615 | Laminin subunit alpha 3 | 8.1 | 9.8 | 19.1 | 22 |
| EPCAM | Canis lupus familiaris OX=9615 | Epithelial cell adhesion molecule | 22 | 11.4 | 22 | 22 |
| TCN2 | Canis lupus familiaris OX=9615 | Transcobalamin 2 | 11.1 | 12.7 | 22.2 | 22 |
| RAB10 | Canis lupus familiaris OX=9615 | Ras-related protein Rab-10 | 6 | 19 | 42 | 22 |
| DLA-64 | Canis lupus familiaris OX=9615 | MHC class I antigen | 22.4 | 13.3 | 30.5 | 21.9 |
| SUMF1 | Canis lupus familiaris OX=9615 | Sulfatase modifying factor 1 | 33.7 | 42.5 | 30.7 | 21.9 |
| IMPAD1 | Canis lupus familiaris OX=9615 | Inositol monophosphatase domain containing 1 | 11.4 | 16.4 | 28.9 | 21.7 |
| GUSB | Canis lupus familiaris OX=9615 | Beta-glucuronidase | 11.8 | 21.8 | 17.7 | 21.7 |
| SPCS3 | Canis lupus familiaris OX=9615 | Signal peptidase complex subunit 3 | 21.7 | 35 | 30.6 | 21.7 |
| SEL1L | Canis lupus familiaris OX=9615 | SEL1L adaptor subunit of ERAD E3 ubiquitin ligase | 19.6 | 21.9 | 20.7 | 21.5 |
| CNPY2 | Canis lupus familiaris OX=9615 | Saposin B-type domain-containing protein | 22.9 | 7.4 | 32.4 | 21.3 |
| MGAT2 | Canis lupus familiaris OX=9615 | Mannosyl (alpha-1,6-)-glycoprotein beta-1,2-N-acetylglucosaminyltransferase | 9 | 16.1 | 19.1 | 21.3 |
| RBMX | Canis lupus familiaris OX=9615 | RNA binding motif protein X-linked | 12.2 | 13 | 26.3 | 21.2 |
| ZCCHC17 | Canis lupus familiaris OX=9615 | Uncharacterized protein | 16.2 | 16.2 | 19.9 | 21.2 |
| MINPP1 | Canis lupus familiaris OX=9615 | Multiple inositol-polyphosphate phosphatase 1 | 10.6 | 10.6 | 17.5 | 20.8 |
| CISD2 | Canis lupus familiaris OX=9615 | CDGSH iron sulfur domain 2 | 25.9 | 20.7 | 20.7 | 20.7 |
| CCDC47 | Canis lupus familiaris OX=9615 | Coiled-coil domain containing 47 | 22.3 | 15.1 | 17.1 | 20.6 |
| PLA2G7 | Canis lupus familiaris OX=9615 | Platelet-activating factor acetylhydrolase | 5.6 | 18.2 | 6.8 | 20.5 |
| MANF | Canis lupus familiaris OX=9615 | Mesencephalic astrocyte derived neurotrophic factor | 20.5 | 24.9 | 23.8 | 20.5 |
| P3H1 | Canis lupus familiaris OX=9615 | Prolyl 3-hydroxylase 1 | 9.3 | 14.5 | 19.1 | 20.3 |
| BCHE | Canis lupus familiaris OX=9615 | Carboxylic ester hydrolase | 17.3 | 19.9 | 19.1 | 20.3 |
| MYDGF | Canis lupus familiaris OX=9615 | Myeloid derived growth factor | 27.2 | 20.2 | 20.2 | 20.2 |
| FUT8 | Canis lupus familiaris OX=9615 | Alpha-(1,6)-fucosyltransferase | 8.2 | 7.8 | 13.9 | 20 |
| PLOD2 | Canis lupus familiaris OX=9615 | Procollagen-lysine,2-oxoglutarate 5-dioxygenase 2 | 12 | 17 | 22.8 | 20 |
| ERLIN1 | Canis lupus familiaris OX=9615 | ER lipid raft associated 1 | 9.7 | 9.7 | 26.1 | 19.9 |
| CRTAP | Canis lupus familiaris OX=9615 | Cartilage associated protein | 6.1 | 6.1 | 21.6 | 19.7 |
| MANEA | Canis lupus familiaris OX=9615 | Mannosidase endo-alpha | 12.1 | 22.1 | 24.5 | 19.7 |

| | | | | | | |
|---|---|---|---|---|---|---|
| CD74 | Canis lupus familiaris OX=9615 | CD74 molecule | 16.2 | 19.5 | 19.5 | 19.5 |
| TIMP3 | Canis lupus familiaris OX=9615 | TIMP metallopeptidase inhibitor 3 | 13.7 | 16.1 | 19.4 | 19.4 |
| MFGE8 | Canis lupus familiaris OX=9615 | Milk fat globule-EGF factor 8 protein | 19.8 | 27.4 | 19.3 | 19.3 |
| RPL5 | Canis lupus familiaris OX=9615 | Ribosomal protein L5 | 12.1 | 16.2 | 19.2 | 19.2 |
| MUC1 | Canis lupus familiaris OX=9615 | Mucin 1, cell surface associated | 18.2 | 11 | 15.1 | 19 |
| RPS11 | Canis lupus familiaris OX=9615 | 40S ribosomal protein S11 | 12.7 | 13.3 | 29.1 | 19 |
| CDCP1 | Canis lupus familiaris OX=9615 | CUB domain containing protein 1 | 8.4 | 7.5 | 16.9 | 18.8 |
| TNFSF10 | Canis lupus familiaris OX=9615 | Tumor necrosis factor ligand superfamily member | 22.1 | 13.9 | 28.8 | 18.5 |
| MYO6 | Canis lupus familiaris OX=9615 | Myosin VI | 6.2 | 5 | 13.5 | 18.4 |
| ADAM9 | Canis lupus familiaris OX=9615 | ADAM metallopeptidase domain 9 | 8.5 | 11.2 | 16.6 | 18.4 |
| EOGT | Canis lupus familiaris OX=9615 | EGF domain-specific O-linked N-acetylglucosamine transferase | 6.5 | 12.9 | 20.5 | 18.4 |
| THBS1 | Canis lupus familiaris OX=9615 | Thrombospondin 1 | 5.9 | 7 | 16.2 | 18.1 |
| CTSH | Canis lupus familiaris OX=9615 | Cathepsin H | 10.8 | 14.8 | 18.7 | 17.7 |
| BLMH | Canis lupus familiaris OX=9615 | Bleomycin hydrolase | 10.5 | 12.3 | 18.2 | 17.6 |
| ATP1B1 | Canis lupus familiaris OX=9615 | Sodium/potassium-transporting ATPase subunit beta-1 | 9.7 | 9.7 | 17.5 | 17.5 |
| MMP15 | Canis lupus familiaris OX=9615 | Matrix metallopeptidase 15 | 18.9 | 10.5 | 22.1 | 17.5 |
| ERBB3 | Canis lupus familiaris OX=9615 | Erb-b2 receptor tyrosine kinase 3 | 5.1 | 8 | 11.6 | 17.4 |
| SUMF2 | Canis lupus familiaris OX=9615 | Sulfatase modifying factor 2 | 18.3 | 8.3 | 22.3 | 17.3 |
| UPK1B | Canis lupus familiaris OX=9615 | Tetraspanin | 10 | 13.8 | 17.3 | 17.3 |
| KRT75 | Canis lupus familiaris OX=9615 | Keratin 75 | 8.7 | 14.5 | 25.7 | 17.2 |
| B3GLCT | Canis lupus familiaris OX=9615 | Beta 3-glucosyltransferase | 7.9 | 22.4 | 14.3 | 17 |
| PLOD1 | Canis lupus familiaris OX=9615 | Fe2OG dioxygenase domain-containing protein | 6.6 | 14.2 | 15.7 | 16.8 |
| TXNDC11 | Canis lupus familiaris OX=9615 | Thioredoxin domain containing 11 | 9.2 | 9.9 | 13.4 | 16.7 |
| STC1 | Canis lupus familiaris OX=9615 | Stanniocalcin 1 | 16.6 | 27.1 | 16.6 | 16.6 |
| PPT1 | Canis lupus familiaris OX=9615 | Palmitoyl-protein thioesterase 1 | 19.6 | 14.7 | 13.7 | 16.3 |
| CLIC1 | Canis lupus familiaris OX=9615 | Chloride intracellular channel protein | 12.4 | 7.5 | 12.4 | 16.2 |
| PAM | Canis lupus familiaris OX=9615 | Peptidylglycine alpha-amidating monooxygenase | 6.9 | 6.2 | 15 | 16 |
| TMX1 | Canis lupus familiaris OX=9615 | Thioredoxin related transmembrane protein 1 | 24.1 | 19.8 | 23 | 15.8 |
| PM20D1 | Canis lupus familiaris OX=9615 | Peptidase M20 domain containing 1 | 8.5 | 9.9 | 17.9 | 15.7 |
| IL1RL1 | Canis lupus familiaris OX=9615 | Interleukin 1 receptor like 1 | 13.8 | 15.1 | 18.5 | 15.4 |
| TPST1 | Canis lupus familiaris OX=9615 | Protein-tyrosine sulfotransferase | 11 | 16.3 | 16.5 | 14 |
| SDF2L1 | Canis lupus familiaris OX=9615 | Stromal cell derived factor 2 like 1 | 21 | 21.3 | 24.7 | 14 |
| CLN5 | Canis lupus familiaris OX=9615 | Ceroid-lipofuscinosis neuronal protein 5 | 5.4 | 8 | 13.7 | 13.7 |
| TGFB1 | Canis lupus familiaris OX=9615 | Transforming growth factor beta-1 proprotein | 21 | 21.3 | 14.1 | 13.6 |
| FKBP2 | Canis lupus familiaris OX=9615 | Peptidylprolyl isomerase | 9.2 | 17.7 | 17 | 13.5 |
| RRBP1 | Canis lupus familiaris OX=9615 | Ribosome-binding protein 1 | 15.7 | 10.1 | 9.7 | 13.4 |
| CTSB | Canis lupus familiaris OX=9615 | Cathepsin B | 9.4 | 9.4 | 9.4 | 13.3 |
| GNS | Canis lupus familiaris OX=9615 | N-acetylglucosamine-6-sulfatase | 10.6 | 17.2 | 13.3 | 13.3 |
| ERGIC3 | Canis lupus familiaris OX=9615 | ERGIC and golgi 3 | 13.1 | 11 | 13.1 | 13.1 |
| TGFBR1 | Canis lupus familiaris OX=9615 | Receptor protein serine/threonine kinase | 11.5 | 13.4 | 5.5 | 13.1 |
| LTBP1 | Canis lupus familiaris OX=9615 | Latent transforming growth factor beta binding protein 1 | 8.5 | 10.5 | 13.1 | 12.9 |
| RNF121 | Canis lupus familiaris OX=9615 | Ring finger protein 121 | 12.8 | 12.8 | 12.8 | 12.8 |
| TMEM106B | Canis lupus familiaris OX=9615 | Uncharacterized protein | 15.7 | 15.7 | 12.3 | 12.6 |
| LMLN | Canis lupus familiaris OX=9615 | Leishmanolysin like peptidase | 9.9 | 6.5 | 16.1 | 12.5 |
| FKBP9 | Canis lupus familiaris OX=9615 | Peptidylprolyl isomerase | 6.1 | 12.3 | 13.9 | 12.3 |
| NPC2 | Canis lupus familiaris OX=9615 | NPC intracellular cholesterol transporter 2 | 26.8 | 13.4 | 13.4 | 12.1 |
| VDAC1 | Canis lupus familiaris OX=9615 | Voltage dependent anion channel 1 | 7.1 | 11 | 19.8 | 12 |
| SCRN1 | Canis lupus familiaris OX=9615 | Peptidylprolyl isomerase | 19 | 19 | 7.1 | 11.8 |
| CD99 | Canis lupus familiaris OX=9615 | CD99 molecule | 11.6 | 11.6 | 11.6 | 11.6 |
| MGAT4B | Canis lupus familiaris OX=9615 | Alpha-1,3-mannosyl-glycoprotein 4-beta-N-acetylglucosaminyltransferase B | 6.6 | 13.7 | 8.2 | 11.5 |
| SEC63 | Canis lupus familiaris OX=9615 | Translocation protein SEC63 homolog | 5 | 5 | 11.4 | 11.4 |
| NXPE3 | Canis lupus familiaris OX=9615 | Neurexophilin and PC-esterase domain family member 3 | 19.7 | 21.3 | 13.1 | 11.3 |
| AGRN | Canis lupus familiaris OX=9615 | Agrin | 7.3 | 7.9 | 7.3 | 10.7 |
| IGFBP7 | Canis lupus familiaris OX=9615 | Insulin like growth factor binding protein 7 | 13.1 | 13.1 | 5 | 10.6 |
| CCN1 | Canis lupus familiaris OX=9615 | Cellular communication network factor 1 | 5.2 | 7.1 | 10.5 | 10.5 |
| TAPBPL | Canis lupus familiaris OX=9615 | TAP binding protein like | 10.5 | 10.5 | 10.5 | 10.5 |
| ORMDL2 | Canis lupus familiaris OX=9615 | ORM1-like protein | 9.8 | 9.8 | 9.8 | 9.8 |
| LAMB1 | Canis lupus familiaris OX=9615 | Laminin subunit beta 1 | 7.1 | 7.3 | 14 | 9.5 |
| HYAL2 | Canis lupus familiaris OX=9615 | Hyaluronidase | 5 | 7.6 | 9.2 | 9.2 |
| COL5A2 | Canis lupus familiaris OX=9615 | Collagen type V alpha 2 chain | 12.3 | 5.3 | 10 | 9.1 |
| GALNT3 | Canis lupus familiaris OX=9615 | Polypeptide N-acetylgalactosaminyltransferase | 5.2 | 7.6 | 12.2 | 9 |
| GPC1 | Canis lupus familiaris OX=9615 | Glypican 1 | 9.8 | 9 | 5.3 | 9 |
| FAU | Canis lupus familiaris OX=9615 | 40S ribosomal protein S30 | 14.3 | 9 | 8.3 | 9 |
| SLC16A1 | Canis lupus familiaris OX=9615 | Solute carrier family 16 member 1 | 8.6 | 8.6 | 12.7 | 8.8 |
| B2M | Canis lupus familiaris OX=9615 | Beta-2-microglobulin | 8.8 | 8.8 | 16.8 | 8.8 |
| PTK7 | Canis lupus familiaris OX=9615 | Uncharacterized protein | 9.8 | 4.8 | 8.7 | 8.4 |
| CLSTN1 | Canis lupus familiaris OX=9615 | Uncharacterized protein | 9.1 | 8.1 | 11.8 | 8.3 |
| RCN2 | Canis lupus familiaris OX=9615 | Reticulocalbin 2 | 21.1 | 20 | 8.3 | 8.3 |
| TMTC2 | Canis lupus familiaris OX=9615 | Transmembrane and tetratricopeptide repeat containing 2 | 5.3 | 5.1 | 8.4 | 8.1 |
| PCSK7 | Canis lupus familiaris OX=9615 | Proprotein convertase subtilisin/kexin type 7 | 5.2 | 5.2 | 6.9 | 8.1 |
| CCPG1 | Canis lupus familiaris OX=9615 | Cell cycle progression 1 | 7.5 | 5.6 | 10.8 | 7.4 |
| EGFL7 | Canis lupus familiaris OX=9615 | EGF like domain multiple 7 | 6.9 | 10.6 | 6.9 | 6.9 |
| SFTPD | Canis lupus familiaris OX=9615 | Surfactant protein D | 17.6 | 18.7 | 12.5 | 6.9 |
| POFUT2 | Canis lupus familiaris OX=9615 | Protein O-fucosyltransferase 2 | 7.8 | 25.8 | 13.2 | 6.3 |
| SPINT2 | Canis lupus familiaris OX=9615 | Serine peptidase inhibitor, Kunitz type 2 | 5.6 | 10.3 | 5.6 | 5.6 |

**Complete proteomic list obtained from MS analysis of C-BioID**

**Bold**- Common hits in both EC-BioID and C-BioID; *Italic* - Transmembrane proteins in C-BioID only

| C-BioID | | | | |
|---|---|---|---|---|
| | | | **Sample 1** | **Sample2** |
| **Gene** | **Organism** | **Protein Description** | **Percent Coverage** | **Percent Coverage** |
| **DSG2** | Canis lupus familiaris OX=9615 | Desmoglein 2 | 54.2 | 61.1 |
| **CXADR** | Canis lupus familiaris OX=9615 | Coxsackievirus and adenovirus receptor | 53.4 | 55.3 |
| **CDH1** | Canis lupus familiaris OX=9615 | Cadherin-1 | 30.7 | 27.2 |
| **DSC3** | Canis lupus familiaris OX=9615 | Desmocollin 3 | 22.7 | 23.9 |
| **ITGA2** | Canis lupus familiaris OX=9615 | Integrin subunit alpha 2 | 18.2 | 18.2 |
| **EFNB1** | Canis lupus familiaris OX=9615 | Ephrin B1 | 18 | 18 |
| **ITGB1** | Canis lupus familiaris OX=9615 | Integrin beta 1 | 11.9 | 10.2 |
| **CDH3** | Canis lupus familiaris OX=9615 | Cadherin 3 | 7 | 12.4 |
| *PCDH1* | Canis lupus familiaris OX=9615 | Protocadherin 1 | 17 | 19 |
| *NECTIN2* | Canis lupus familiaris OX=9615 | Nectin cell 2 | 44.5 | 38.1 |
| *DSC2* | Canis lupus familiaris OX=9615 | Desmocollin 2 | 16.9 | 19.7 |
| *NECTIN3* | Canis lupus familiaris OX=9615 | Nectin 3 | 14 | 18.4 |
| *CDH2* | Canis lupus familiaris OX=9615 | Cadherin 2 | 7.2 | 10.8 |
| *OCLN* | Canis lupus familiaris OX=9615 | Occludin | 21.7 | 19.2 |
| *NECTIN1* | Canis lupus familiaris OX=9615 | Nectin-1 | 6 | 7.2 |
| *DSG1* | Canis lupus familiaris OX=9615 | Desmoglein-1 | 8.7 | 4.8 |
| AHNAK | Canis lupus familiaris OX=9615 | AHNAK nucleoprotein | 88.4 | 88.9 |
| MISP | Canis lupus familiaris OX=9615 | Mitotic spindle positioning | 76.7 | 66 |
| AHNAK2 | Canis lupus familiaris OX=9615 | PDZ domain-containing protein | 72.2 | 77.4 |
| CDV3 | Canis lupus familiaris OX=9615 | CDV3 homolog | 69.2 | 69.2 |
| KIAA1217 | Canis lupus familiaris OX=9615 | KIAA1217 | 66.4 | 66 |
| TJP1 | Canis lupus familiaris OX=9615 | Tight junction protein ZO-1 | 66.4 | 68.7 |
| KIAA1217 | Canis lupus familiaris OX=9615 | KIAA1217 | 66.3 | 66 |
| DBI | Canis lupus familiaris OX=9615 | Acyl-CoA-binding protein | 65.5 | 54 |
| SNAP23 | Canis lupus familiaris OX=9615 | Synaptosomal-associated protein | 65.4 | 72.5 |
| KRT8 | Canis lupus familiaris OX=9615 | IF rod domain-containing protein | 63.7 | 52.1 |
| CTNNA1 | Canis lupus familiaris OX=9615 | Catenin alpha 1 | 63.6 | 70.6 |
| ENSA | Canis lupus familiaris OX=9615 | Endosulfine alpha | 62.8 | 67.8 |
| ANXA2 | Canis lupus familiaris OX=9615 | Annexin A2 | 61.7 | 65.5 |
| LOC10215270[illegible] | Canis lupus familiaris OX=9615 | Thymosin beta | 61.4 | 61.4 |
| ZDHHC5 | Canis lupus familiaris OX=9615 | Palmitoyltransferase ZDHHC5 | 61 | 39.7 |
| PALM | Canis lupus familiaris OX=9615 | Paralemmin | 60.2 | 61.5 |
| CD2AP | Canis lupus familiaris OX=9615 | CD2 associated protein | 59.9 | 57.1 |
| HS1BP3 | Canis lupus familiaris OX=9615 | HCLS1 binding protein 3 | 59.7 | 56.1 |
| DHRS13 | Canis lupus familiaris OX=9615 | Flotillin | 59.3 | 59.6 |
| ZDHHC5 | Canis lupus familiaris OX=9615 | Palmitoyltransferase ZDHHC5 | 58.8 | 39.7 |
| PDLIM1 | Canis lupus familiaris OX=9615 | PDZ and LIM domain 1 | 58.7 | 55.8 |
| EFHD2 | Canis lupus familiaris OX=9615 | EF-hand domain family member D2 | 58 | 63.8 |
| YAP1 | Canis lupus familiaris OX=9615 | Yes associated protein 1 | 57.8 | 52.6 |
| SLC9A3R1 | Canis lupus familiaris OX=9615 | Na(+)/H(+) exchange regulatory cofactor NHE-RF | 57.8 | 66.5 |
| PHACTR4 | Canis lupus familiaris OX=9615 | Phosphatase and actin regulator | 57.7 | 68.9 |
| BAIAP2L1 | Canis lupus familiaris OX=9615 | BAI1 associated protein 2 like 1 | 56.5 | 57.9 |
| LAD1 | Canis lupus familiaris OX=9615 | Ladinin 1 | 56.3 | 56.7 |
| TAGLN2 | Canis lupus familiaris OX=9615 | Transgelin | 56.3 | 63.8 |
| ERBIN | Canis lupus familiaris OX=9615 | Erbb2 interacting protein | 56.2 | 59.6 |
| KRT14 | Canis lupus familiaris OX=9615 | Keratin 14 | 56 | 45.2 |
| PLEKHA6 | Canis lupus familiaris OX=9615 | Pleckstrin homology domain containing A6 | 55.9 | 52.6 |
| PLEKHA6 | Canis lupus familiaris OX=9615 | Pleckstrin homology domain containing A6 | 55.7 | 52.6 |
| HS1BP3 | Canis lupus familiaris OX=9615 | HCLS1 binding protein 3 | 55.4 | 56.1 |
| CDCA3 | Canis lupus familiaris OX=9615 | Cell division cycle associated 3 | 53.4 | 42.2 |
| LASP1 | Canis lupus familiaris OX=9615 | LIM and SH3 protein 1 | 53.1 | 70.2 |
| MYO6 | Canis lupus familiaris OX=9615 | Myosin VI | 52.2 | 54 |
| SNAP29 | Canis lupus familiaris OX=9615 | Synaptosome associated protein 29 | 51.9 | 65.5 |
| RPS21 | Canis lupus familiaris OX=9615 | 40S ribosomal protein S21 | 51.8 | 53 |
| RIPK1 | Canis lupus familiaris OX=9615 | Receptor interacting serine/threonine kinase 1 | 51.6 | 50.2 |
| SNCG | Canis lupus familiaris OX=9615 | Gamma-synuclein | 51.5 | 59.6 |
| PPP1R2 | Canis lupus familiaris OX=9615 | Uncharacterized protein | 51.2 | 65.9 |
| PPP1R13L | Canis lupus familiaris OX=9615 | Protein phosphatase 1 regulatory subunit 13 like | 50.8 | 50.4 |
| AFDN | Canis lupus familiaris OX=9615 | Afadin, adherens junction formation factor | 50.6 | 52.9 |
| PLEKHA6 | Canis lupus familiaris OX=9615 | Pleckstrin homology domain containing A6 | 50.5 | 52.6 |
| PAK2 | Canis lupus familiaris OX=9615 | Non-specific serine/threonine protein kinase | 50 | 43.9 |
| HSPA8 | Canis lupus familiaris OX=9615 | Heat shock protein family A (Hsp70) member 8 | 49.5 | 47.4 |
| PKP4 | Canis lupus familiaris OX=9615 | Plakophilin 4 | 48.8 | 52.3 |
| JPT1 | Canis lupus familiaris OX=9615 | HN1 | 48.3 | 48.3 |
| C4H1orf198 | Canis lupus familiaris OX=9615 | Chromosome 4 C1orf198 homolog | 48.2 | 48 |
| PFN1 | Canis lupus familiaris OX=9615 | Profilin | 48 | 48 |
| PLEKHA5 | Canis lupus familiaris OX=9615 | Pleckstrin homology domain containing A5 | 48 | 49 |
| ANXA1 | Canis lupus familiaris OX=9615 | Annexin | 47.8 | 44.9 |
| CDC42EP1 | Canis lupus familiaris OX=9615 | CDC42 effector protein 1 | 47.8 | 50.4 |
| FNBP1L | Canis lupus familiaris OX=9615 | Formin binding protein 1 like | 47.8 | 51 |

| | | | | |
|---|---|---|---|---|
| PDLIM7 | Canis lupus familiaris OX=9615 | Uncharacterized protein | 47.7 | 45.6 |
| HSBP1 | Canis lupus familiaris OX=9615 | Heat shock factor binding protein 1 | 47.4 | 47.4 |
| KRT19 | Canis lupus familiaris OX=9615 | Keratin 19 | 47.4 | 50.1 |
| PDLIM5 | Canis lupus familiaris OX=9615 | PDZ and LIM domain 5 | 47.3 | 47.7 |
| TSC22D1 | Canis lupus familiaris OX=9615 | TSC22 domain family member 1 | 47.2 | 38.9 |
| ZDHHC5 | Canis lupus familiaris OX=9615 | Palmitoyltransferase ZDHHC5 | 47 | 39.7 |
| PPFIBP1 | Canis lupus familiaris OX=9615 | PPFIA binding protein 1 | 47 | 45.1 |
| EPS15L1 | Canis lupus familiaris OX=9615 | Epidermal growth factor receptor pathway substrate 15 like 1 | 47 | 50.7 |
| JPT2 | Canis lupus familiaris OX=9615 | Jupiter microtubule associated homolog 2 | 46.8 | 50.6 |
| LPP | Canis lupus familiaris OX=9615 | Uncharacterized protein | 46.7 | 43.8 |
| VAPA | Canis lupus familiaris OX=9615 | VAMP associated protein A | 46.6 | 44.2 |
| PC | Canis lupus familiaris OX=9615 | Pyruvate carboxylase | 46.3 | 48.1 |
| TNKS1BP1 | Canis lupus familiaris OX=9615 | Tankyrase 1 binding protein 1 | 45.6 | 46.9 |
| CAV1 | Canis lupus familiaris OX=9615 | Caveolin-1 | 45.5 | 45.5 |
| KRT5 | Canis lupus familiaris OX=9615 | IF rod domain-containing protein | 45.3 | 36.4 |
| SEPTIN7 | Canis lupus familiaris OX=9615 | Septin 7 | 44.8 | 45.3 |
| TJP2 | Canis lupus familiaris OX=9615 | Tight junction protein ZO-2 | 44.8 | 46.3 |
| LMO7 | Canis lupus familiaris OX=9615 | LIM domain 7 | 44.8 | 46.9 |
| EZR | Canis lupus familiaris OX=9615 | Ezrin | 44.7 | 48 |
| VIM | Canis lupus familiaris OX=9615 | Vimentin | 44.6 | 40.3 |
| PPFIBP1 | Canis lupus familiaris OX=9615 | PPFIA binding protein 1 | 44.5 | 45.1 |
| TJP2 | Canis lupus familiaris OX=9615 | Tight junction protein ZO-2 | 44.3 | 46.3 |
| CCT8 | Canis lupus familiaris OX=9615 | Chaperonin containing TCP1 subunit 8 | 44.2 | 47.3 |
| ABLIM1 | Canis lupus familiaris OX=9615 | Actin binding LIM protein 1 | 43.9 | 37.2 |
| EPB41L5 | Canis lupus familiaris OX=9615 | Band 4.1-like protein 5 | 43.4 | 42.8 |
| EPB41L5 | Canis lupus familiaris OX=9615 | Band 4.1-like protein 5 | 43.4 | 42.8 |
| JUP | Canis lupus familiaris OX=9615 | Junction plakoglobin | 43.4 | 44.5 |
| SEPTIN10 | Canis lupus familiaris OX=9615 | Septin 10 | 43.2 | 35.9 |
| CTTN | Canis lupus familiaris OX=9615 | Cortactin | 43.1 | 43.3 |
| TNRC6B | Canis lupus familiaris OX=9615 | Uncharacterized protein | 42.8 | 47.9 |
| PCCA | Canis lupus familiaris OX=9615 | Propionyl-CoA carboxylase subunit alpha | 42.8 | 56 |
| SHROOM3 | Canis lupus familiaris OX=9615 | Shroom family member 3 | 42.7 | 44.3 |
| EPB41 | Canis lupus familiaris OX=9615 | Protein 4.1 | 42.4 | 30.3 |
| ANK3 | Canis lupus familiaris OX=9615 | Ankyrin 3 | 42.1 | 20.4 |
| BCAR1 | Canis lupus familiaris OX=9615 | BCAR1 scaffold protein, Cas family member | 41.9 | 46.3 |
| DPYSL3 | Canis lupus familiaris OX=9615 | Dihydropyrimidinase like 3 | 41.8 | 38.9 |
| ARHGAP32 | Canis lupus familiaris OX=9615 | Rho GTPase activating protein 32 | 41.7 | 37.5 |
| WASHC2C | Canis lupus familiaris OX=9615 | CAP-ZIP_m domain-containing protein | 41.7 | 42.2 |
| C7H1orf116 | Canis lupus familiaris OX=9615 | Chromosome 7 C1orf116 homolog | 41.7 | 42.9 |
| CAT | Canis lupus familiaris OX=9615 | Catalase | 41.6 | 40.2 |
| SPTAN1 | Canis lupus familiaris OX=9615 | Spectrin alpha, non-erythrocytic 1 | 41.6 | 41.7 |
| DPYSL2 | Canis lupus familiaris OX=9615 | Dihydropyrimidinase like 2 | 41.5 | 42.8 |
| PARD3B | Canis lupus familiaris OX=9615 | Par-3 family cell polarity regulator beta | 41.2 | 41 |
| UPF1 | Canis lupus familiaris OX=9615 | UPF1 RNA helicase and ATPase | 41.1 | 38.3 |
| MICALL2 | Canis lupus familiaris OX=9615 | MICAL like 2 | 41 | 43.2 |
| EPB41L2 | Canis lupus familiaris OX=9615 | Erythrocyte membrane protein band 4.1 like 2 | 40.8 | 33.2 |
| SYNPO | Canis lupus familiaris OX=9615 | Synaptopodin | 40.8 | 35.9 |
| SPTAN1 | Canis lupus familiaris OX=9615 | Spectrin alpha, non-erythrocytic 1 | 40.8 | 41.7 |
| WASL | Canis lupus familiaris OX=9615 | WASP like actin nucleation promoting factor | 40.6 | 30.9 |
| CTNNB1 | Canis lupus familiaris OX=9615 | Catenin beta-1 | 40.5 | 39.2 |
| FAM110B | Canis lupus familiaris OX=9615 | Family with sequence similarity 110 member B | 40.5 | 42.1 |
| ARPP19 | Canis lupus familiaris OX=9615 | cAMP regulated phosphoprotein 19 | 40.2 | 40.2 |
| SNX3 | Canis lupus familiaris OX=9615 | Sorting nexin 3 | 40.1 | 35.2 |
| DBNL | Canis lupus familiaris OX=9615 | Uncharacterized protein | 40.1 | 42 |
| AURKA | Canis lupus familiaris OX=9615 | Aurora kinase A | 40 | 45.1 |
| MID1IP1 | Canis lupus familiaris OX=9615 | MID1 interacting protein 1 | 39.9 | 42.1 |
| CAPZA1 | Canis lupus familiaris OX=9615 | F-actin-capping protein subunit alpha | 39.9 | 46.4 |
| TXNL1 | Canis lupus familiaris OX=9615 | Thioredoxin like 1 | 39.8 | 28.4 |
| FLNA | Canis lupus familiaris OX=9615 | Filamin A | 39.8 | 40.9 |
| NF2 | Canis lupus familiaris OX=9615 | Neurofibromin 2 | 39.1 | 44 |
| SAV1 | Canis lupus familiaris OX=9615 | Salvador family WW domain containing protein 1 | 38.8 | 46.4 |
| IGBP1 | Canis lupus familiaris OX=9615 | Immunoglobulin binding protein 1 | 38.5 | 42.1 |
| LIMA1 | Canis lupus familiaris OX=9615 | LIM domain and actin binding 1 | 38.5 | 43.3 |
| EDC3 | Canis lupus familiaris OX=9615 | Enhancer of mRNA decapping 3 | 38.4 | 30.9 |
| RDX | Canis lupus familiaris OX=9615 | Radixin | 38.4 | 34.9 |
| PPP1R13B | Canis lupus familiaris OX=9615 | Protein phosphatase 1 regulatory subunit 13B | 38.3 | 35.9 |
| CARMIL1 | Canis lupus familiaris OX=9615 | Capping protein regulator and myosin 1 linker 1 | 38.3 | 40.5 |
| CARMIL1 | Canis lupus familiaris OX=9615 | Capping protein regulator and myosin 1 linker 1 | 38.2 | 40.5 |
| MYH9 | Canis lupus familiaris OX=9615 | Myosin-9 | 38.1 | 37.4 |
| WASHC2C | Canis lupus familiaris OX=9615 | CAP-ZIP_m domain-containing protein | 37.9 | 42.2 |
| SYNRG | Canis lupus familiaris OX=9615 | Synergin gamma | 37.9 | 44.1 |
| SNRPD1 | Canis lupus familiaris OX=9615 | Small nuclear ribonucleoprotein Sm D1 | 37.8 | 54.6 |
| KRT42 | Canis lupus familiaris OX=9615 | IF rod domain-containing protein | 37.7 | 26.3 |
| RPS6KA1 | Canis lupus familiaris OX=9615 | Ribosomal protein S6 kinase | 37.6 | 34.8 |
| SHROOM2 | Canis lupus familiaris OX=9615 | Shroom family member 2 | 37.5 | 40.4 |
| SYNRG | Canis lupus familiaris OX=9615 | Synergin gamma | 37.5 | 44.1 |

| | | | | |
|---|---|---|---|---|
| NDUFAF2 | Canis lupus familiaris OX=9615 | NADH:ubiquinone oxidoreductase complex assembly factor 2 | 37.3 | 35.5 |
| CLINT1 | Canis lupus familiaris OX=9615 | Clathrin interactor 1 | 37.3 | 41.1 |
| FRS2 | Canis lupus familiaris OX=9615 | Fibroblast growth factor receptor substrate 2 | 37.2 | 28.5 |
| RAB11B | Canis lupus familiaris OX=9615 | RAB11B, member RAS oncogene family | 37.2 | 37.2 |
| ABLIM3 | Canis lupus familiaris OX=9615 | Actin binding LIM protein family member 3 | 37.1 | 36.1 |
| KRT5 | Canis lupus familiaris OX=9615 | IF rod domain-containing protein | 37.1 | 36.4 |
| AGFG1 | Canis lupus familiaris OX=9615 | ArfGAP with FG repeats 1 | 37.1 | 37.5 |
| DDX3X | Canis lupus familiaris OX=9615 | DEAD-box helicase 3 X-linked | 37 | 34.7 |
| JCAD | Canis lupus familiaris OX=9615 | Junctional cadherin 5 associated | 37 | 38.6 |
| EHD1 | Canis lupus familiaris OX=9615 | EH domain containing 1 | 36.9 | 32.1 |
| ABLIM3 | Canis lupus familiaris OX=9615 | Actin binding LIM protein family member 3 | 36.9 | 36.1 |
| MARK2 | Canis lupus familiaris OX=9615 | Microtubule affinity regulating kinase 2 | 36.7 | 32.6 |
| ARHGAP21 | Canis lupus familiaris OX=9615 | Rho GTPase activating protein 21 | 36.6 | 36.9 |
| HNRNPA1 | Canis lupus familiaris OX=9615 | Uncharacterized protein | 36.5 | 30.3 |
| BCAS1 | Canis lupus familiaris OX=9615 | Uncharacterized protein | 36.5 | 30.9 |
| SIPA1L3 | Canis lupus familiaris OX=9615 | Signal induced proliferation associated 1 like 3 | 36.5 | 38.5 |
| EMD | Canis lupus familiaris OX=9615 | Emerin | 36.5 | 50.4 |
| ARHGAP21 | Canis lupus familiaris OX=9615 | Rho GTPase activating protein 21 | 36.4 | 36.9 |
| KRT10 | Canis lupus familiaris OX=9615 | Keratin, type I cytoskeletal 10 | 36.4 | 38.9 |
| TMSB10 | Canis lupus familiaris OX=9615 | Thymosin beta | 36.4 | 65.9 |
| KRT17 | Canis lupus familiaris OX=9615 | Keratin 17 | 36.3 | 30.9 |
| PKP3 | Canis lupus familiaris OX=9615 | Plakophilin 3 | 36.3 | 32.5 |
| LZIC | Canis lupus familiaris OX=9615 | Leucine zipper and CTNNBIP1 domain containing | 36.3 | 36.3 |
| CRIP2 | Canis lupus familiaris OX=9615 | Cysteine rich protein 2 | 36.1 | 36.1 |
| CAPG | Canis lupus familiaris OX=9615 | Capping actin protein, gelsolin like | 36.1 | 37 |
| EIF2B1 | Canis lupus familiaris OX=9615 | Eukaryotic translation initiation factor 2B subunit alpha | 36.1 | 38.7 |
| C12H6orf132 | Canis lupus familiaris OX=9615 | Chromosome 12 C6orf132 homolog | 36 | 46.6 |
| SERBP1 | Canis lupus familiaris OX=9615 | SERPINE1 mRNA binding protein 1 | 35.6 | 37.3 |
| TMOD3 | Canis lupus familiaris OX=9615 | Tropomodulin 3 | 35.5 | 33 |
| LUZP1 | Canis lupus familiaris OX=9615 | Leucine zipper protein 1 | 35.1 | 38.7 |
| EIF4B | Canis lupus familiaris OX=9615 | Eukaryotic translation initiation factor 4B | 34.6 | 33.8 |
| CHMP4B | Canis lupus familiaris OX=9615 | Charged multivesicular body protein 4B | 34.4 | 34.4 |
| CPLX1 | Canis lupus familiaris OX=9615 | Complexin 1 | 34.3 | 23.9 |
| PRDX2 | Canis lupus familiaris OX=9615 | Peroxiredoxin 2 | 34.3 | 27.3 |
| ARHGDIA | Canis lupus familiaris OX=9615 | Rho GDP dissociation inhibitor alpha | 34.3 | 45.6 |
| C15H1orf50 | Canis lupus familiaris OX=9615 | Chromosome 15 C1orf50 homolog | 34.2 | 26.1 |
| HNRNPK | Canis lupus familiaris OX=9615 | Heterogeneous nuclear ribonucleoprotein K | 34.2 | 27.8 |
| NUDT5 | Canis lupus familiaris OX=9615 | Nudix hydrolase 5 | 34.2 | 34.2 |
| GLO1 | Canis lupus familiaris OX=9615 | Lactoylglutathione lyase | 34.2 | 34.2 |
| RUVBL1 | Canis lupus familiaris OX=9615 | RuvB-like helicase | 34.2 | 36.6 |
| PKP2 | Canis lupus familiaris OX=9615 | Plakophilin 2 | 34 | 33.6 |
| KRT18 | Canis lupus familiaris OX=9615 | IF rod domain-containing protein | 34 | 40 |
| SH3BP4 | Canis lupus familiaris OX=9615 | SH3 domain binding protein 4 | 33.9 | 42 |
| MARK3 | Canis lupus familiaris OX=9615 | Non-specific serine/threonine protein kinase | 33.8 | 30.7 |
| SNRPB | Canis lupus familiaris OX=9615 | Small nuclear ribonucleoprotein-associated protein | 33.8 | 39.8 |
| PICALM | Canis lupus familiaris OX=9615 | Phosphatidylinositol binding clathrin assembly protein | 33.8 | 42.7 |
| CAPZA2 | Canis lupus familiaris OX=9615 | F-actin-capping protein subunit alpha-2 | 33.6 | 27.6 |
| WIPF2 | Canis lupus familiaris OX=9615 | WAS/WASL interacting protein family member 2 | 33.6 | 30.6 |
| PDAP1 | Canis lupus familiaris OX=9615 | PDGFA associated protein 1 | 33.5 | 29.7 |
| ACTC1 | Canis lupus familiaris OX=9615 | Actin alpha cardiac muscle 1 | 33.4 | 29.7 |
| EPN2 | Canis lupus familiaris OX=9615 | Epsin 2 | 33.1 | 24.5 |
| ARFGAP3 | Canis lupus familiaris OX=9615 | ADP ribosylation factor GTPase activating protein 3 | 33.1 | 27.5 |
| KIAA1191 | Canis lupus familiaris OX=9615 | KIAA1191 | 33.1 | 33.4 |
| COBLL1 | Canis lupus familiaris OX=9615 | Cordon-bleu WH2 repeat protein like 1 | 33.1 | 34.1 |
| EIF4G2 | Canis lupus familiaris OX=9615 | Eukaryotic translation initiation factor 4 gamma 2 | 33.1 | 38.1 |
| SEPTIN2 | Canis lupus familiaris OX=9615 | Septin 2 | 32.9 | 32.9 |
| CAST | Canis lupus familiaris OX=9615 | Calpastatin | 32.9 | 33.3 |
| CEP55 | Canis lupus familiaris OX=9615 | Centrosomal protein 55 | 32.9 | 35.7 |
| HIST1H1C | Canis lupus familiaris OX=9615 | H15 domain-containing protein | 32.9 | 38.5 |
| AKAP12 | Canis lupus familiaris OX=9615 | A-kinase anchoring protein 12 | 32.7 | 32.4 |
| MARK2 | Canis lupus familiaris OX=9615 | Non-specific serine/threonine protein kinase | 32.7 | 32.6 |
| ARF6 | Canis lupus familiaris OX=9615 | ADP ribosylation factor 6 | 32.6 | 32.6 |
| CYCS | Canis lupus familiaris OX=9615 | Cytochrome c | 32.4 | 55.2 |
| HEBP1 | Canis lupus familiaris OX=9615 | Heme binding protein 1 | 32.3 | 14.8 |
| HNRNPA2B1 | Canis lupus familiaris OX=9615 | Heterogeneous nuclear ribonucleoprotein A2/B1 | 32.2 | 34.2 |
| LSR | Canis lupus familiaris OX=9615 | Lipolysis stimulated lipoprotein receptor | 32.1 | 33.9 |
| LOC488269 | Canis lupus familiaris OX=9615 | H15 domain-containing protein | 32 | 37.4 |
| TMPO | Canis lupus familiaris OX=9615 | Thymopoietin | 31.9 | 35.2 |
| EPB41L1 | Canis lupus familiaris OX=9615 | Erythrocyte membrane protein band 4.1 like 1 | 31.8 | 26.7 |
| NCK1 | Canis lupus familiaris OX=9615 | Cytoplasmic protein | 31.8 | 47.2 |
| VAMP3 | Canis lupus familiaris OX=9615 | Vesicle associated membrane protein 3 | 31.7 | 15.4 |
| FAM83B | Canis lupus familiaris OX=9615 | Family with sequence similarity 83 member B | 31.6 | 29.1 |
| RAB23 | Canis lupus familiaris OX=9615 | RAB23, member RAS oncogene family | 31.6 | 29.5 |
| DOK1 | Canis lupus familiaris OX=9615 | Docking protein 1 | 31.5 | 31.7 |
| EPB41L1 | Canis lupus familiaris OX=9615 | Erythrocyte membrane protein band 4.1 like 1 | 31.4 | 26.7 |
| CCDC102A | Canis lupus familiaris OX=9615 | Coiled-coil domain containing 102A | 31.4 | 37.4 |

| | | | | |
|---|---|---|---|---|
| RPS17 | Canis lupus familiaris OX=9615 | 40S ribosomal protein S17 | 31.3 | 23.9 |
| NQO1 | Canis lupus familiaris OX=9615 | NAD(P)H quinone dehydrogenase 1 | 31.3 | 33.8 |
| HNRNPA3 | Canis lupus familiaris OX=9615 | Heterogeneous nuclear ribonucleoprotein A3 | 31.2 | 28.6 |
| MAGED2 | Canis lupus familiaris OX=9615 | MAGE family member D2 | 31.1 | 34 |
| USP6NL | Canis lupus familiaris OX=9615 | USP6 N-terminal like | 31 | 26.9 |
| EIF3H | Canis lupus familiaris OX=9615 | Eukaryotic translation initiation factor 3 subunit H | 31 | 39.3 |
| TRIM25 | Canis lupus familiaris OX=9615 | Tripartite motif containing 25 | 30.9 | 29.2 |
| BCAS1 | Canis lupus familiaris OX=9615 | Uncharacterized protein | 30.9 | 30.9 |
| TUBB4B | Canis lupus familiaris OX=9615 | Tubulin beta chain | 30.8 | 29.9 |
| VAMP8 | Canis lupus familiaris OX=9615 | Vesicle associated membrane protein 8 | 30.7 | 21.8 |
| CDC42EP4 | Canis lupus familiaris OX=9615 | CDC42 effector protein 4 | 30.7 | 23.3 |
| CSNK1A1 | Canis lupus familiaris OX=9615 | Protein kinase domain-containing protein | 30.6 | 35.3 |
| MCRIP1 | Canis lupus familiaris OX=9615 | MAPK regulated corepressor interacting protein 1 | 30.5 | 42.1 |
| MAP4 | Canis lupus familiaris OX=9615 | Microtubule-associated protein | 30.4 | 33.8 |
| PRDX6 | Canis lupus familiaris OX=9615 | Peroxiredoxin 6 | 30.3 | 37.6 |
| RPS23 | Canis lupus familiaris OX=9615 | Ribosomal protein S23 | 30.1 | 16.1 |
| EPB41 | Canis lupus familiaris OX=9615 | Protein 4.1 | 30.1 | 30.3 |
| PKM | Canis lupus familiaris OX=9615 | Pyruvate kinase | 30 | 30.5 |
| LOC476006 | Canis lupus familiaris OX=9615 | Uncharacterized protein | 30 | 32.9 |
| COL17A1 | Canis lupus familiaris OX=9615 | Collagen alpha-1(XVII) chain | 29.8 | 28.4 |
| KRT7 | Canis lupus familiaris OX=9615 | IF rod domain-containing protein | 29.7 | 6.4 |
| PPP1R18 | Canis lupus familiaris OX=9615 | Protein phosphatase 1 regulatory subunit 18 | 29.7 | 32.6 |
| TWF1 | Canis lupus familiaris OX=9615 | Twinfilin actin binding protein 1 | 29.7 | 33.7 |
| TAB1 | Canis lupus familiaris OX=9615 | TGF-beta activated kinase 1 (MAP3K7) binding protein 1 | 29.5 | 26.7 |
| SCEL | Canis lupus familiaris OX=9615 | Sciellin | 29.5 | 31.5 |
| SPTBN1 | Canis lupus familiaris OX=9615 | Spectrin beta chain | 29.5 | 35.7 |
| CALM2 | Canis lupus familiaris OX=9615 | Uncharacterized protein | 29.5 | 38.3 |
| CHMP6 | Canis lupus familiaris OX=9615 | Charged multivesicular body protein 6 | 29.4 | 25.6 |
| SH3KBP1 | Canis lupus familiaris OX=9615 | SH3 domain containing kinase binding protein 1 | 29.3 | 29.2 |
| RHEB | Canis lupus familiaris OX=9615 | Ras homolog, mTORC1 binding | 29.3 | 29.3 |
| LOC102153759 | Canis lupus familiaris OX=9615 | Small nuclear ribonucleoprotein E | 29.3 | 29.3 |
| EGFR | Canis lupus familiaris OX=9615 | Receptor protein-tyrosine kinase | 29.2 | 24.2 |
| NECAP2 | Canis lupus familiaris OX=9615 | NECAP endocytosis associated 2 | 29.2 | 29.2 |
| IST1 | Canis lupus familiaris OX=9615 | IST1 factor associated with ESCRT-III | 29 | 20.2 |
| SMG9 | Canis lupus familiaris OX=9615 | SMG9 nonsense mediated mRNA decay factor | 29 | 35.4 |
| FLNB | Canis lupus familiaris OX=9615 | Filamin B | 28.8 | 28.4 |
| SEC22B | Canis lupus familiaris OX=9615 | Uncharacterized protein | 28.7 | 18.2 |
| ANLN | Canis lupus familiaris OX=9615 | Anillin actin binding protein | 28.7 | 34.8 |
| RPS27 | Canis lupus familiaris OX=9615 | 40S ribosomal protein S27 | 28.6 | 29.8 |
| NDRG1 | Canis lupus familiaris OX=9615 | N-Myc downstream regulated gene 1 | 28.6 | 34.4 |
| DAB2 | Canis lupus familiaris OX=9615 | DAB adaptor protein 2 | 28.4 | 33 |
| RPL3 | Canis lupus familiaris OX=9615 | Uncharacterized protein | 28.3 | 17.9 |
| SQSTM1 | Canis lupus familiaris OX=9615 | Sequestosome 1 | 28.3 | 21.8 |
| PARD3 | Canis lupus familiaris OX=9615 | Par-3 family cell polarity regulator | 28.3 | 34.2 |
| TAX1BP3 | Canis lupus familiaris OX=9615 | Tax1-binding protein 3 | 28.2 | 64.5 |
| ARFGAP2 | Canis lupus familiaris OX=9615 | ADP ribosylation factor GTPase activating protein 2 | 28.1 | 14 |
| NUDC | Canis lupus familiaris OX=9615 | Nuclear distribution C, dynein complex regulator | 28 | 24.1 |
| CSRP1 | Canis lupus familiaris OX=9615 | Cysteine and glycine rich protein 1 | 28 | 28 |
| ARL15 | Canis lupus familiaris OX=9615 | ADP ribosylation factor like GTPase 15 | 27.9 | 10.8 |
| EFR3A | Canis lupus familiaris OX=9615 | EFR3 homolog A | 27.9 | 24.6 |
| ZYX | Canis lupus familiaris OX=9615 | Zyxin | 27.9 | 30.9 |
| SORBS2 | Canis lupus familiaris OX=9615 | Sorbin and SH3 domain containing 2 | 27.9 | 41.5 |
| PLEC | Canis lupus familiaris OX=9615 | Plectin | 27.8 | 33 |
| PCBP1 | Canis lupus familiaris OX=9615 | Uncharacterized protein | 27.8 | 35.1 |
| PSMA8 | Canis lupus familiaris OX=9615 | Proteasome subunit alpha type | 27.7 | 13.9 |
| LDHA | Canis lupus familiaris OX=9615 | L-lactate dehydrogenase | 27.7 | 21.9 |
| TES | Canis lupus familiaris OX=9615 | Testin | 27.7 | 35.4 |
| VASP | Canis lupus familiaris OX=9615 | Vasodilator-stimulated phosphoprotein | 27.6 | 26.6 |
| SDCBP | Canis lupus familiaris OX=9615 | Uncharacterized protein | 27.6 | 43.8 |
| COL17A1 | Canis lupus familiaris OX=9615 | Collagen alpha-1(XVII) chain | 27.5 | 28.4 |
| PPL | Canis lupus familiaris OX=9615 | Periplakin | 27.5 | 33.5 |
| DSP | Canis lupus familiaris OX=9615 | Desmoplakin | 27.4 | 21.9 |
| MSN | Canis lupus familiaris OX=9615 | Moesin | 27.4 | 25.3 |
| SLC3A2 | Canis lupus familiaris OX=9615 | Aamy domain-containing protein | 27.4 | 28.7 |
| HIST3H2A | Canis lupus familiaris OX=9615 | Histone H2A | 27.3 | 14.8 |
| PSMB4 | Canis lupus familiaris OX=9615 | Proteasome subunit beta | 27.3 | 15.9 |
| NIBAN2 | Canis lupus familiaris OX=9615 | Family with sequence similarity 129 member B | 27.1 | 26.8 |
| COL17A1 | Canis lupus familiaris OX=9615 | Collagen alpha-1(XVII) chain | 27.1 | 28.4 |
| SFN | Canis lupus familiaris OX=9615 | Stratifin | 27 | 8.9 |
| CSNK1D | Canis lupus familiaris OX=9615 | Casein kinase 1 delta | 27 | 23.4 |
| TUBB | Canis lupus familiaris OX=9615 | Tubulin beta chain | 27 | 30 |
| EDF1 | Canis lupus familiaris OX=9615 | Endothelial differentiation related factor 1 | 27 | 35.8 |
| PBDC1 | Canis lupus familiaris OX=9615 | Polysacc_synt_4 domain-containing protein | 26.8 | 32.7 |
| LOC607207 | Canis lupus familiaris OX=9615 | Uncharacterized protein | 26.7 | 16.3 |
| UTRN | Canis lupus familiaris OX=9615 | Utrophin | 26.7 | 20 |
| ABI1 | Canis lupus familiaris OX=9615 | Abl interactor 1 | 26.7 | 22.7 |

| | | | | |
|---|---|---|---|---|
| NEBL | Canis lupus familiaris OX=9615 | Nebulette | 26.7 | 38.1 |
| KIRREL1 | Canis lupus familiaris OX=9615 | Kirre like nephrin family adhesion molecule 1 | 26.5 | 21.6 |
| S100A11 | Canis lupus familiaris OX=9615 | Protein S100 | 26.5 | 26.5 |
| STK10 | Canis lupus familiaris OX=9615 | Serine/threonine kinase 10 | 26.5 | 31.9 |
| HAUS8 | Canis lupus familiaris OX=9615 | HAUS augmin like complex subunit 8 | 26.3 | 18.4 |
| CCDC85C | Canis lupus familiaris OX=9615 | Coiled-coil domain containing 85C | 26.3 | 19.1 |
| LYPLAL1 | Canis lupus familiaris OX=9615 | Lysophospholipase like 1 | 26.3 | 20.8 |
| KRT1 | Canis lupus familiaris OX=9615 | Keratin, type II cytoskeletal 1 | 26.3 | 25.2 |
| RTRAF | Canis lupus familiaris OX=9615 | Uncharacterized protein | 26.2 | 25.8 |
| NHS | Canis lupus familiaris OX=9615 | NHS actin remodeling regulator | 26.1 | 25 |
| NUDT9 | Canis lupus familiaris OX=9615 | Nudix hydrolase 9 | 26 | 22.3 |
| SCRIB | Canis lupus familiaris OX=9615 | Scribble planar cell polarity protein | 26 | 22.6 |
| WWTR1 | Canis lupus familiaris OX=9615 | WW domain containing transcription regulator 1 | 26 | 25.8 |
| LYPLA2 | Canis lupus familiaris OX=9615 | Lysophospholipase 2 | 26 | 26.8 |
| EIF2A | Canis lupus familiaris OX=9615 | Eukaryotic translation initiation factor 2A | 26 | 27.9 |
| MAP4K4 | Canis lupus familiaris OX=9615 | Mitogen-activated protein kinase kinase kinase kinase 4 | 25.9 | 28.4 |
| CDK1 | Canis lupus familiaris OX=9615 | Cyclin dependent kinase 1 | 25.9 | 35.4 |
| PXN | Canis lupus familiaris OX=9615 | Paxillin | 25.8 | 28.5 |
| HNRNPH3 | Canis lupus familiaris OX=9615 | Heterogeneous nuclear ribonucleoprotein H3 | 25.7 | 20.2 |
| PCBP2 | Canis lupus familiaris OX=9615 | Uncharacterized protein | 25.7 | 23.5 |
| MINK1 | Canis lupus familiaris OX=9615 | Misshapen like kinase 1 | 25.7 | 28.4 |
| CSDE1 | Canis lupus familiaris OX=9615 | Cold shock domain containing E1 | 25.6 | 19.2 |
| TBCE | Canis lupus familiaris OX=9615 | Tubulin folding cofactor E | 25.6 | 26.9 |
| AHCYL1 | Canis lupus familiaris OX=9615 | Adenosylhomocysteinase like 1 | 25.5 | 19.5 |
| MACC1 | Canis lupus familiaris OX=9615 | MET transcriptional regulator MACC1 | 25.5 | 22.9 |
| PATJ | Canis lupus familiaris OX=9615 | PATJ crumbs cell polarity complex component | 25.5 | 30.9 |
| SCAMP1 | Canis lupus familiaris OX=9615 | Secretory carrier-associated membrane protein | 25.4 | 11.5 |
| GSPT1 | Canis lupus familiaris OX=9615 | G1 to S phase transition 1 | 25.3 | 20.7 |
| IDO1 | Canis lupus familiaris OX=9615 | Indoleamine 2,3-dioxygenase 1 | 25.3 | 36.9 |
| EHBP1 | Canis lupus familiaris OX=9615 | EH domain binding protein 1 | 25.1 | 25.7 |
| DDX6 | Canis lupus familiaris OX=9615 | DEAD-box helicase 6 | 25.1 | 28.6 |
| DYNC1LI2 | Canis lupus familiaris OX=9615 | Dynein light intermediate chain | 24.9 | 22 |
| VBP1 | Canis lupus familiaris OX=9615 | Prefoldin subunit 3 | 24.9 | 26.9 |
| LRATD2 | Canis lupus familiaris OX=9615 | Family with sequence similarity 84 member B | 24.9 | 29.9 |
| DLG1 | Canis lupus familiaris OX=9615 | Discs large MAGUK scaffold protein 1 | 24.9 | 34.2 |
| NDE1 | Canis lupus familiaris OX=9615 | NudE neurodevelopment protein 1 | 24.8 | 27.8 |
| PPP1R8 | Canis lupus familiaris OX=9615 | t-SNARE coiled-coil homology domain-containing protein | 24.7 | 12.7 |
| NUP35 | Canis lupus familiaris OX=9615 | Nucleoporin NUP53 | 24.7 | 20.5 |
| ARPC2 | Canis lupus familiaris OX=9615 | Arp2/3 complex 34 kDa subunit | 24.7 | 26.9 |
| NHSL1 | Canis lupus familiaris OX=9615 | NHS like 1 | 24.7 | 29.2 |
| PCNP | Canis lupus familiaris OX=9615 | PEST proteolytic signal containing nuclear protein | 24.7 | 31.6 |
| PLEKHO2 | Canis lupus familiaris OX=9615 | Pleckstrin homology domain containing O2 | 24.5 | 17.3 |
| CFAP298 | Canis lupus familiaris OX=9615 | Uncharacterized protein | 24.5 | 17.9 |
| FAM83H | Canis lupus familiaris OX=9615 | Family with sequence similarity 83 member H | 24.4 | 25.9 |
| VAMP5 | Canis lupus familiaris OX=9615 | Vesicle associated membrane protein 5 | 24.3 | 24.3 |
| RBPMS | Canis lupus familiaris OX=9615 | RNA binding protein, mRNA processing factor | 24.2 | 16 |
| HSPA9 | Canis lupus familiaris OX=9615 | Stress-70 protein, mitochondrial | 24.2 | 16.9 |
| NUMB | Canis lupus familiaris OX=9615 | NUMB, endocytic adaptor protein | 24.1 | 28.8 |
| VPS37A | Canis lupus familiaris OX=9615 | VPS37A subunit of ESCRT-I | 24 | 22.5 |
| SIPA1L1 | Canis lupus familiaris OX=9615 | Signal induced proliferation associated 1 like 1 | 24 | 25.5 |
| ARVCF | Canis lupus familiaris OX=9615 | ARVCF, delta catenin family member | 24 | 30.1 |
| RRM2 | Canis lupus familiaris OX=9615 | Ribonucleotide reductase regulatory subunit M2 | 23.9 | 20.7 |
| GOLGA5 | Canis lupus familiaris OX=9615 | Golgin A5 | 23.7 | 32.6 |
| PPP1CA | Canis lupus familiaris OX=9615 | Serine/threonine-protein phosphatase PP1-alpha catalytic subunit | 23.6 | 23.9 |
| SHC1 | Canis lupus familiaris OX=9615 | SHC adaptor protein 1 | 23.4 | 12 |
| ARCN1 | Canis lupus familiaris OX=9615 | Coatomer subunit delta | 23.3 | 21.7 |
| PDCL3 | Canis lupus familiaris OX=9615 | Phosducin like 3 | 23.3 | 24.2 |
| PYM1 | Canis lupus familiaris OX=9615 | PYM homolog 1, exon junction complex associated factor | 23.2 | 16.7 |
| TNS1 | Canis lupus familiaris OX=9615 | Tensin 1 | 23.2 | 25.4 |
| TRIP6 | Canis lupus familiaris OX=9615 | Thyroid hormone receptor interactor 6 | 23.1 | 35.5 |
| CCSER1 | Canis lupus familiaris OX=9615 | Coiled-coil serine rich protein 1 | 23 | 13.7 |
| RPL15 | Canis lupus familiaris OX=9615 | 60S ribosomal protein L15 | 23 | 19.1 |
| MICALL1 | Canis lupus familiaris OX=9615 | MICAL like 1 | 23 | 28.1 |
| LOC483167 | Canis lupus familiaris OX=9615 | Histone H4 | 22.9 | 34.4 |
| RACK1 | Canis lupus familiaris OX=9615 | Receptor for activated C kinase 1 | 22.7 | 20.5 |
| EHBP1L1 | Canis lupus familiaris OX=9615 | EH domain binding protein 1 like 1 | 22.7 | 27.6 |
| VAPB | Canis lupus familiaris OX=9615 | VAMP associated protein B and C | 22.6 | 14.8 |
| TMEM263 | Canis lupus familiaris OX=9615 | Transmembrane protein 263 | 22.6 | 22.6 |
| PRAG1 | Canis lupus familiaris OX=9615 | PEAK1 related, kinase-activating pseudokinase 1 | 22.6 | 23.1 |
| PDXDC1 | Canis lupus familiaris OX=9615 | Uncharacterized protein | 22.6 | 25 |
| UBAP2L | Canis lupus familiaris OX=9615 | Ubiquitin associated protein 2 like | 22.5 | 24.6 |
| RAB11FIP1 | Canis lupus familiaris OX=9615 | Uncharacterized protein | 22.4 | 19.1 |
| MAPT | Canis lupus familiaris OX=9615 | Microtubule-associated protein | 22.3 | 24.7 |
| SLC4A7 | Canis lupus familiaris OX=9615 | Anion exchange protein | 22.2 | 16 |
| PLEKHN1 | Canis lupus familiaris OX=9615 | Pleckstrin homology domain containing N1 | 22.2 | 22 |
| ERBB2 | Canis lupus familiaris OX=9615 | Receptor protein-tyrosine kinase | 22.2 | 23.3 |

| | | | | |
|---|---|---|---|---|
| SNRPD2 | Canis lupus familiaris OX=9615 | Small nuclear ribonucleoprotein Sm D2 | 22.2 | 24.8 |
| GLTP | Canis lupus familiaris OX=9615 | ANK_REP_REGION domain-containing protein | 22.2 | 25.4 |
| EIF5A | Canis lupus familiaris OX=9615 | Eukaryotic translation initiation factor 5A | 22.1 | 23.4 |
| TNRC6A | Canis lupus familiaris OX=9615 | Trinucleotide repeat containing adaptor 6A | 22 | 24.8 |
| CCT2 | Canis lupus familiaris OX=9615 | Chaperonin containing TCP1 subunit 2 | 21.9 | 20.7 |
| UACA | Canis lupus familiaris OX=9615 | Uveal autoantigen with coiled-coil domains and ankyrin repeats | 21.9 | 23.5 |
| FGD6 | Canis lupus familiaris OX=9615 | FYVE, RhoGEF and PH domain containing 6 | 21.9 | 25.9 |
| ARHGAP12 | Canis lupus familiaris OX=9615 | Rho GTPase activating protein 12 | 21.9 | 28.8 |
| STX7 | Canis lupus familiaris OX=9615 | Syntaxin 7 | 21.8 | 17.2 |
| CDC37 | Canis lupus familiaris OX=9615 | Cell division cycle 37 | 21.8 | 29.7 |
| BAIAP2 | Canis lupus familiaris OX=9615 | BAI1 associated protein 2 | 21.8 | 35.7 |
| LLGL1 | Canis lupus familiaris OX=9615 | LLGL scribble cell polarity complex component 1 | 21.7 | 19.8 |
| FERMT2 | Canis lupus familiaris OX=9615 | Fermitin family member 2 | 21.7 | 21 |
| UFD1 | Canis lupus familiaris OX=9615 | Ubiquitin recognition factor in ER associated degradation 1 | 21.5 | 17.9 |
| SF1 | Canis lupus familiaris OX=9615 | Splicing factor 1 | 21.5 | 19.3 |
| EIF4H | Canis lupus familiaris OX=9615 | Eukaryotic translation initiation factor 4H | 21.5 | 27.1 |
| LIMD1 | Canis lupus familiaris OX=9615 | LIM domains containing 1 | 21.5 | 28.1 |
| GEN1 | Canis lupus familiaris OX=9615 | GEN1 Holliday junction 5' flap endonuclease | 21.4 | 18.4 |
| OTUD7B | Canis lupus familiaris OX=9615 | OTU deubiquitinase 7B | 21.4 | 20.3 |
| FAM110A | Canis lupus familiaris OX=9615 | Family with sequence similarity 110 member A | 21.4 | 26.8 |
| SOD1 | Canis lupus familiaris OX=9615 | Superoxide dismutase [Cu-Zn] | 21.3 | 30.1 |
| SMAP2 | Canis lupus familiaris OX=9615 | Small ArfGAP2 | 21.2 | 26.4 |
| CTNND2 | Canis lupus familiaris OX=9615 | Catenin delta 2 | 21 | 17.6 |
| G6PD | Canis lupus familiaris OX=9615 | Glucose-6-phosphate 1-dehydrogenase | 21 | 36.7 |
| PLCH1 | Canis lupus familiaris OX=9615 | Phosphoinositide phospholipase C | 20.9 | 24 |
| PGAM1 | Canis lupus familiaris OX=9615 | Phosphoglycerate mutase | 20.9 | 29.9 |
| PFDN6 | Canis lupus familiaris OX=9615 | Prefoldin subunit 6 | 20.9 | 42.6 |
| CISD2 | Canis lupus familiaris OX=9615 | CDGSH iron sulfur domain 2 | 20.7 | 20 |
| CSNK1E | Canis lupus familiaris OX=9615 | Casein kinase 1 epsilon | 20.7 | 20.7 |
| RAB21 | Canis lupus familiaris OX=9615 | Ras-related protein Rab-21 | 20.6 | 12.1 |
| RPL21 | Canis lupus familiaris OX=9615 | Uncharacterized protein | 20.6 | 20.6 |
| PEAK1 | Canis lupus familiaris OX=9615 | Pseudopodium enriched atypical kinase 1 | 20.6 | 22.8 |
| TP53BP2 | Canis lupus familiaris OX=9615 | Tumor protein p53 binding protein 2 | 20.6 | 25.1 |
| STAMBPL1 | Canis lupus familiaris OX=9615 | STAM binding protein like 1 | 20.6 | 31 |
| SH3PXD2B | Canis lupus familiaris OX=9615 | SH3 and PX domains 2B | 20.3 | 23.3 |
| STOM | Canis lupus familiaris OX=9615 | Stomatin | 20.2 | 6.9 |
| CFAP36 | Canis lupus familiaris OX=9615 | Cilia and flagella associated protein 36 | 20.2 | 24.6 |
| SNX2 | Canis lupus familiaris OX=9615 | Sorting nexin 2 | 20.1 | 13.2 |
| KRT75 | Canis lupus familiaris OX=9615 | Keratin 75 | 20.1 | 20.3 |
| CYTH2 | Canis lupus familiaris OX=9615 | Cytohesin 2 | 20.1 | 20.4 |
| STMN1 | Canis lupus familiaris OX=9615 | Stathmin | 20.1 | 20.8 |
| KANK2 | Canis lupus familiaris OX=9615 | KN motif and ankyrin repeat domains 2 | 20 | 24.4 |
| RPL29 | Canis lupus familiaris OX=9615 | Uncharacterized protein | 19.9 | 12.7 |
| LGALS3 | Canis lupus familiaris OX=9615 | Galectin-3 | 19.9 | 20.3 |
| PPP1R11 | Canis lupus familiaris OX=9615 | Protein phosphatase 1 regulatory inhibitor subunit 11 | 19.8 | 19.8 |
| PDCD5 | Canis lupus familiaris OX=9615 | Programmed cell death 5 | 19.8 | 21.6 |
| CCT4 | Canis lupus familiaris OX=9615 | T-complex protein 1 subunit delta | 19.7 | 18.9 |
| RPL18 | Canis lupus familiaris OX=9615 | Ribosomal protein L18 | 19.7 | 19.7 |
| ARHGAP29 | Canis lupus familiaris OX=9615 | Rho GTPase activating protein 29 | 19.7 | 22.2 |
| ARFIP1 | Canis lupus familiaris OX=9615 | ADP ribosylation factor interacting protein 1 | 19.7 | 25.9 |
| YTHDF3 | Canis lupus familiaris OX=9615 | YTH N6-methyladenosine RNA binding protein 3 | 19.7 | 26.3 |
| FKBP3 | Canis lupus familiaris OX=9615 | Peptidylprolyl isomerase | 19.6 | 14.7 |
| PPP1CB | Canis lupus familiaris OX=9615 | Serine/threonine-protein phosphatase PP1-beta catalytic subunit | 19.6 | 15 |
| SPATA2 | Canis lupus familiaris OX=9615 | Spermatogenesis associated 2 | 19.6 | 17.7 |
| PLEKHA4 | Canis lupus familiaris OX=9615 | Pleckstrin homology domain containing A4 | 19.6 | 25.1 |
| PSME3 | Canis lupus familiaris OX=9615 | Proteasome activator subunit 3 | 19.5 | 19.1 |
| TULP3 | Canis lupus familiaris OX=9615 | Tubby-like protein | 19.5 | 22.9 |
| RPS4X | Canis lupus familiaris OX=9615 | 40S ribosomal protein S4 | 19.4 | 19.8 |
| MAPRE1 | Canis lupus familiaris OX=9615 | Uncharacterized protein | 19.3 | 19.3 |
| NAV2 | Canis lupus familiaris OX=9615 | Neuron navigator 2 | 19.3 | 22.3 |
| SNX9 | Canis lupus familiaris OX=9615 | Sorting nexin | 19.3 | 24.5 |
| ANK3 | Canis lupus familiaris OX=9615 | Ankyrin 3 | 19.2 | 20.4 |
| HDLBP | Canis lupus familiaris OX=9615 | High density lipoprotein binding protein | 19.2 | 22.7 |
| RPL11 | Canis lupus familiaris OX=9615 | Ribosomal protein L11 | 19.1 | 6.7 |
| LLGL2 | Canis lupus familiaris OX=9615 | LLGL scribble cell polarity complex component 2 | 19 | 19.7 |
| RPL6 | Canis lupus familiaris OX=9615 | 60S ribosomal protein L6 | 18.8 | 24 |
| ZDHHC8 | Canis lupus familiaris OX=9615 | Probable palmitoyltransferase ZDHHC8 | 18.6 | 16.3 |
| LMNA | Canis lupus familiaris OX=9615 | Lamin A/C | 18.6 | 24.2 |
| CHORDC1 | Canis lupus familiaris OX=9615 | Cysteine and histidine rich domain containing 1 | 18.5 | 14.2 |
| STX4 | Canis lupus familiaris OX=9615 | Syntaxin 4 | 18.5 | 20.9 |
| SORBS1 | Canis lupus familiaris OX=9615 | Sorbin and SH3 domain containing 1 | 18.5 | 23 |
| LZTS2 | Canis lupus familiaris OX=9615 | Leucine zipper tumor suppressor 2 | 18.5 | 24.6 |
| RAB7A | Canis lupus familiaris OX=9615 | Ras-related protein Rab-7a | 18.4 | 11.6 |
| EPN3 | Canis lupus familiaris OX=9615 | Epsin 3 | 18.4 | 17.1 |
| TUFM | Canis lupus familiaris OX=9615 | Elongation factor Tu | 18.4 | 17.7 |
| YWHAZ | Canis lupus familiaris OX=9615 | 14_3_3 domain-containing protein | 18.4 | 22.4 |

| | | | | |
|---|---|---|---|---|
| TANK | Canis lupus familiaris OX=9615 | TRAF family member associated NFKB activator | 18.4 | 23.9 |
| RIDA | Canis lupus familiaris OX=9615 | Reactive intermediate imine deaminase A homolog | 18.3 | 18.3 |
| SWAP70 | Canis lupus familiaris OX=9615 | Switching B cell complex subunit SWAP70 | 18.3 | 18.6 |
| EXO5 | Canis lupus familiaris OX=9615 | Exonuclease 5 | 18.3 | 22.5 |
| CDC42EP3 | Canis lupus familiaris OX=9615 | CDC42 effector protein 3 | 18.2 | 19 |
| MEAK7 | Canis lupus familiaris OX=9615 | MTOR associated protein, eak-7 homolog | 18.2 | 19.8 |
| LOC10085663 | Canis lupus familiaris OX=9615 | PNP_UDP_1 domain-containing protein | 18.1 | 15.2 |
| PHLDB2 | Canis lupus familiaris OX=9615 | Pleckstrin homology like domain family B member 2 | 18.1 | 19.7 |
| MAGI1 | Canis lupus familiaris OX=9615 | Uncharacterized protein | 18 | 17.6 |
| EXOC3 | Canis lupus familiaris OX=9615 | Uncharacterized protein | 18 | 23.9 |
| CAP1 | Canis lupus familiaris OX=9615 | Adenylyl cyclase-associated protein | 17.9 | 13.7 |
| UFM1 | Canis lupus familiaris OX=9615 | Ubiquitin fold modifier 1 | 17.9 | 17.9 |
| CHMP5 | Canis lupus familiaris OX=9615 | Uncharacterized protein | 17.8 | 17.8 |
| RPS11 | Canis lupus familiaris OX=9615 | 40S ribosomal protein S11 | 17.7 | 10.8 |
| SLC12A6 | Canis lupus familiaris OX=9615 | Solute carrier family 12 member 6 | 17.7 | 17.8 |
| ATP6V1A | Canis lupus familiaris OX=9615 | ATPase H+ transporting V1 subunit A | 17.6 | 13.9 |
| DDX19A | Canis lupus familiaris OX=9615 | DEAD-box helicase 19A | 17.6 | 25.9 |
| MAGI1 | Canis lupus familiaris OX=9615 | Uncharacterized protein | 17.5 | 17.6 |
| KIAA1671 | Canis lupus familiaris OX=9615 | KIAA1671 | 17.5 | 22.8 |
| LMCD1 | Canis lupus familiaris OX=9615 | LIM and cysteine rich domains 1 | 17.4 | 16.3 |
| ABLIM2 | Canis lupus familiaris OX=9615 | Actin binding LIM protein family member 2 | 17.3 | 14.3 |
| ACACA | Canis lupus familiaris OX=9615 | Acetyl-CoA carboxylase alpha | 17.3 | 14.5 |
| GAK | Canis lupus familiaris OX=9615 | Cyclin G associated kinase | 17.3 | 16.7 |
| USP43 | Canis lupus familiaris OX=9615 | Ubiquitin specific peptidase 43 | 17.3 | 21.6 |
| SYAP1 | Canis lupus familiaris OX=9615 | Synapse associated protein 1 | 17.3 | 25.1 |
| REPS1 | Canis lupus familiaris OX=9615 | RALBP1 associated Eps domain containing 1 | 17.3 | 25.2 |
| VCL | Canis lupus familiaris OX=9615 | Vinculin | 17.2 | 20.1 |
| STAT1 | Canis lupus familiaris OX=9615 | Signal transducer and activator of transcription | 17.1 | 14.9 |
| GAB1 | Canis lupus familiaris OX=9615 | GRB2 associated binding protein 1 | 17 | 17.3 |
| PATL1 | Canis lupus familiaris OX=9615 | PAT1 homolog 1, processing body mRNA decay factor | 16.9 | 15 |
| CFL1 | Canis lupus familiaris OX=9615 | Cofilin 1 | 16.9 | 16.9 |
| HMGN2 | Canis lupus familiaris OX=9615 | Non-histone chromosomal protein HMG-17 | 16.9 | 16.9 |
| SCYL2 | Canis lupus familiaris OX=9615 | SCY1 like pseudokinase 2 | 16.9 | 17.3 |
| DENND4C | Canis lupus familiaris OX=9615 | DENN domain containing 4C | 16.9 | 18 |
| H3F3B | Canis lupus familiaris OX=9615 | Histone H3 | 16.9 | 19.9 |
| FRMD4B | Canis lupus familiaris OX=9615 | FERM domain containing 4B | 16.9 | 20 |
| PCTP | Canis lupus familiaris OX=9615 | Phosphatidylcholine transfer protein | 16.8 | 13.6 |
| DBT | Canis lupus familiaris OX=9615 | Dihydrolipoamide acetyltransferase component of pyruvate dehydrogenase complex | 16.8 | 23.7 |
| GSTP1 | Canis lupus familiaris OX=9615 | Uncharacterized protein | 16.7 | 12.9 |
| RAPH1 | Canis lupus familiaris OX=9615 | Uncharacterized protein | 16.7 | 16.3 |
| XAB2 | Canis lupus familiaris OX=9615 | XPA binding protein 2 | 16.7 | 21.2 |
| PAK4 | Canis lupus familiaris OX=9615 | Non-specific serine/threonine protein kinase | 16.7 | 23.6 |
| ARPC5 | Canis lupus familiaris OX=9615 | Actin-related protein 2/3 complex subunit 5 | 16.7 | 29.8 |
| STIM1 | Canis lupus familiaris OX=9615 | Stromal interaction molecule 1 | 16.6 | 16.8 |
| PIK3C2A | Canis lupus familiaris OX=9615 | Phosphatidylinositol-4-phosphate 3-kinase catalytic subunit type 2 alpha | 16.6 | 19.1 |
| FAM107B | Canis lupus familiaris OX=9615 | Family with sequence similarity 107 member B | 16.6 | 19.9 |
| NONO | Canis lupus familiaris OX=9615 | Non-POU domain containing octamer binding | 16.6 | 32.6 |
| SHROOM4 | Canis lupus familiaris OX=9615 | Shroom family member 4 | 16.4 | 24.1 |
| ATP2B1 | Canis lupus familiaris OX=9615 | Calcium-transporting ATPase | 16.3 | 15.4 |
| ECHDC1 | Canis lupus familiaris OX=9615 | Ethylmalonyl-CoA decarboxylase 1 | 16.3 | 16.3 |
| CAVIN3 | Canis lupus familiaris OX=9615 | Caveolae associated protein 3 | 16.3 | 31.2 |
| WWOX | Canis lupus familiaris OX=9615 | WW domain containing oxidoreductase | 16.2 | 12.6 |
| C5H11orf52 | Canis lupus familiaris OX=9615 | Chromosome 5 C11orf52 homolog | 16.2 | 16.2 |
| EPS15 | Canis lupus familiaris OX=9615 | Epidermal growth factor receptor pathway substrate 15 | 16.2 | 23.9 |
| PFDN2 | Canis lupus familiaris OX=9615 | Prefoldin subunit 2 | 16.2 | 29.2 |
| TPI1 | Canis lupus familiaris OX=9615 | Triosephosphate isomerase | 16.1 | 10 |
| NUP214 | Canis lupus familiaris OX=9615 | Nucleoporin 214 | 16.1 | 15 |
| RAB1A | Canis lupus familiaris OX=9615 | Ras-related protein Rab-1A | 16.1 | 16.1 |
| TNK1 | Canis lupus familiaris OX=9615 | Tyrosine kinase non receptor 1 | 16.1 | 17.5 |
| AIF1L | Canis lupus familiaris OX=9615 | Allograft inflammatory factor 1 like | 16 | 10 |
| TANC2 | Canis lupus familiaris OX=9615 | Tetratricopeptide repeat, ankyrin repeat and coiled-coil containing 2 | 16 | 15.3 |
| ICA1 | Canis lupus familiaris OX=9615 | Islet cell autoantigen 1 | 16 | 15.8 |
| NOS1AP | Canis lupus familiaris OX=9615 | PID domain-containing protein | 16 | 15.9 |
| RPS2 | Canis lupus familiaris OX=9615 | S5 DRBM domain-containing protein | 16 | 16 |
| RELL1 | Canis lupus familiaris OX=9615 | RELT like 1 | 15.9 | 11.4 |
| DBN1 | Canis lupus familiaris OX=9615 | Drebrin 1 | 15.9 | 12.2 |
| TARDBP | Canis lupus familiaris OX=9615 | TAR DNA binding protein | 15.9 | 12.3 |
| APBB2 | Canis lupus familiaris OX=9615 | Amyloid beta precursor protein binding family B member 2 | 15.9 | 17 |
| MICAL3 | Canis lupus familiaris OX=9615 | Microtubule associated monooxygenase, calponin and LIM domain containing 3 | 15.9 | 20.1 |
| MSRB3 | Canis lupus familiaris OX=9615 | Peptide-methionine (R)-S-oxide reductase | 15.8 | 23 |
| SRSF11 | Canis lupus familiaris OX=9615 | Serine and arginine rich splicing factor 11 | 15.7 | 11.3 |
| KRT80 | Canis lupus familiaris OX=9615 | Keratin 80 | 15.7 | 12.1 |
| LBR | Canis lupus familiaris OX=9615 | Lamin B receptor | 15.7 | 12.5 |
| TPT1 | Canis lupus familiaris OX=9615 | TCTP domain-containing protein | 15.7 | 15.7 |
| PLEKHG3 | Canis lupus familiaris OX=9615 | Pleckstrin homology and RhoGEF domain containing G3 | 15.7 | 16.1 |
| SEC61B | Canis lupus familiaris OX=9615 | Protein transport protein Sec61 subunit beta | 15.6 | 15.6 |

| | | | | |
|---|---|---|---|---|
| COG1 | Canis lupus familiaris OX=9615 | Component of oligomeric golgi complex 1 | 15.5 | 14 |
| RNF214 | Canis lupus familiaris OX=9615 | Ring finger protein 214 | 15.5 | 30.3 |
| EIF3J | Canis lupus familiaris OX=9615 | Eukaryotic translation initiation factor 3 subunit J | 15.4 | 15.4 |
| TAF15 | Canis lupus familiaris OX=9615 | TATA-box binding protein associated factor 15 | 15.4 | 32.6 |
| NANP | Canis lupus familiaris OX=9615 | Uncharacterized protein | 15.3 | 16.5 |
| YES1 | Canis lupus familiaris OX=9615 | Tyrosine-protein kinase | 15.3 | 17.7 |
| EIF4A1 | Canis lupus familiaris OX=9615 | Uncharacterized protein | 15.3 | 25.4 |
| EIF4G1 | Canis lupus familiaris OX=9615 | Eukaryotic translation initiation factor 4 gamma 1 | 15.2 | 12.5 |
| PPP1CC | Canis lupus familiaris OX=9615 | Serine/threonine-protein phosphatase PP1-gamma catalytic subunit | 15.2 | 15.5 |
| UTRN | Canis lupus familiaris OX=9615 | Utrophin | 15.2 | 20 |
| ARPC1B | Canis lupus familiaris OX=9615 | Actin-related protein 2/3 complex subunit | 15.1 | 13.2 |
| TIPRL | Canis lupus familiaris OX=9615 | TOR signaling pathway regulator | 15.1 | 15.1 |
| RPL9 | Canis lupus familiaris OX=9615 | Ribosomal protein L9 | 15.1 | 17.2 |
| KIAA0930 | Canis lupus familiaris OX=9615 | KIAA0930 | 15 | 6.2 |
| HSPB1 | Canis lupus familiaris OX=9615 | Heat shock protein beta-1 | 15 | 10.2 |
| VDAC2 | Canis lupus familiaris OX=9615 | Voltage dependent anion channel 2 | 15 | 15 |
| SLC30A1 | Canis lupus familiaris OX=9615 | Solute carrier family 30 member 1 | 15 | 15 |
| CAMK2D | Canis lupus familiaris OX=9615 | Calcium/calmodulin dependent protein kinase II delta | 15 | 15 |
| ARHGAP31 | Canis lupus familiaris OX=9615 | Rho GTPase activating protein 31 | 14.9 | 10.1 |
| YTHDF1 | Canis lupus familiaris OX=9615 | YTH N6-methyladenosine RNA binding protein 1 | 14.9 | 10.2 |
| STX5 | Canis lupus familiaris OX=9615 | Syntaxin 5 | 14.9 | 13 |
| RPL14 | Canis lupus familiaris OX=9615 | Ribosomal_L14e domain-containing protein | 14.8 | 9.5 |
| CAV2 | Canis lupus familiaris OX=9615 | Isoform Beta of Caveolin-2 | 14.8 | 14.8 |
| GAB2 | Canis lupus familiaris OX=9615 | PH domain-containing protein | 14.8 | 16.2 |
| EPS8 | Canis lupus familiaris OX=9615 | Epidermal growth factor receptor pathway substrate 8 | 14.8 | 20.1 |
| MED15 | Canis lupus familiaris OX=9615 | Mediator of RNA polymerase II transcription subunit 15 | 14.8 | 26.4 |
| SUMO2 | Canis lupus familiaris OX=9615 | Small ubiquitin-related modifier | 14.7 | 14.7 |
| CNOT8 | Canis lupus familiaris OX=9615 | CCR4-NOT transcription complex subunit 8 | 14.7 | 19.2 |
| MARVELD2 | Canis lupus familiaris OX=9615 | MARVEL domain containing 2 | 14.6 | 13 |
| EEF2 | Canis lupus familiaris OX=9615 | Eukaryotic translation elongation factor 2 | 14.6 | 17 |
| ARFGAP1 | Canis lupus familiaris OX=9615 | ADP ribosylation factor GTPase activating protein 1 | 14.6 | 20.9 |
| TBC1D5 | Canis lupus familiaris OX=9615 | TBC1 domain family member 5 | 14.6 | 21.6 |
| ARHGAP1 | Canis lupus familiaris OX=9615 | Cdc42 GTPase-activating protein | 14.6 | 25.7 |
| PPME1 | Canis lupus familiaris OX=9615 | Protein phosphatase methylesterase 1 | 14.5 | 11.8 |
| PPP3CA | Canis lupus familiaris OX=9615 | Serine/threonine-protein phosphatase | 14.5 | 19.6 |
| PTTG1 | Canis lupus familiaris OX=9615 | Uncharacterized protein | 14.4 | 8 |
| FAM114A2 | Canis lupus familiaris OX=9615 | Family with sequence similarity 114 member A2 | 14.4 | 8.2 |
| KIF23 | Canis lupus familiaris OX=9615 | Kinesin-like protein | 14.4 | 9.5 |
| HSP90AB1 | Canis lupus familiaris OX=9615 | HATPase_c domain-containing protein | 14.4 | 12.7 |
| GPRIN1 | Canis lupus familiaris OX=9615 | G protein regulated inducer of neurite outgrowth 1 | 14.4 | 22 |
| MOB3B | Canis lupus familiaris OX=9615 | MOB kinase activator 3B | 14.4 | 22.2 |
| PRUNE1 | Canis lupus familiaris OX=9615 | Prune exopolyphosphatase 1 | 14.3 | 17 |
| CCT6A | Canis lupus familiaris OX=9615 | Chaperonin containing TCP1 subunit 6A | 14.2 | 11.9 |
| CETN2 | Canis lupus familiaris OX=9615 | Uncharacterized protein | 14.2 | 14.2 |
| KTN1 | Canis lupus familiaris OX=9615 | Kinectin 1 | 14.2 | 16.7 |
| HSPH1 | Canis lupus familiaris OX=9615 | Heat shock protein family H (Hsp110) member 1 | 14.2 | 17.2 |
| SPC25 | Canis lupus familiaris OX=9615 | SPC25 component of NDC80 kinetochore complex | 14.2 | 24.8 |
| FAM83F | Canis lupus familiaris OX=9615 | Family with sequence similarity 83 member F | 14.2 | 30.5 |
| BLMH | Canis lupus familiaris OX=9615 | Bleomycin hydrolase | 14.1 | 12.3 |
| NUDCD2 | Canis lupus familiaris OX=9615 | NudC domain containing 2 | 14.1 | 13.5 |
| SPART | Canis lupus familiaris OX=9615 | Spartin | 14.1 | 14.4 |
| PHYHIPL | Canis lupus familiaris OX=9615 | Phytanoyl-CoA 2-hydroxylase interacting protein like | 14.1 | 16 |
| RPL8 | Canis lupus familiaris OX=9615 | Ribosomal protein L8 | 14 | 6.6 |
| ASAP2 | Canis lupus familiaris OX=9615 | ArfGAP with SH3 domain, ankyrin repeat and PH domain 2 | 14 | 14 |
| PCMT1 | Canis lupus familiaris OX=9615 | Protein-L-isoaspartate O-methyltransferase | 14 | 14.3 |
| HNRNPA0 | Canis lupus familiaris OX=9615 | Heterogeneous nuclear ribonucleoprotein A0 | 13.9 | 6.5 |
| DENND2C | Canis lupus familiaris OX=9615 | DENN domain containing 2C | 13.9 | 7.4 |
| TAB3 | Canis lupus familiaris OX=9615 | TGF-beta activated kinase 1 (MAP3K7) binding protein 3 | 13.9 | 17.7 |
| KRT12 | Canis lupus familiaris OX=9615 | Keratin 12 | 13.8 | 9.6 |
| RNF121 | Canis lupus familiaris OX=9615 | Ring finger protein 121 | 13.8 | 13.8 |
| MYZAP | Canis lupus familiaris OX=9615 | Uncharacterized protein | 13.8 | 14 |
| ATP2B4 | Canis lupus familiaris OX=9615 | Calcium-transporting ATPase | 13.8 | 14.1 |
| TCP1 | Canis lupus familiaris OX=9615 | T-complex 1 | 13.8 | 14.7 |
| ETF1 | Canis lupus familiaris OX=9615 | eRF1_1 domain-containing protein | 13.8 | 16.1 |
| SH3BP2 | Canis lupus familiaris OX=9615 | SH3 domain binding protein 2 | 13.8 | 18.4 |
| PSMB5 | Canis lupus familiaris OX=9615 | Proteasome subunit beta | 13.7 | 10.6 |
| CIAPIN1 | Canis lupus familiaris OX=9615 | Anamorsin | 13.7 | 13.7 |
| CEP41 | Canis lupus familiaris OX=9615 | Centrosomal protein 41 | 13.6 | 13.6 |
| CHMP2B | Canis lupus familiaris OX=9615 | Charged multivesicular body protein 2B | 13.6 | 17.8 |
| GOLGA2 | Canis lupus familiaris OX=9615 | Uncharacterized protein | 13.5 | 14.2 |
| GRK2 | Canis lupus familiaris OX=9615 | G protein-coupled receptor kinase | 13.4 | 8 |
| NAV1 | Canis lupus familiaris OX=9615 | Neuron navigator 1 | 13.4 | 13.7 |
| SEPTIN11 | Canis lupus familiaris OX=9615 | Septin 11 | 13.4 | 14.5 |
| TIMP3 | Canis lupus familiaris OX=9615 | TIMP metallopeptidase inhibitor 3 | 13.3 | 5.7 |
| GLOD4 | Canis lupus familiaris OX=9615 | Glyoxalase domain containing 4 | 13.3 | 10.8 |
| WASF2 | Canis lupus familiaris OX=9615 | WAS protein family member 2 | 13.3 | 13.3 |

| | | | | |
|---|---|---|---|---|
| GOLGA4 | Canis lupus familiaris OX=9615 | Golgin A4 | 13.3 | 19.3 |
| BIRC4 | Canis lupus familiaris OX=9615 | Baculoviral IAP-repeat containing protein 4 | 13.2 | 14.6 |
| SHB | Canis lupus familiaris OX=9615 | SH2 domain containing adaptor protein B | 13.2 | 17.9 |
| SRPRA | Canis lupus familiaris OX=9615 | Signal recognition particle receptor subunit alpha | 13.2 | 18.1 |
| VANGL1 | Canis lupus familiaris OX=9615 | Vang-like protein | 13.2 | 18.1 |
| FN1 | Canis lupus familiaris OX=9615 | Fibronectin | 13.1 | 9.4 |
| DCP1A | Canis lupus familiaris OX=9615 | Decapping mRNA 1A | 13.1 | 13.1 |
| SHTN1 | Canis lupus familiaris OX=9615 | Shootin 1 | 13.1 | 15.1 |
| SNX27 | Canis lupus familiaris OX=9615 | Sorting nexin family member 27 | 13.1 | 17.7 |
| RBBP6 | Canis lupus familiaris OX=9615 | RB binding protein 6, ubiquitin ligase | 13.1 | 18.6 |
| TBCB | Canis lupus familiaris OX=9615 | Tubulin folding cofactor B | 13.1 | 25.3 |
| ARG1 | Canis lupus familiaris OX=9615 | Arginase | 13 | 9 |
| CGN | Canis lupus familiaris OX=9615 | Cingulin | 13 | 9.5 |
| MRPS31 | Canis lupus familiaris OX=9615 | Mitochondrial ribosomal protein S31 | 13 | 17.6 |
| PAICS | Canis lupus familiaris OX=9615 | AIRC domain-containing protein | 12.9 | 10.4 |
| DCBLD2 | Canis lupus familiaris OX=9615 | Discoidin, CUB and LCCL domain containing 2 | 12.9 | 18.1 |
| EBAG9 | Canis lupus familiaris OX=9615 | Receptor-binding cancer antigen expressed on SiSo cells | 12.9 | 33.9 |
| CPNE3 | Canis lupus familiaris OX=9615 | Copine 3 | 12.8 | 7.3 |
| CACUL1 | Canis lupus familiaris OX=9615 | CDK2 associated cullin domain 1 | 12.8 | 9.7 |
| CDC37L1 | Canis lupus familiaris OX=9615 | Cell division cycle 37 like 1 | 12.8 | 10.4 |
| CRYBG3 | Canis lupus familiaris OX=9615 | Uncharacterized protein | 12.8 | 10.8 |
| TLN1 | Canis lupus familiaris OX=9615 | Talin 1 | 12.8 | 12 |
| GOPC | Canis lupus familiaris OX=9615 | Golgi associated PDZ and coiled-coil motif containing | 12.8 | 12.1 |
| AKAP2 | Canis lupus familiaris OX=9615 | AKAP2_C domain-containing protein | 12.8 | 13.5 |
| SMAP1 | Canis lupus familiaris OX=9615 | SMAP1 protein | 12.8 | 20.4 |
| DENR | Canis lupus familiaris OX=9615 | Density-regulated protein | 12.7 | 12.7 |
| PLEKHA7 | Canis lupus familiaris OX=9615 | Pleckstrin homology domain containing A7 | 12.7 | 13.1 |
| LOC10215224( | Canis lupus familiaris OX=9615 | Uncharacterized protein | 12.7 | 13.1 |
| PLEK2 | Canis lupus familiaris OX=9615 | Pleckstrin 2 | 12.6 | 10.9 |
| RPS20 | Canis lupus familiaris OX=9615 | Ribosomal_S10 domain-containing protein | 12.6 | 12.6 |
| SYNJ1 | Canis lupus familiaris OX=9615 | Uncharacterized protein | 12.5 | 10.6 |
| BMP2K | Canis lupus familiaris OX=9615 | BMP2 inducible kinase | 12.5 | 11.5 |
| NEK6 | Canis lupus familiaris OX=9615 | NIMA related kinase 6 | 12.5 | 12.5 |
| SRGAP2 | Canis lupus familiaris OX=9615 | Uncharacterized protein | 12.5 | 19.3 |
| NRBF2 | Canis lupus familiaris OX=9615 | Nuclear receptor binding factor 2 | 12.5 | 27.4 |
| CLU | Canis lupus familiaris OX=9615 | Clusterin | 12.4 | 6.5 |
| NUFIP2 | Canis lupus familiaris OX=9615 | Nuclear FMR1 interacting protein 2 | 12.4 | 11.4 |
| RPS26 | Canis lupus familiaris OX=9615 | 40S ribosomal protein S26 | 12.4 | 12.4 |
| CLIC1 | Canis lupus familiaris OX=9615 | Chloride intracellular channel protein | 12.4 | 31.1 |
| PTPN12 | Canis lupus familiaris OX=9615 | Tyrosine-protein phosphatase non-receptor type 12 | 12.3 | 17 |
| PEBP1 | Canis lupus familiaris OX=9615 | Phosphatidylethanolamine-binding protein 1 | 12.3 | 18.7 |
| MAGI3 | Canis lupus familiaris OX=9615 | Membrane associated guanylate kinase, WW and PDZ domain containing 3 | 12.3 | 18.9 |
| CORO1C | Canis lupus familiaris OX=9615 | Coronin | 12.2 | 10.5 |
| CNN3 | Canis lupus familiaris OX=9615 | Calponin | 12.2 | 20.3 |
| VPS37C | Canis lupus familiaris OX=9615 | VPS37C subunit of ESCRT-I | 12.1 | 6.5 |
| LOC10085587 | Canis lupus familiaris OX=9615 | Uncharacterized protein | 12.1 | 7.2 |
| GTSE1 | Canis lupus familiaris OX=9615 | G2 and S-phase expressed 1 | 12.1 | 9.9 |
| SEPTIN6 | Canis lupus familiaris OX=9615 | Septin 6 | 12.1 | 18.2 |
| ARPC4 | Canis lupus familiaris OX=9615 | Actin-related protein 2/3 complex subunit 4 | 12 | 10.2 |
| CEP192 | Canis lupus familiaris OX=9615 | Centrosomal protein 192 | 12 | 11.1 |
| MOB1B | Canis lupus familiaris OX=9615 | MOB kinase activator 1B | 12 | 12 |
| GORASP2 | Canis lupus familiaris OX=9615 | Golgi reassembly stacking protein 2 | 12 | 12 |
| BLOC1S5 | Canis lupus familiaris OX=9615 | Biogenesis of lysosomal organelles complex 1 subunit 5 | 12 | 12 |
| SEC24B | Canis lupus familiaris OX=9615 | SEC24 homolog B, COPII coat complex component | 11.9 | 13.8 |
| RBM22 | Canis lupus familiaris OX=9615 | RNA binding motif protein 22 | 11.8 | 7.1 |
| TTK | Canis lupus familiaris OX=9615 | TTK protein kinase | 11.8 | 10.7 |
| ASAP1 | Canis lupus familiaris OX=9615 | ArfGAP with SH3 domain, ankyrin repeat and PH domain 1 | 11.8 | 11 |
| RPL31 | Canis lupus familiaris OX=9615 | Uncharacterized protein | 11.8 | 11.8 |
| CD44 | Canis lupus familiaris OX=9615 | CD44 antigen | 11.8 | 13.2 |
| MAP2K1 | Canis lupus familiaris OX=9615 | Mitogen-activated protein kinase kinase 1 | 11.8 | 14.7 |
| EIF5B | Canis lupus familiaris OX=9615 | Eukaryotic translation initiation factor 5B | 11.8 | 18.4 |
| MAP4K3 | Canis lupus familiaris OX=9615 | Mitogen-activated protein kinase kinase kinase kinase 3 | 11.7 | 8.5 |
| SH3D19 | Canis lupus familiaris OX=9615 | SH3 domain containing 19 | 11.7 | 11.4 |
| SNRPF | Canis lupus familiaris OX=9615 | Sm domain-containing protein | 11.7 | 11.7 |
| MPP1 | Canis lupus familiaris OX=9615 | Membrane palmitoylated protein 1 | 11.7 | 12.7 |
| SPDL1 | Canis lupus familiaris OX=9615 | Protein Spindly | 11.6 | 7.3 |
| SORBS3 | Canis lupus familiaris OX=9615 | Sorbin and SH3 domain containing 3 | 11.6 | 9.8 |
| STK38 | Canis lupus familiaris OX=9615 | Non-specific serine/threonine protein kinase | 11.6 | 11.6 |
| EPS8L2 | Canis lupus familiaris OX=9615 | EPS8 like 2 | 11.6 | 13 |
| NAP1L1 | Canis lupus familiaris OX=9615 | Nucleosome assembly protein 1 like 1 | 11.5 | 11.8 |
| RPS8 | Canis lupus familiaris OX=9615 | 40S ribosomal protein S8 | 11.5 | 12 |
| MTHFD1 | Canis lupus familiaris OX=9615 | Methylenetetrahydrofolate dehydrogenase, cyclohydrolase and formyltetrahydrofolate synthetase 1 | 11.5 | 12.6 |
| SLC9A1 | Canis lupus familiaris OX=9615 | Sodium/hydrogen exchanger | 11.4 | 8.1 |
| SFPQ | Canis lupus familiaris OX=9615 | Uncharacterized protein | 11.4 | 10.3 |
| EPCAM | Canis lupus familiaris OX=9615 | Epithelial cell adhesion molecule | 11.4 | 11.4 |
| RPL13 | Canis lupus familiaris OX=9615 | 60S ribosomal protein L13 | 11.4 | 16.1 |

| | | | | |
|---|---|---|---|---|
| GBP6 | Canis lupus familiaris OX=9615 | Guanylate binding protein family member 6 | 11.3 | 11 |
| COMMD3 | Canis lupus familiaris OX=9615 | COMM domain containing 3 | 11.3 | 11.3 |
| PPP1R9A | Canis lupus familiaris OX=9615 | Protein phosphatase 1 regulatory subunit 9A | 11.3 | 12 |
| STK25 | Canis lupus familiaris OX=9615 | Serine/threonine kinase 25 | 11.3 | 12.9 |
| YTHDF2 | Canis lupus familiaris OX=9615 | YTH N6-methyladenosine RNA binding protein 2 | 11.3 | 13.2 |
| ACBD3 | Canis lupus familiaris OX=9615 | Acyl-CoA binding domain containing 3 | 11.3 | 17.2 |
| SYTL4 | Canis lupus familiaris OX=9615 | Synaptotagmin like 4 | 11.2 | 7.6 |
| KRT78 | Canis lupus familiaris OX=9615 | Keratin 78 | 11.2 | 9.3 |
| ERC1 | Canis lupus familiaris OX=9615 | ELKS/RAB6-interacting/CAST family member 1 | 11.2 | 13.7 |
| OSMR | Canis lupus familiaris OX=9615 | Oncostatin M receptor | 11.2 | 14 |
| ANKRD50 | Canis lupus familiaris OX=9615 | Ankyrin repeat domain 50 | 11.1 | 5 |
| CNNM4 | Canis lupus familiaris OX=9615 | Cyclin and CBS domain divalent metal cation transport mediator 4 | 11.1 | 5.7 |
| CHCHD3 | Canis lupus familiaris OX=9615 | MIC | 11 | 11 |
| DGKA | Canis lupus familiaris OX=9615 | Diacylglycerol kinase | 11 | 14 |
| GSK3B | Canis lupus familiaris OX=9615 | Glycogen synthase kinase 3 beta | 11 | 14.5 |
| NUMBL | Canis lupus familiaris OX=9615 | NUMB like, endocytic adaptor protein | 10.9 | 7.4 |
| LOC10068361 | Canis lupus familiaris OX=9615 | S1 motif domain-containing protein | 10.9 | 10.9 |
| UBAP2 | Canis lupus familiaris OX=9615 | Ubiquitin associated protein 2 | 10.9 | 11.9 |
| PLK1 | Canis lupus familiaris OX=9615 | Serine/threonine-protein kinase PLK | 10.9 | 18.1 |
| GOLGA3 | Canis lupus familiaris OX=9615 | Golgin A3 | 10.8 | 16.5 |
| SRRM2 | Canis lupus familiaris OX=9615 | Serine/arginine repetitive matrix 2 | 10.7 | 5.2 |
| SLK | Canis lupus familiaris OX=9615 | STE20 like kinase | 10.7 | 14.5 |
| PRPF38B | Canis lupus familiaris OX=9615 | Pre-mRNA processing factor 38B | 10.6 | 7.7 |
| SNX11 | Canis lupus familiaris OX=9615 | Sorting nexin 11 | 10.6 | 10.6 |
| VDAC3 | Canis lupus familiaris OX=9615 | Voltage dependent anion channel 3 | 10.6 | 10.6 |
| HNRNPF | Canis lupus familiaris OX=9615 | Heterogeneous nuclear ribonucleoprotein F | 10.6 | 12.1 |
| BCAP31 | Canis lupus familiaris OX=9615 | B cell receptor associated protein 31 | 10.6 | 15 |
| STX6 | Canis lupus familiaris OX=9615 | Syntaxin 6 | 10.6 | 23.5 |
| ABCF1 | Canis lupus familiaris OX=9615 | ATP binding cassette subfamily F member 1 | 10.4 | 5.6 |
| MYO1E | Canis lupus familiaris OX=9615 | Myosin IE | 10.4 | 9.7 |
| PSMA5 | Canis lupus familiaris OX=9615 | Proteasome endopeptidase complex | 10.4 | 10.4 |
| TTF2 | Canis lupus familiaris OX=9615 | Transcription termination factor 2 | 10.4 | 11.9 |
| GPBP1L1 | Canis lupus familiaris OX=9615 | GC-rich promoter binding protein 1 like 1 | 10.4 | 12.9 |
| CNNM2 | Canis lupus familiaris OX=9615 | Cyclin and CBS domain divalent metal cation transport mediator 2 | 10.4 | 12.9 |
| SRCIN1 | Canis lupus familiaris OX=9615 | SRC kinase signaling inhibitor 1 | 10.4 | 16.7 |
| PARVA | Canis lupus familiaris OX=9615 | Parvin alpha | 10.4 | 17.8 |
| ZFYVE16 | Canis lupus familiaris OX=9615 | Zinc finger FYVE domain-containing protein | 10.3 | 8.2 |
| ATXN10 | Canis lupus familiaris OX=9615 | Ataxin 10 | 10.3 | 10.1 |
| CLIC4 | Canis lupus familiaris OX=9615 | Chloride intracellular channel protein | 10.3 | 10.3 |
| GGA3 | Canis lupus familiaris OX=9615 | Golgi associated, gamma adaptin ear containing, ARF binding protein 3 | 10.3 | 15.5 |
| ISG15 | Canis lupus familiaris OX=9615 | ISG15 ubiquitin like modifier | 10.3 | 22.4 |
| LGALS9 | Canis lupus familiaris OX=9615 | Galectin | 10.2 | 7.7 |
| TBC1D15 | Canis lupus familiaris OX=9615 | TBC1 domain family member 15 | 10.2 | 7.9 |
| RPL18A | Canis lupus familiaris OX=9615 | 60S ribosomal protein L18a | 10.2 | 10.2 |
| NOTCH2 | Canis lupus familiaris OX=9615 | Uncharacterized protein | 10.2 | 10.4 |
| PHLDB1 | Canis lupus familiaris OX=9615 | Pleckstrin homology like domain family B member 1 | 10.2 | 12.9 |
| RBM14 | Canis lupus familiaris OX=9615 | RNA binding motif protein 14 | 10.2 | 12.9 |
| HSPD1 | Canis lupus familiaris OX=9615 | 60 kDa heat shock protein, mitochondrial | 10.1 | 7.2 |
| NOXA1 | Canis lupus familiaris OX=9615 | NADPH oxidase activator 1 | 10.1 | 8.4 |
| ATP5F1B | Canis lupus familiaris OX=9615 | ATP synthase subunit beta | 10.1 | 10.1 |
| RBSN | Canis lupus familiaris OX=9615 | Uncharacterized protein | 10.1 | 13.3 |
| ASPSCR1 | Canis lupus familiaris OX=9615 | ASPSCR1 tether for SLC2A4, UBX domain containing | 10 | 6.2 |
| RASSF8 | Canis lupus familiaris OX=9615 | Ras association domain family member 8 | 10 | 10.5 |
| AGFG2 | Canis lupus familiaris OX=9615 | ArfGAP with FG repeats 2 | 10 | 14.2 |
| ACAT2 | Canis lupus familiaris OX=9615 | Acetyl-CoA acetyltransferase 2 | 9.9 | 9.9 |
| MGST3 | Canis lupus familiaris OX=9615 | Microsomal glutathione S-transferase 3 | 9.9 | 9.9 |
| NUP62 | Canis lupus familiaris OX=9615 | Nucleoporin 62 | 9.9 | 12.5 |
| SPECC1 | Canis lupus familiaris OX=9615 | Sperm antigen with calponin homology and coiled-coil domains 1 | 9.9 | 14.5 |
| PSD3 | Canis lupus familiaris OX=9615 | Pleckstrin and Sec7 domain containing 3 | 9.9 | 14.9 |
| FOXO3 | Canis lupus familiaris OX=9615 | Forkhead box O3 | 9.8 | 9.8 |
| TAB2 | Canis lupus familiaris OX=9615 | TGF-beta activated kinase 1 (MAP3K7) binding protein 2 | 9.8 | 11.7 |
| EPN1 | Canis lupus familiaris OX=9615 | Epsin 1 | 9.8 | 17.2 |
| PSMB10 | Canis lupus familiaris OX=9615 | Proteasome subunit beta | 9.8 | 20.4 |
| COMMD8 | Canis lupus familiaris OX=9615 | COMM domain containing 8 | 9.8 | 31.1 |
| RPL12 | Canis lupus familiaris OX=9615 | 60S ribosomal protein L12 | 9.7 | 9.7 |
| HAUS6 | Canis lupus familiaris OX=9615 | HAUS augmin like complex subunit 6 | 9.7 | 10.6 |
| HNRNPM | Canis lupus familiaris OX=9615 | Heterogeneous nuclear ribonucleoprotein M | 9.7 | 12.4 |
| SLC25A5 | Canis lupus familiaris OX=9615 | Solute carrier family 25 member 5 | 9.7 | 13.1 |
| BPNT1 | Canis lupus familiaris OX=9615 | 3'(2'), 5'-bisphosphate nucleotidase 1 | 9.7 | 18.1 |
| CCNB1 | Canis lupus familiaris OX=9615 | Cyclin N-terminal domain-containing protein | 9.6 | 6.6 |
| USP54 | Canis lupus familiaris OX=9615 | Ubiquitin specific peptidase 54 | 9.6 | 6.7 |
| GORAB | Canis lupus familiaris OX=9615 | Golgin, RAB6 interacting | 9.6 | 9.3 |
| RCAN1 | Canis lupus familiaris OX=9615 | Regulator of calcineurin 1 | 9.6 | 9.6 |
| SF3B6 | Canis lupus familiaris OX=9615 | Splicing factor 3b subunit 6 | 9.6 | 9.6 |
| TNS3 | Canis lupus familiaris OX=9615 | Tensin 3 | 9.6 | 19.6 |
| NMT1 | Canis lupus familiaris OX=9615 | Glycylpeptide N-tetradecanoyltransferase | 9.5 | 5.6 |

| | | | | |
|---|---|---|---|---|
| TMC5 | Canis lupus familiaris OX=9615 | Transmembrane channel-like protein | 9.5 | 8.6 |
| ZFYVE19 | Canis lupus familiaris OX=9615 | Zinc finger FYVE-type containing 19 | 9.5 | 9.5 |
| EIF3D | Canis lupus familiaris OX=9615 | Eukaryotic translation initiation factor 3 subunit D | 9.5 | 13.5 |
| SRP54 | Canis lupus familiaris OX=9615 | Signal recognition particle 54 kDa protein | 9.5 | 23.6 |
| ARHGAP23 | Canis lupus familiaris OX=9615 | Rho GTPase activating protein 23 | 9.4 | 9.5 |
| CMC1 | Canis lupus familiaris OX=9615 | COX assembly mitochondrial protein | 9.4 | 18.9 |
| ABCE1 | Canis lupus familiaris OX=9615 | ATP binding cassette subfamily E member 1 | 9.3 | 7.4 |
| ACOT7 | Canis lupus familiaris OX=9615 | Acyl-CoA thioesterase 7 | 9.3 | 8.4 |
| UBE2QL1 | Canis lupus familiaris OX=9615 | Ubiquitin conjugating enzyme E2 Q family like 1 | 9.3 | 9.3 |
| SLITRK4 | Canis lupus familiaris OX=9615 | SLIT and NTRK like family member 4 | 9.3 | 12.5 |
| ZC3HAV1 | Canis lupus familiaris OX=9615 | Zinc finger CCCH-type containing, antiviral 1 | 9.2 | 5.8 |
| CAPZB | Canis lupus familiaris OX=9615 | F-actin-capping protein subunit beta | 9.2 | 11.6 |
| SNX1 | Canis lupus familiaris OX=9615 | Sorting nexin 1 | 9.2 | 13.2 |
| CHMP3 | Canis lupus familiaris OX=9615 | Uncharacterized protein | 9.2 | 14.7 |
| FUBP3 | Canis lupus familiaris OX=9615 | Far upstream element binding protein 3 | 9.2 | 16.2 |
| GOSR1 | Canis lupus familiaris OX=9615 | Golgi SNAP receptor complex member 1 | 9.2 | 23.2 |
| NME1 | Canis lupus familiaris OX=9615 | Nucleoside diphosphate kinase | 9.2 | 23.3 |
| ARHGEF7 | Canis lupus familiaris OX=9615 | Rho guanine nucleotide exchange factor 7 | 9.1 | 9.7 |
| TDRP | Canis lupus familiaris OX=9615 | Testis development related protein | 9.1 | 15.6 |
| PSMD14 | Canis lupus familiaris OX=9615 | Proteasome 26S subunit, non-ATPase 14 | 9 | 10.4 |
| RSL1D1 | Canis lupus familiaris OX=9615 | Ribosomal L1 domain containing 1 | 9 | 11.6 |
| SASS6 | Canis lupus familiaris OX=9615 | SAS-6 centriolar assembly protein | 9 | 11.6 |
| ACTR3 | Canis lupus familiaris OX=9615 | Uncharacterized protein | 9 | 11.7 |
| TACC3 | Canis lupus familiaris OX=9615 | Transforming acidic coiled-coil containing protein 3 | 9 | 13.1 |
| ADAR | Canis lupus familiaris OX=9615 | Adenosine deaminase RNA specific | 8.9 | 7.5 |
| MVB12A | Canis lupus familiaris OX=9615 | Multivesicular body subunit 12A | 8.8 | 5.5 |
| SMG7 | Canis lupus familiaris OX=9615 | SMG7 nonsense mediated mRNA decay factor | 8.8 | 7.2 |
| UBXN6 | Canis lupus familiaris OX=9615 | UBX domain protein 6 | 8.8 | 17.9 |
| NAA50 | Canis lupus familiaris OX=9615 | N(alpha)-acetyltransferase 50, NatE catalytic subunit | 8.8 | 19 |
| PUF60 | Canis lupus familiaris OX=9615 | Poly(U) binding splicing factor 60 | 8.8 | 23.2 |
| HAL | Canis lupus familiaris OX=9615 | Histidine ammonia-lyase | 8.7 | 6.2 |
| U2SURP | Canis lupus familiaris OX=9615 | U2 snRNP associated SURP domain containing | 8.7 | 6.4 |
| EEF1G | Canis lupus familiaris OX=9615 | Uncharacterized protein | 8.7 | 12.2 |
| RASSF7 | Canis lupus familiaris OX=9615 | Ras association domain family member 7 | 8.6 | 6.9 |
| SKP1 | Canis lupus familiaris OX=9615 | S-phase kinase associated protein 1 | 8.6 | 8.6 |
| NR3C1 | Canis lupus familiaris OX=9615 | Glucocorticoid receptor | 8.6 | 11.7 |
| MTDH | Canis lupus familiaris OX=9615 | Metadherin | 8.6 | 13.7 |
| EIF5 | Canis lupus familiaris OX=9615 | Eukaryotic translation initiation factor 5 | 8.6 | 14.9 |
| HELLS | Canis lupus familiaris OX=9615 | Helicase, lymphoid specific | 8.5 | 8.3 |
| SH3PXD2A | Canis lupus familiaris OX=9615 | SH3 and PX domains 2A | 8.5 | 12 |
| CEP85 | Canis lupus familiaris OX=9615 | Centrosomal protein 85 | 8.5 | 15.8 |
| COPZ1 | Canis lupus familiaris OX=9615 | Coatomer subunit zeta | 8.5 | 15.8 |
| TAGLN | Canis lupus familiaris OX=9615 | Transgelin | 8.5 | 19.2 |
| ARFIP2 | Canis lupus familiaris OX=9615 | ADP ribosylation factor interacting protein 2 | 8.5 | 22.3 |
| PPP2R5D | Canis lupus familiaris OX=9615 | Serine/threonine-protein phosphatase 2A 56 kDa regulatory subunit | 8.4 | 6.6 |
| SYNJ2 | Canis lupus familiaris OX=9615 | Synaptojanin 2 | 8.4 | 11.4 |
| WIPI2 | Canis lupus familiaris OX=9615 | WD repeat domain, phosphoinositide interacting 2 | 8.3 | 5.3 |
| FAU | Canis lupus familiaris OX=9615 | 40S ribosomal protein S30 | 8.3 | 8.3 |
| UEVLD | Canis lupus familiaris OX=9615 | UEV and lactate/malate dehyrogenase domains | 8.3 | 9.3 |
| PPP1R12A | Canis lupus familiaris OX=9615 | Protein phosphatase 1 regulatory subunit | 8.3 | 10.9 |
| CCDC115 | Canis lupus familiaris OX=9615 | Coiled-coil domain containing 115 | 8.3 | 13.9 |
| NAA10 | Canis lupus familiaris OX=9615 | N(alpha)-acetyltransferase 10, NatA catalytic subunit | 8.3 | 14 |
| RACGAP1 | Canis lupus familiaris OX=9615 | Rac GTPase activating protein 1 | 8.2 | 6.3 |
| ENAH | Canis lupus familiaris OX=9615 | ENAH actin regulator | 8.2 | 10.3 |
| CRYBG1 | Canis lupus familiaris OX=9615 | Crystallin beta-gamma domain containing 1 | 8.1 | 8.8 |
| RSRC2 | Canis lupus familiaris OX=9615 | Arginine and serine rich coiled-coil 2 | 8 | 5.1 |
| EPS8L1 | Canis lupus familiaris OX=9615 | EPS8 like 1 | 8 | 5.6 |
| MYL3 | Canis lupus familiaris OX=9615 | Myosin light chain 3 | 8 | 8 |
| PIP5K1A | Canis lupus familiaris OX=9615 | PIPK domain-containing protein | 8 | 9.7 |
| HSPA5 | Canis lupus familiaris OX=9615 | Heat shock protein family A (Hsp70) member 5 | 8 | 10.2 |
| SUB1 | Canis lupus familiaris OX=9615 | PC4 domain-containing protein | 8 | 10.4 |
| FAM126A | Canis lupus familiaris OX=9615 | Family with sequence similarity 126 member A | 8 | 11.6 |
| MYO1C | Canis lupus familiaris OX=9615 | Myosin IC | 8 | 11.9 |
| RAC1 | Canis lupus familiaris OX=9615 | Ras-related C3 botulinum toxin substrate 1 | 7.9 | 7.9 |
| PAWR | Canis lupus familiaris OX=9615 | Uncharacterized protein | 7.8 | 7.8 |
| MBNL1 | Canis lupus familiaris OX=9615 | Muscleblind like splicing regulator 1 | 7.8 | 12.7 |
| FLVCR1 | Canis lupus familiaris OX=9615 | MFS domain-containing protein | 7.7 | 7.7 |
| EML2 | Canis lupus familiaris OX=9615 | EMAP like 2 | 7.7 | 9.4 |
| LOC10085582 | Canis lupus familiaris OX=9615 | DUF3338 domain-containing protein | 7.7 | 11.1 |
| LRRC59 | Canis lupus familiaris OX=9615 | Leucine rich repeat containing 59 | 7.7 | 14.2 |
| KAZN | Canis lupus familiaris OX=9615 | Kazrin, periplakin interacting protein | 7.7 | 17.9 |
| ILKAP | Canis lupus familiaris OX=9615 | ILK associated serine/threonine phosphatase | 7.6 | 9 |
| TMED10 | Canis lupus familiaris OX=9615 | Transmembrane p24 trafficking protein 10 | 7.6 | 12.5 |
| TMF1 | Canis lupus familiaris OX=9615 | TATA element modulatory factor 1 | 7.5 | 7.7 |
| ATP1A1 | Canis lupus familiaris OX=9615 | Sodium/potassium-transporting ATPase subunit alpha | 7.5 | 8.8 |
| FAM126B | Canis lupus familiaris OX=9615 | Family with sequence similarity 126 member B | 7.5 | 9.4 |

| | | | | |
|---|---|---|---|---|
| MARCKSL1 | Canis lupus familiaris OX=9615 | MARCKS like 1 | 7.5 | 18.1 |
| WDR1 | Canis lupus familiaris OX=9615 | WD repeat domain 1 | 7.4 | 5.9 |
| RBM47 | Canis lupus familiaris OX=9615 | RNA-binding protein 47 | 7.4 | 7.4 |
| EIF6 | Canis lupus familiaris OX=9615 | Eukaryotic translation initiation factor 6 | 7.4 | 7.4 |
| UGP2 | Canis lupus familiaris OX=9615 | UTP--glucose-1-phosphate uridylyltransferase | 7.4 | 7.4 |
| ZDHHC3 | Canis lupus familiaris OX=9615 | Palmitoyltransferase | 7.4 | 7.4 |
| FAM171B | Canis lupus familiaris OX=9615 | Family with sequence similarity 171 member B | 7.4 | 10.8 |
| PPP1R9B | Canis lupus familiaris OX=9615 | Protein phosphatase 1 regulatory subunit 9B | 7.3 | 7.3 |
| GPATCH1 | Canis lupus familiaris OX=9615 | G-patch domain containing 1 | 7.3 | 11.4 |
| RALA | Canis lupus familiaris OX=9615 | RAS like proto-oncogene A | 7.3 | 13.1 |
| KPNA3 | Canis lupus familiaris OX=9615 | Importin subunit alpha | 7.2 | 7.2 |
| NAP1L4 | Canis lupus familiaris OX=9615 | Nucleosome assembly protein 1 like 4 | 7.2 | 7.5 |
| STRN3 | Canis lupus familiaris OX=9615 | Striatin 3 | 7.2 | 9.8 |
| PLEKHA1 | Canis lupus familiaris OX=9615 | Pleckstrin homology domain containing A1 | 7.2 | 9.9 |
| TANC1 | Canis lupus familiaris OX=9615 | Tetratricopeptide repeat, ankyrin repeat and coiled-coil containing 1 | 7.2 | 10.5 |
| CCDC88C | Canis lupus familiaris OX=9615 | Coiled-coil domain containing 88C | 7.2 | 11.4 |
| PKN3 | Canis lupus familiaris OX=9615 | Protein kinase N3 | 7.2 | 11.5 |
| GINS4 | Canis lupus familiaris OX=9615 | DNA replication complex GINS protein SLD5 | 7.2 | 13.5 |
| CAMLG | Canis lupus familiaris OX=9615 | Calcium signal-modulating cyclophilin ligand | 7.1 | 10.1 |
| VDAC1 | Canis lupus familiaris OX=9615 | Voltage dependent anion channel 1 | 7.1 | 12 |
| CC2D1A | Canis lupus familiaris OX=9615 | Coiled-coil and C2 domain containing 1A | 7 | 6.6 |
| MAPK1IP1L | Canis lupus familiaris OX=9615 | Mitogen-activated protein kinase 1 interacting protein 1 like | 7 | 7 |
| GPRC5C | Canis lupus familiaris OX=9615 | G protein-coupled receptor class C group 5 member C | 7 | 10.2 |
| SUGP1 | Canis lupus familiaris OX=9615 | SURP and G-patch domain containing 1 | 6.9 | 5 |
| ZFP36L2 | Canis lupus familiaris OX=9615 | ZFP36 ring finger protein like 2 | 6.9 | 6.9 |
| CDK14 | Canis lupus familiaris OX=9615 | Cyclin dependent kinase 14 | 6.9 | 7.6 |
| C5H16orf70 | Canis lupus familiaris OX=9615 | Chromosome 5 C16orf70 homolog | 6.9 | 8.3 |
| IFNGR1 | Canis lupus familiaris OX=9615 | Interferon gamma receptor 1 | 6.9 | 9.5 |
| RPL4 | Canis lupus familiaris OX=9615 | 60S ribosomal protein L4 | 6.8 | 5.3 |
| PLSCR3 | Canis lupus familiaris OX=9615 | Phospholipid scramblase | 6.8 | 6.8 |
| ARHGAP39 | Canis lupus familiaris OX=9615 | Rho GTPase activating protein 39 | 6.7 | 6.6 |
| ZC2HC1A | Canis lupus familiaris OX=9615 | Zinc finger C2HC-type containing 1A | 6.7 | 6.7 |
| DLA | Canis lupus familiaris OX=9615 | MHC class II DR alpha chain | 6.7 | 6.7 |
| PABPC1 | Canis lupus familiaris OX=9615 | Polyadenylate-binding protein | 6.7 | 10.8 |
| ACTN4 | Canis lupus familiaris OX=9615 | Actinin alpha 4 | 6.7 | 10.9 |
| PKN1 | Canis lupus familiaris OX=9615 | Protein kinase N1 | 6.6 | 5.2 |
| ILK | Canis lupus familiaris OX=9615 | Integrin linked kinase | 6.6 | 6.6 |
| KLK7 | Canis lupus familiaris OX=9615 | Kallikrein D | 6.6 | 6.6 |
| PEA15 | Canis lupus familiaris OX=9615 | Proliferation and apoptosis adaptor protein 15 | 6.6 | 7.9 |
| IDH2 | Canis lupus familiaris OX=9615 | Isocitrate dehydrogenase [NADP] | 6.6 | 8.3 |
| ALKBH3 | Canis lupus familiaris OX=9615 | AlkB homolog 3, alpha-ketoglutaratedependent dioxygenase | 6.6 | 10 |
| TBC1D10A | Canis lupus familiaris OX=9615 | TBC1 domain family member 10A | 6.6 | 11.4 |
| RHOJ | Canis lupus familiaris OX=9615 | Ras homolog family member J | 6.5 | 6.5 |
| RAB5B | Canis lupus familiaris OX=9615 | RAB5B, member RAS oncogene family | 6.5 | 12.1 |
| USP53 | Canis lupus familiaris OX=9615 | Ubiquitin specific peptidase 53 | 6.5 | 12.4 |
| MYO1D | Canis lupus familiaris OX=9615 | Unconventional myosin-Id | 6.5 | 14.2 |
| DLGAP5 | Canis lupus familiaris OX=9615 | DLG associated protein 5 | 6.4 | 5.3 |
| LUC7L2 | Canis lupus familiaris OX=9615 | Uncharacterized protein | 6.4 | 5.9 |
| SSR4 | Canis lupus familiaris OX=9615 | Signal sequence receptor subunit 4 | 6.4 | 6.4 |
| STON2 | Canis lupus familiaris OX=9615 | Stonin-2 | 6.4 | 10.6 |
| TBCC | Canis lupus familiaris OX=9615 | Tubulin folding cofactor C | 6.4 | 10.7 |
| TACC2 | Canis lupus familiaris OX=9615 | Transforming acidic coiled-coil containing protein 2 | 6.3 | 5.1 |
| NUP50 | Canis lupus familiaris OX=9615 | Nucleoporin 50 | 6.3 | 6.3 |
| FN3KRP | Canis lupus familiaris OX=9615 | Fructosamine 3 kinase related protein | 6.3 | 6.3 |
| KANK1 | Canis lupus familiaris OX=9615 | KN motif and ankyrin repeat domains 1 | 6.3 | 6.4 |
| HADHA | Canis lupus familiaris OX=9615 | Hydroxyacyl-CoA dehydrogenase trifunctional multienzyme complex subunit alpha | 6.3 | 6.7 |
| HMGCS1 | Canis lupus familiaris OX=9615 | 3-hydroxy-3-methylglutaryl coenzyme A synthase | 6.3 | 7.7 |
| PTPN13 | Canis lupus familiaris OX=9615 | Protein tyrosine phosphatase, non-receptor type 13 | 6.3 | 9.8 |
| EXOC4 | Canis lupus familiaris OX=9615 | Exocyst complex component 4 | 6.3 | 10.9 |
| AP3S1 | Canis lupus familiaris OX=9615 | AP complex subunit sigma | 6.3 | 21.2 |
| CCDC92 | Canis lupus familiaris OX=9615 | Coiled-coil domain containing 92 | 6.3 | 24.7 |
| CELF1 | Canis lupus familiaris OX=9615 | CUGBP Elav-like family member 1 | 6.2 | 6.2 |
| SNF8 | Canis lupus familiaris OX=9615 | Vacuolar-sorting protein SNF8 | 6.2 | 9.3 |
| ERRFI1 | Canis lupus familiaris OX=9615 | ERBB receptor feedback inhibitor 1 | 6.2 | 10.8 |
| SEC23IP | Canis lupus familiaris OX=9615 | SEC23 interacting protein | 6.2 | 10.9 |
| CKAP4 | Canis lupus familiaris OX=9615 | Cytoskeleton associated protein 4 | 6.2 | 11.7 |
| MCCC1 | Canis lupus familiaris OX=9615 | Methylcrotonoyl-CoA carboxylase 1 | 6.2 | 14.9 |
| VIPAS39 | Canis lupus familiaris OX=9615 | VPS33B interacting protein, apical-basolateral polarity regulator, spe-39 homolog | 6.1 | 11.5 |
| RAB10 | Canis lupus familiaris OX=9615 | Ras-related protein Rab-10 | 6 | 6 |
| CAVIN2 | Canis lupus familiaris OX=9615 | Caveolae associated protein 2 | 6 | 6 |
| SH3RF1 | Canis lupus familiaris OX=9615 | SH3 domain containing ring finger 1 | 6 | 8.5 |
| ANXA8L1 | Canis lupus familiaris OX=9615 | Annexin | 6 | 10.1 |
| PRXL2A | Canis lupus familiaris OX=9615 | Peroxiredoxin like 2A | 6 | 11.6 |
| SNAP47 | Canis lupus familiaris OX=9615 | Uncharacterized protein | 6 | 11.8 |
| PGK1 | Canis lupus familiaris OX=9615 | Phosphoglycerate kinase | 6 | 16.5 |
| CNOT4 | Canis lupus familiaris OX=9615 | CCR4-NOT transcription complex subunit 4 | 5.9 | 5.9 |

| | | | | |
|---|---|---|---|---|
| GIGYF2 | Canis lupus familiaris OX=9615 | GRB10 interacting GYF protein 2 | 5.9 | 5.9 |
| AKAP13 | Canis lupus familiaris OX=9615 | A-kinase anchoring protein 13 | 5.9 | 7 |
| C2CD2 | Canis lupus familiaris OX=9615 | C2 calcium dependent domain containing 2 | 5.8 | 5.8 |
| PTPN23 | Canis lupus familiaris OX=9615 | Protein tyrosine phosphatase, non-receptor type 23 | 5.8 | 6.4 |
| CLTC | Canis lupus familiaris OX=9615 | Clathrin heavy chain | 5.8 | 6.6 |
| GPSM2 | Canis lupus familiaris OX=9615 | G protein signaling modulator 2 | 5.8 | 7.4 |
| WDR44 | Canis lupus familiaris OX=9615 | WD repeat domain 44 | 5.8 | 7.6 |
| TTC7A | Canis lupus familiaris OX=9615 | Tetratricopeptide repeat domain 7A | 5.8 | 10.3 |
| TRIOBP | Canis lupus familiaris OX=9615 | TRIO and F-actin binding protein | 5.8 | 11.6 |
| LMNB1 | Canis lupus familiaris OX=9615 | Lamin B1 | 5.7 | 8.6 |
| TPD52L2 | Canis lupus familiaris OX=9615 | TPD52 like 2 | 5.7 | 16.6 |
| PTBP1 | Canis lupus familiaris OX=9615 | Polypyrimidine tract binding protein 1 | 5.6 | 5.2 |
| ATE1 | Canis lupus familiaris OX=9615 | Arginyl-tRNA--protein transferase 1 | 5.6 | 5.4 |
| STX11 | Canis lupus familiaris OX=9615 | Syntaxin 11 | 5.6 | 6.3 |
| CLIP2 | Canis lupus familiaris OX=9615 | CAP-Gly domain containing linker protein 2 | 5.6 | 6.9 |
| DLG5 | Canis lupus familiaris OX=9615 | Discs large MAGUK scaffold protein 5 | 5.6 | 8.9 |
| ABCC5 | Canis lupus familiaris OX=9615 | ATP binding cassette subfamily C member 5 | 5.6 | 9.3 |
| TSG101 | Canis lupus familiaris OX=9615 | Tumor susceptibility 101 | 5.6 | 11 |
| UBAP1 | Canis lupus familiaris OX=9615 | Ubiquitin associated protein 1 | 5.6 | 12.3 |
| CORO7 | Canis lupus familiaris OX=9615 | Coronin | 5.5 | 5.5 |
| ZW10 | Canis lupus familiaris OX=9615 | Zw10 kinetochore protein | 5.5 | 7.4 |
| PTPN6 | Canis lupus familiaris OX=9615 | Tyrosine-protein phosphatase non-receptor type | 5.5 | 9.7 |
| CENPH | Canis lupus familiaris OX=9615 | Centromere protein H | 5.5 | 11.1 |
| RPSA | Canis lupus familiaris OX=9615 | 40S ribosomal protein SA | 5.4 | 5.4 |
| VPS8 | Canis lupus familiaris OX=9615 | VPS8 subunit of CORVET complex | 5.4 | 5.5 |
| RAB11FIP5 | Canis lupus familiaris OX=9615 | RAB11 family interacting protein 5 | 5.4 | 5.5 |
| RAE1 | Canis lupus familiaris OX=9615 | Ribonucleic acid export 1 | 5.4 | 11.4 |
| SND1 | Canis lupus familiaris OX=9615 | Staphylococcal nuclease domain-containing protein | 5.3 | 6.8 |
| NUDT21 | Canis lupus familiaris OX=9615 | Cleavage and polyadenylation specificity factor subunit 5 | 5.3 | 9.3 |
| BMPR2 | Canis lupus familiaris OX=9615 | Receptor protein serine/threonine kinase | 5.3 | 10.4 |
| RICTOR | Canis lupus familiaris OX=9615 | RPTOR independent companion of MTOR complex 2 | 5.2 | 5.1 |
| ZWINT | Canis lupus familiaris OX=9615 | ZW10 interacting kinetochore protein | 5.2 | 5.2 |
| PSME1 | Canis lupus familiaris OX=9615 | Proteasome activator subunit 1 | 5.2 | 5.2 |
| AP3D1 | Canis lupus familiaris OX=9615 | Adaptor related protein complex 3 subunit delta 1 | 5.2 | 6.3 |
| XRN1 | Canis lupus familiaris OX=9615 | 5'-3' exoribonuclease 1 | 5.2 | 8.2 |
| LYSMD4 | Canis lupus familiaris OX=9615 | LysM domain containing 4 | 5.2 | 17.9 |
| TWF2 | Canis lupus familiaris OX=9615 | Toll-like receptor 9 | 5.2 | 28.9 |
| PSMD2 | Canis lupus familiaris OX=9615 | 26S proteasome non-ATPase regulatory subunit 2 | 5.1 | 5.1 |
| CARD6 | Canis lupus familiaris OX=9615 | Caspase recruitment domain family member 6 | 5.1 | 5.6 |
| IFRD1 | Canis lupus familiaris OX=9615 | Interferon related developmental regulator 1 | 5.1 | 8.9 |
